# Dopamine and schizophrenia from bench to bedside: Discovery of a striatal co-expression risk gene set that predicts in vivo measures of striatal function

**DOI:** 10.1101/2023.09.20.558594

**Authors:** Leonardo Sportelli, Daniel P. Eisenberg, Roberta Passiatore, Enrico D’Ambrosio, Linda A. Antonucci, Qiang Chen, Jasmine Czarapata, Aaron L. Goldman, Michael Gregory, Kira Griffiths, Thomas M. Hyde, Joel E. Kleinman, Antonio F. Pardiñas, Madhur Parihar, Teresa Popolizio, Antonio Rampino, Joo Heon Shin, Mattia Veronese, William S. Ulrich, Caroline F. Zink, Alessandro Bertolino, Oliver D. Howes, Karen F. Berman, Daniel R. Weinberger, Giulio Pergola

## Abstract

Schizophrenia (SCZ) is characterized by a polygenic risk architecture implicating diverse molecular pathways important for synaptic function. However, how polygenic risk funnels through these pathways to translate into syndromic illness is unanswered. To evaluate biologically meaningful pathways of risk, we used tensor decomposition to characterize gene co-expression in post-mortem brain (of neurotypicals: N=154; patients with SCZ: N=84; and GTEX samples N=120) from caudate nucleus (CN), hippocampus (HP), and dorsolateral prefrontal cortex (DLPFC). We identified a CN-predominant gene set showing dopaminergic selectivity that was enriched for genes associated with clinical state and for genes associated with SCZ risk. Parsing polygenic risk score for SCZ based on this specific gene set (parsed-PRS), we found that greater pathway-specific SCZ risk predicted greater *in vivo* striatal dopamine synthesis capacity measured by [^18^F]-FDOPA PET in three independent cohorts of neurotypicals and patients (total N=235) and greater fMRI striatal activation during reward anticipation in two additional independent neurotypical cohorts (total N=141). These results reveal a ‘bench to bedside’ translation of dopamine-linked genetic risk variation in driving *in vivo* striatal neurochemical and hemodynamic phenotypes that have long been implicated in the pathophysiology of SCZ.

## Introduction

Schizophrenia (SCZ) is a mental illness with complex heritability and polygenic architecture^1^. The largest genome-wide association study (GWAS) to date has identified an extensive set of potential SCZ risk genes converging on synaptic biology of central nervous system neurons^2^. To the extent that the downstream consequences of diverse risk alleles might affect shared biological functions, genetic risk for SCZ is likely best understood in the context of molecular ensembles, rather than at a single gene level. This perspective puts gene co-expression at the forefront of investigating genetic risk convergence as an instrumental approach to model the effect of many variants on interconnected genetic systems and, ultimately, downstream neurochemical and neural functioning^3–5^.

There is a large body of evidence which implicates synaptic dysfunction and neurotransmission across several key brain circuits which bridge the striatum, the dorsolateral prefrontal cortex (DLPFC) and the hippocampus (HP) as key pathological mechanisms in SCZ^6–9^. As such, understanding gene co-expression across multiple brain regions may increase understanding of how broad genetic variation translates into increased risk of illness^10^.

There is also a large body of evidence for dopamine involvement in SCZ, including emergence of psychotic symptoms (e.g., hallucinations and delusions) following administration of pro-dopaminergic agents and therapeutic antipsychotic effects elicited by dopamine blocking drugs targeting D_2_ receptors^11^. In the D_2_-rich striatum where illness-related dysfunction has been observed, positron emission tomography (PET) studies have found an array of dopamine system disturbances in SCZ suggesting increased dopaminergic drive from mesencephalic synaptic terminals, including elevated presynaptic dopamine synthesis ^7,12–15^. There is also evidence that individuals at clinical risk for SCZ, e.g., with subthreshold psychotic symptoms, as well as first degree relatives show a similar pattern of elevated striatal presynaptic dopamine synthesis capacity^16,17^, which may be enhanced with progression to frank illness^18^. Importantly, striatal dopamine synthesis shows heterogeneity across patients with SCZ^19^, particularly in treatment-resistant individuals, who have demonstrated synthesis capacity decreases^20^ and in whom different mechanisms may be at play^21^. Recent evidence from *post-mortem* human caudate has revealed that decreased expression of the short (predominantly autoreceptor) isoform of the D_2_ dopamine receptor gene *DRD2* – and not the long (predominantly postsynaptic) isoform – may be the causative mechanism for association of the SCZ GWAS risk allele mapped to the *DRD2* locus^20^. By identifying diminished expression of the inhibitory D_2_ presynaptic autoreceptor as one potential mechanism of SCZ risk, this work further implicates exaggerated presynaptic dopamine activity in pathogenesis^22^, consistent with earlier work associating a single-nucleotide polymorphism (SNP) with differential *DRD2* splicing, striatal dopamine D2 signaling, and prefrontal and striatal activity during working memory^23,24^.

Functional magnetic resonance imaging (fMRI) studies have reported altered brain activity in patients with SCZ while performing dopamine-dependent reward processing tasks, possibly arising from synaptic dysfunction and neurotransmission dysregulation^25,26^. Moreover, anticipatory striatal activation during reward task performance has been shown to be a heritable trait^27^ (*h^2^*= .20-.73) suggesting that genetic investigations may help better define important connections between this phenotype and dopamine-relevant SCZ risk molecular factors. In light of this and because dopamine dysfunction in SCZ generally appears to have at least in part a genetic basis^22,28–30^, we hypothesized that a SCZ-related genetically driven increase of striatal presynaptic dopamine synthesis might be reflected functionally in an increase of striatal fMRI activation during reward anticipation at least in neurotypical individuals.

While most of the same genes are expressed across brain regions, mRNA expression patterns vary consistently with differing functions subserved at a system-level. A widely used approach to analyze gene co-expression patterns is a combination of graph theory and clustering ^3,31–33^, such as in the popular weighted gene co-expression network analysis^34^. This approach, however, has important limitations, in its handling of higher-dimensional data, particularly in accounting for the multiplicity of co-expression contexts across brain regions and cell types, crucial aspects to capture the biological reality in which different tissues, cells and molecular pathways share common genes. Another class of co-expression detection methods called sparse decomposition of arrays (SDA) circumvents these limitations ^35^. SDA is based on singular value decomposition, a family of techniques that includes independent component analysis (ICA) and principal component analysis (PCA) and is able to effectively identify relationships between genes in multi-tissue experiments^35^. SDA decomposes a 3D Array (also called a “Tensor”) with dimensions representing individuals, genes and tissues, respectively, into several latent components (or factors) that represent major directions of variation in the data set. This approach identifies components that uncover functional biology^35,36^, and outperforms other co-expression detection strategies in the identification of functionally related and co-regulated groups of genes^37^.

Using SDA and two independent post-mortem brain samples, we investigated human RNA sequencing data from three brain regions prominently implicated in SCZ, i.e., CN, HP, and DLPFC (Fig. 1). We sought to identify gene sets enriched both for genes differentially expressed in SCZ and for genes associated with SCZ genetic risk. Focusing on genes sets with convergence of illness state and illness risk in neurotypical brain avoids epiphenomena related to drug treatment in patient samples and the same directionality of effects supports genetic risk inferences.

**Fig. 1:**
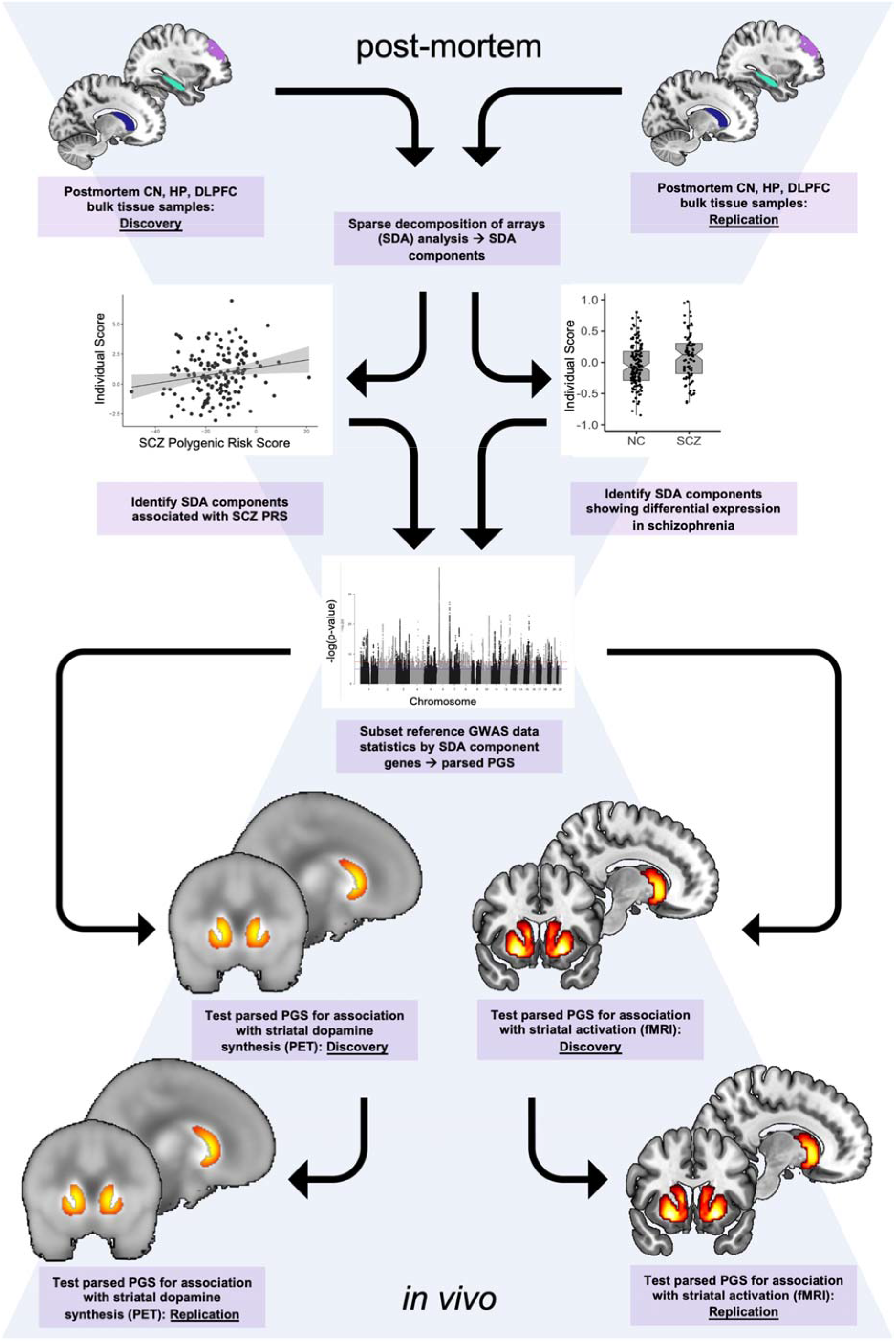
Graphic summary of the study design.

We identified a co-expression component in the SDA data that meets these criteria and is especially enriched for DA function genes. We then aimed to evaluate whether this component specifically translates into SCZ-relevant brain functional correlates in vivo. To that end, we studied striatal dopamine synthesis capacity determined via PET in both neurotypical controls (NC) and patients with SCZ and obtained corroborative evidence in an independent replication dataset. We then measured brain physiological activation during reward anticipation measured with functional magnetic resonance imaging (fMRI) in two independent neurotypical cohorts performing different reward tasks. We sought to translate dopamine-linked gene sets in postmortem brain involved in manifest illness and in illness risk into neurochemical and neurofunctional outcomes in the living human brain concordant with known SCZ associated phenotypes.

## Results

### Gene co-expression analysis

From SDA of postmortem CN, HP, and DLPFC tissue from our discovery cohort (Table 1) we obtained 69 robust components not associated with confounding variables. Supplementary Data 1 and 2 report output of SDA as well as the association with biological covariates and technical confounders and summary information regarding the number of genes included in each component.

**Table 1.** Demographics. Demographics of cohorts used in the gene co-expression (upper rows) and neuroimaging (lower rows) analyses are tabulated. Neuroimaging samples are those with both genetic and imaging data after ancestry stratification. *Abbreviations: CN: Caudate Nucleus bulk tissue data; HP: hippocampus bulk tissue data; DLPFC: dorsolateral prefrontal cortex bulk tissue data; NC: Neurotypical controls; SCZ: Patients with schizophrenia; AA: African American; EUR: European*

| Modality | Cohort | Diagnosis (NC/SCZ) | N | Age (years±SD) | Sex (N female) | Ethnicity (EUR/AA) | Genes |
| --- | --- | --- | --- | --- | --- | --- | --- |
| Postmortem data (CN, HP, DLPFC) | Discovery | NC | 154 | 44.9 ± 16.9 | 46 | 70/84 | 22,356 |
| | | SCZ | 84 | 49.4 ± 16.5 | 24 | 39/45 | |
|  | Replication | NC | 120 | range: 20 - 79* | 30 | 120/0 | 20,475 |
| <sup>18</sup>F-FDOPA PET | Discovery | NC | 65 | 29±9 | 31 | 65/0 | - |
| | | SCZ | 20 | 29±8 | 3 | 20/0 | - |
| | Replication | NC | 150 | 35±11 | 75 | 150/0 | - |
| Reward fMRI | Discovery | NC | 86 | 32±6 | 47 | 86/0 | - |
| | Replication | NC | 55 | 26±6 | 33 | 55/0 | - |

\* For the postmortem replication (GTEx) dataset, age information was only available as a discrete variable and range is reported the age range of the cohort

When comparing samples from neurotypical controls (NC) and individuals with SCZ, two of 69 filtered components (C80: 2497 genes; C109: 1211 genes; see Supplementary Data 2 for component gene membership) were associated with diagnosis (C80: F_[1,210]_=11.4, p=.0009, p_[FDR]_=.038; C109: F_[1,210]_=10.9, p=.001, p_[FDR]_=.039) (Fig. 2a). To identify SCZ-associated components more likely linked to pathogenic biology rather than treatment history or other factors, we additionally tested these components across samples for association with SCZ genetic risk before proceeding with further analyses. Only the SDA component C80 was also significantly associated with SCZ polygenic risk score (PRS), a measure of overall cumulative risk burden, in a diagnosis-consistent direction (see *Online methods* for PRS computation; C80: t_[93]_=1.67, one-tailed p=.048; C109: t_[93]_=-1.2, one-tailed p=.11) (Fig. 2a). Patients with SCZ had greater C80 scores and, consistently, healthy controls with greater SCZ PRS had relatively greater C80 scores.

**Fig. 2:**
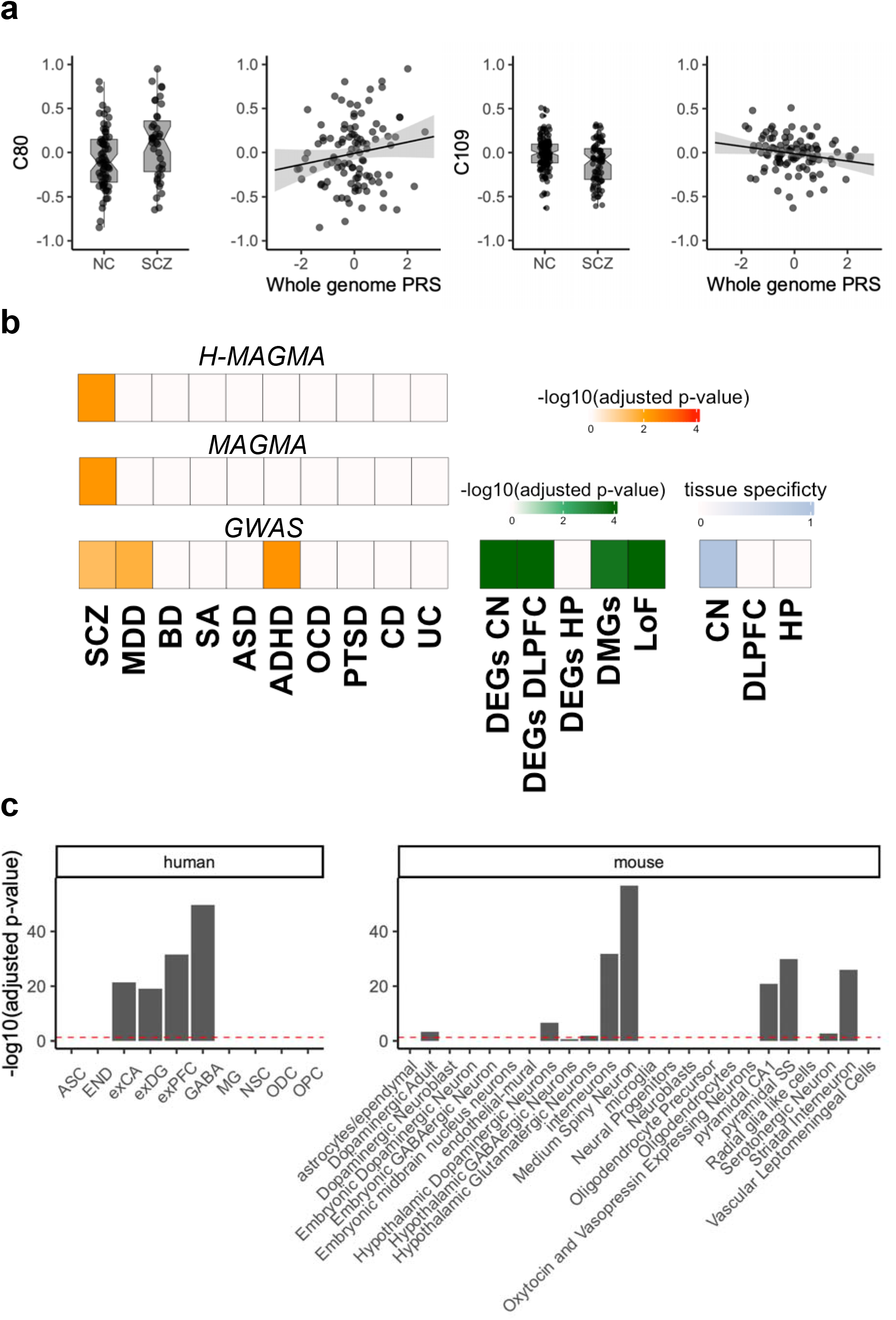
Sparse decomposition of arrays (SDA) component characterization. **a**, Notched box plots show SDA component C80 and C109 scores for post-mortem data samples in SCZ and NC groups. These were the only component showing a significant group effect. Group medians (horizontal line), 95% confidence intervals (notches), interquartile range (box edges), and whiskers (25^th^/75^th^ percentiles or extrema) are shown. Scatter plot demonstrates SDA component C80 and C109 scores as a function of polygenic risk for schizophrenia and includes regression fit line with shaded 95% confidence interval. C80 is the only one with a significant PRS association consistent with diagnosis direction. **b**, Gene enrichment analysis results are shown for C80 component. From the bottom, the first (GWAS), second (MAGMA) and third orange grids (H-MAGMA) show enrichment results for schizophrenia risk genes, other psychiatric illness risk genes, and immune condition risk genes. Enrichment testing results are shown for differentially expressed genes, differentially methylated genes, and loss of function variant intolerant genes in the green grid. The final lightblue grid show C80 tissue specificity as determined by the tissue scores generated during the SDA process and reflects the relative contribution of component gene networks within each of the sampled regions to the overall component. Adjusted p-values shown are empirical p-values obtained from permutation tests. **c**, Cell-type specificity of C80 component using human (left) and mouse (right) single-cell atlases. y-axes show FDR-adjusted p-values. Red dashed lines represent p-value=.05. Barplots demonstrates a higher specicifty for GABAergic, medium spiny and dopaminergic neurons. *Abbreviations: ADHD: attention deficit hyperactivity disorder; ASC: astrocytes; ASD: autism spectrum disorder; BD: bipolar disorder; CD: Crohn’s disease; CN: Caudate Nucleus; DEGs: differentially expressed genes; DLPFC: dorsolateral prefrontal cortex; DMGs: differentially methylated genes*; *END: endothelial cells; HP: hippocampus; exCA: pyramidal neurons from the hippocampal CA region; exDG: granule neurons from the hippocampal dentate gyrus; exPFC: pyramidal neurons from the prefrontal cortex; GABA: GABAergic interneurons; LoF: loss of function intolerant genes; MDD: major depressive disorder; MG: microglia; NC: Neurotypical controls; NSC: neuronal stem cells; OCD: obsessive compulsive disorder; ODC: oligodendrocytes; OPC: oligodendrocyte precursor cells; PRS: polygenic risk score as reported by the third wave (primary) analyses of the Psychiatric Genetics Consortium*^2^*; PTSD: posttraumatic stress disorder; SA: suicide attempt; SCZ: Patients with schizophrenia and UC: ulcerative colitis*.

Biological characterization of this component showed enrichment for SCZ, major depressive disorder (MDD) and attention deficit hyperactivity disorder (ADHD) risk genes, SCZ differentially expressed genes (DEGs) previously observed in the CN^22^ and in the DLPFC, and differentially methylated genes (DMG; i.e., genes proximal to regions enriched in CpG islands differentially methylated in SCZ compared to healthy controls) and also loss of function intolerant genes (all empirical p <.05; Fig. 2b). Moreover, we used Multi-marker Analysis of GenoMic Annotation (MAGMA)^38^ and H-MAGMA^39^, which leverages chromatin accessibility datasets, to perform a gene-set enrichment analysis for pathology-specific GWAS variants and found that the association with SCZ risk of the variants falling within or regulating (based on chromatin interactions) C80 genes is greater than that in the remaining sets (p_[FDR]_ < .05; Fig. 2b). Interestingly, this technique shows greater specificity to SCZ, as there was no consistency across MAGMA, H-MAGMA, and GWAS variant analyses with MDD and ADHD results (Fig. 2b). See Supplementary Fig. 1a for results on biological characterization of all 69 robust components.

To determine which tissue contributed more to the inter-individual variation within a given component, we checked the tissue score matrix obtained by SDA, which represents the covariance between the overall gene expression derived from one tissue and the component identified. Using a threshold of |0.5| (as previously reported by SDA developers^35^) in the tissue loading matrix, we found the C80 component to be most active in the CN (Fig. 2b). Accordingly, cell specificity analysis suggested a highly significant preponderance of medium spiny neurons (MSNs) and dopaminergic terminals (p_[FDR]_=1.9×10^-57^; Fig. 2c), consistent with CN localization. Gene ontology analysis (Fig. 3a, Supplementary Fig. 1b-c and Supplementary Data 3) characterized C80 as a predominantly synaptic component (133 genes, fold enrichment=2.3, p_[FDR]_=1×10^-18^) with both pre (128 genes, fold enrichment=1.9, p_[FDR]_=2.2×10^-12^) and postsynaptic specializations (103 genes, fold enrichment=2, p_[FDR]_=8×10^-12^, Fig. 3a). KEGG pathway analysis showed enrichment for dopaminergic, GABAergic, glutamatergic, and cholinergic synapses, all characteristic of the CN (Fig. 3b and Supplementary Data 3).

**Fig. 3:**
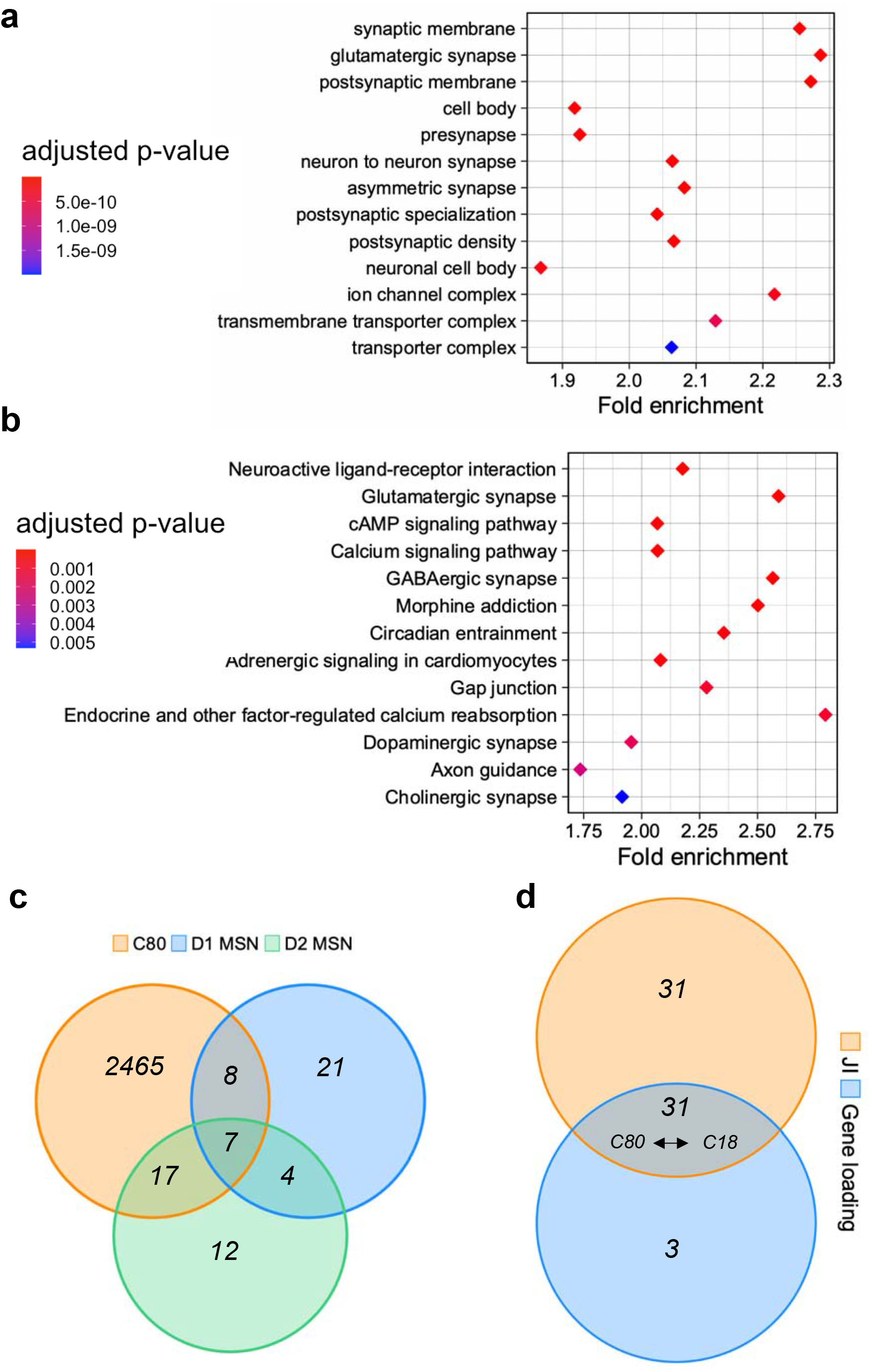
Synaptic dopaminergic specificity of C80. **a**, and **b**, Gene ontology (cellular compartment) and KEGG enrichment of C80 for both pre and post synaptic compartments as well as dopaminergic, GABAergic and glutamatergic synapses. FDR-adjusted p-values are reported. **c**, Venn Diagram showing intersection between C80 genes and genes expressed in subpopulations of D1- and D2-expressing MSNs in the nucleus accumbens. A larger intersection is found with D2-MSN over D1-MSN. **d**, Overlap between SDA components generated from the LIBD and GTEx datasets that are significantly replicated (empirical p<.001) using JI or gene loading correlation. Discovery C80 and replication C18 are one of the 4 pairs of components consistent with both JI and gene loading. *Abbreviations: JI: jaccard index; MSN: medium spiny neurons*.

To follow-up these network-level findings suggesting a role for this component in dopaminergic neurotransmission, we investigated the membership of C80 for dopamine receptor and synthesis genes. C80 included the dopamine D_2_ receptor gene *DRD2* but not *DRD1*, along with tyrosine hydroxylase (*TH*) and DOPA decarboxylase (*DDC*) genes, necessary for presynaptic dopamine biosynthesis. We found that C80 was positively correlated to *DDC* expression (t_[201]_=5.3, p=2.9×10^-7^) and negatively correlated to *DRD2* expression (t_[201]_=-3.01, p=.003) in the CN after covarying for biological and technical confounders (see *Online Methods* for details). Because DDC catalyzes the last committed step of dopamine synthesis and D_2_ receptor signaling inhibits dopamine synthesis, these results are consistent with greater dopamine synthesis capacity in individuals with greater C80, who also have higher polygenic risk for SCZ. Greater DA synthesis may thus be expected to be associated with decreased *DRD2* expression in this context.

Interestingly, when restricting the analysis to only healthy individuals, we also found that C80 negatively correlated with *DRD2* expression in the DLPFC (t_[218]_=-2.1; p=.04). We further leveraged transcript level information to disentangle to which extent the *DRD2* expression variance was related to the short or long isoform in the CN (see *Online Methods* for details), and found a significant association with the long isoform expression in consistent direction with previous gene level analyses (*DRD2* short isoform transcript: t_[183]_=-1.65, p=.1; *DRD2* long isoform transcript: t_[183]_=-2.2, p=.029). A similar result was found with only healthy individuals (*DRD2* short isoform transcript: t_[116]_=-1.38, p=.16; DRD2 long isoform transcript: t_[116]_=-2.8, p=.006). Due to the MSN enrichment and selective presence of *DRD2* compared with *DRD1* in this component, we also examined this component’s membership for the 29 most preferentially expressed genes in each MSN class identified by Tran et al.^40^ in the nucleus accumbens, with specific focus to the D1_A and D2_A clusters as they represented the largest D1-MSN (67%) and D2-MSN (87%) subclasses, respectively. We assessed the statistical significance of these intersections via permutation tests (see *Online Methods* for details). Interestingly, 17 out of 29 genes were shared between C80 and D2_A (including PENK, enkephalin typically expressed by indirect pathway MSNs^41^; empirical p < 1×10^-4^) and only 8 out 29 were shared with D1_A (with the exclusion of DRD1 and PDYN typically expressed by direct pathway MSNs; empirical p = .09), suggesting that the D_2_-expressing neuronal population may contribute more to the clustering observed in C80 (Fig. 3c).

Finally, to replicate our findings, we applied SDA to the Genotype-Tissue Expression (GTEx) dataset (https://gtexportal.org/home/)^42^. RNA-seq data are available for 120 NC across CN, DLPFC and HP (see Table 1 for demographics). This replication analysis yielded 84 components which we used to assess replication of the 69 LIBD components (see Supplementary Data 1 for SDA output). We assessed the Jaccard Index (JI), representing the overlap between gene sets, and the correlation of component-specific gene loadings. The former revealed more statistically significant replicated components (JI:62 vs gene loading:34; empirical p < .001; Fig. 3d and Supplementary Fig. 2a). Indeed, most filtered components were replicated in GTEx (90%; Fig. 3d) with a median JI = 0.13 in the framework of a gene universe overlap between the LIBD and GTEx dataset of JI = 0.67. Interestingly, C80 was among the only four replicated components out of 69 in which the associated GTEx component (C18) was consistently found using both JI and gene loading (JI = .19; gene loading R^2^ = .19; empirical p < .001; Fig 3d and Supplementary Fig. 2a). Moreover, the GTEx C18 component was again mainly active in the CN with a similar neurobiological profile by cell specificity, gene ontology, and KEGG pathway analyses and a similar enrichment for dopaminergic synapse (33 genes; fold enrichment=1.7; p_[FDR]_=.01) (Supplementary Fig. 2b-d and Supplementary Data 3). Accordingly, we replicated the association between C18 component loadings and *DRD2* expression in the CN with effect directions consistent with our discovery C80 component (t_[94]_=-2.6; p=.01). We did not find significant association with either *DRD2* long or short isoform transcripts (*DRD2* short isoform: t_[116]_=-1.9, p=.056; *DRD2* long isoform: t_[116]_=-1, p=.3; see *Online Methods* for details), suggesting that a transcript level tensor decomposition might be best suited to capture the variance at this fine-grained biological resolution.

In summary, we identified replicable co-expression patterns relative to the dopaminergic neurotransmission in a completely independent dataset of neurotypical individuals. The list of genes within discovery C80 and replication C18 components is reported in Supplementary Data 3.

### Brain functional association analysis

Based on these results in support of C80’s role in SCZ, SCZ risk, and dopaminergic function, we computed a PRS stratified for genes within C80 (C80-PRS) to examine the relationships between C80-specific SCZ genetic risk burden and neurochemical and neurofunctional parameters in the living human brain (see *Online Methods* for details about PRS computation and p-value thresholds used). C80-PRS was positively associated with greater striatal dopamine synthesis capacity as measured by [^18^F]-FDOPA specific uptake in NC and in patients with SCZ (C80-PRS1: Fisher’s z_r_ coefficient_[99.5% CI]_: 0.33_[0.01, 0.65]_; p=.0037; p_[Bonferroni]_=.037) in the whole striatum ROI analyzed in our discovery cohorts (Table 1; Fig. 4a). We also found a significant association for PRS2 in the whole striatum (C80-PRS2: Fisher’s z_r_ coefficient_[99.5% CI]_: 0.34_[0.03, 0.66]_; p=.0024; p_[Bonferroni]_=.024) (Supplementary Fig. 3a). Furthermore, both C80-PRS1 and C80-PRS2 were significantly and more strongly associated with [^18^F]-DOPA PET uptake in the associative striatum (C80-PRS1: Fisher’s z_r_ coefficient_[99.5% CI]_: 0.38_[0.07, 0.70]_ ; p=.0006; p_[Bonferroni]_=.006; C80-PRS2: Fisher’s z_r_ coefficient_[99.5% CI]_: 0.35_[0.03, 0.66]_; p=.002; p_[Bonferroni]_=.02), (Supplementary Fig. 3b-c). There was no significant correlation with limbic striatum or sensorimotor striatum when correcting for multiple comparisons. Interestingly, these results remain consistent even across different genetic ancestry definitions (Supplementary Fig 4a-b; see *Supplementary Information* for details).

**Fig. 4:**
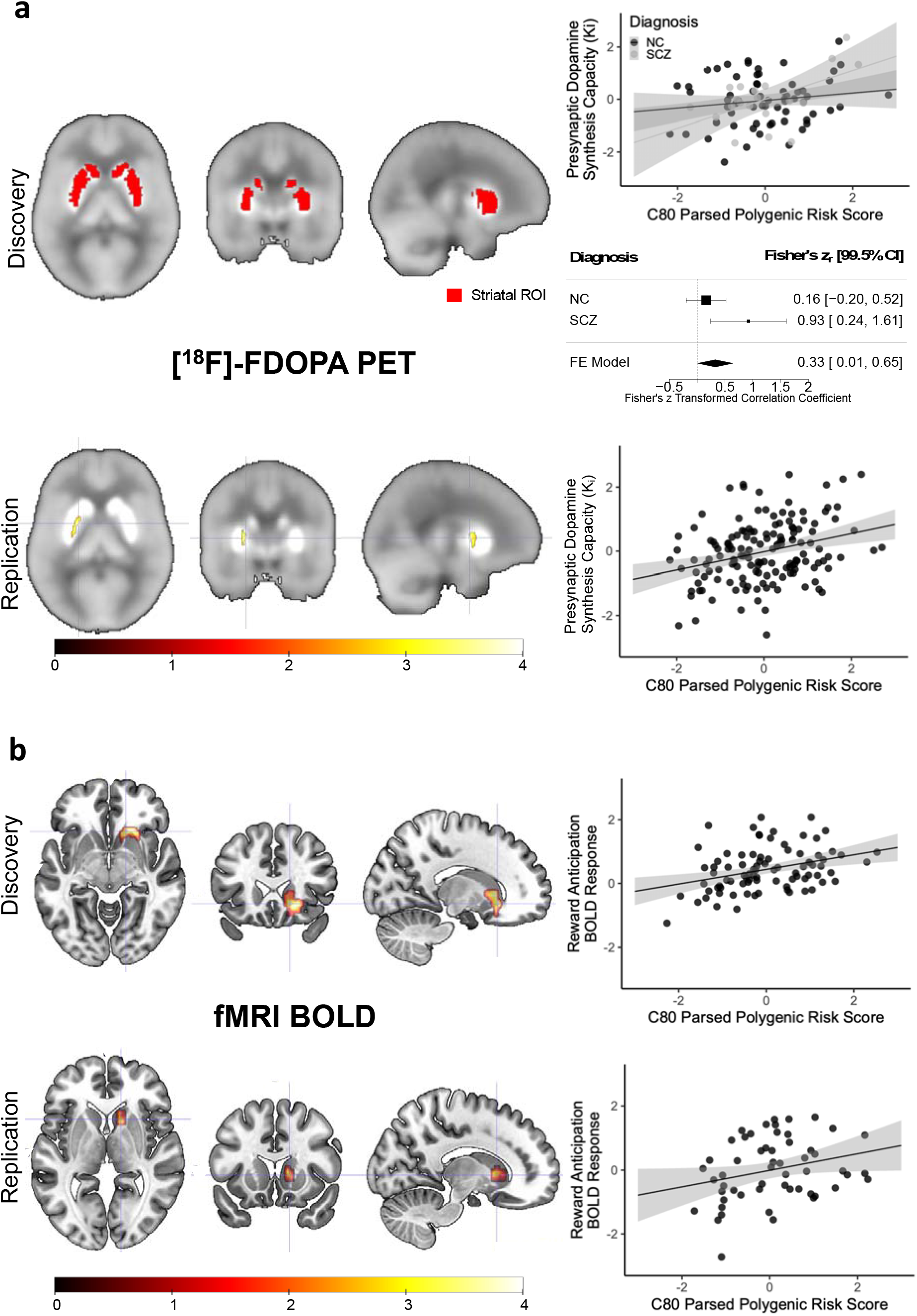
C80-PRS association with neuroimaging parameters: striatal dopamine synthesis capacity ([^18^F]-FDOPA PET) and reward anticipation-related fMRI activation (fMRI BOLD). **a**, Associations between C80-PRS and both PET cohorts are shown. First row (PET discovery): on the left whole-striatum region of interest (ROI) coverage (red) is shown overlaid on a grayscale standardized [^18^F]-FDOPA PET activity map; on the right graphs shows standardized individual mean K_i_ values for this ROI plotted against C80-PRS for the neurotypical control and SCZ subjects (upper) as well as the forest plot of the metanalysis (lower). Second row (PET replication): Region of positive association between C80-PRS and presynaptic dopamine synthesis capacity ([^18^F]-FDOPA K_i_) is shown as a statistic parametric map (color indicates t-statistic value) overlaid on a grayscale standardized [^18^F]-FDOPA PET activity map (p<0.005, uncorrected for display). Scatter plot shows standardized individual mean K_i_ values for a 2mm sphere around the peak voxel plotted against C80-PRS. **b**, Associations between C80-PRS and both fMRI cohorts are shown. First (fMRI discovery) and second (fMRI replication) rows: Regions of positive association between C80-PRS and fMRI BOLD response during reward anticipation are shown as statistic parametric maps (color indicates the threshold-free cluster enhancement (TFCE) statistics expressed in the −log10 scale). All results meet thresholds of p_[TFCE-FDR]_<0.05 and cluster extent >20 voxels. Scatter plots show standardized individual MID-related fMRI BOLD contrasts plotted against C80-PRS.

In our independent replication cohort (Table 1), C80-PRS was also positively associated with greater striatal [^18^F]-FDOPA specific uptake, albeit at a different PRS threshold (C80-PRS6: t_[149]_=3.95; k=16; p_[FWE]_<0.05), which localized to the dorsomedial striatum (Fig. 4a).

Finally, we investigated the association of C80-PRS with striatal functioning using fMRI in participants who performed a reward processing task. We found that C80-PRS1 was positively correlated to the physiological activation in the right CN during high motivation vs low motivation during reward anticipation assessed in a discovery sample of 86 NC (Table 1; C80-PRS1: p_[TFCE-FDR]_=0.01; Z=3.54; x = 17; y=15; z=-5; 73 voxels). Specifically, participants with higher C80-PRS and thus higher predicted striatal dopamine synthesis showed greater CN activation when they expected a reward during the task (Fig. 4b). We replicated this association in an additional independent sample of 55 NC (Table 1; C80-PRS1: p_[TFCE-FDR]_=0.03; Z=4.04; x = 18; y=17; z=5; 20 voxels) in a cluster partially overlapping with the one identified in the discovery sample analysis (Fig. 4b).

We also computed a measure of cumulative SCZ risk burden based on GWAS risk genes not in C80 (C80-PRS-complementary) and did not find any significant association in any of the PET and fMRI samples (p>0.05; Supplementary Fig. 5a-b). It is also worth mentioning that the number of SNPs included at each threshold for the C80-PRS is always lower than for the C80-PRS-complementary (Table 2), indicating the SNPs mapped to C80 genes represent a minority more closely involved in dopaminergic processes than the rest. Supplementary Data 4 include SNPs mapping to C80 genes and used to compute C80-PRSs.

**Table 2.** Number of SNPs used for parsed PRS. Number of SCZ risk SNPs used for the C80-PRS and C80-PRS-complementary (compl) for each cohort analyzed in the brain functional association analysis. Here are shown the first six GWAS p-value thresholds considered.

| Cohort | Diagnosis (NC/SCZ) | PRS1 (5e-08) |  | PRS2 (1e-06) |  | PRS3 (1e-04) |  | PRS4 (.001) |  | PRS5 (.01) |  | PRS6 (.05) |  |
| --- | --- | --- | --- | --- | --- | --- | --- | --- | --- | --- | --- | --- | --- |
|  |  | C80 | compl | C80 | compl | C80 | compl | C80 | compl | C80 | compl | C80 | compl |
| Discovery | NC | 89 | 141 | 174 | 277 | 588 | 1,234 | 1,347 | 3,029 | 3,444 | 8,843 | 7,309 | 19,771 |
|  | SCZ | 83 | 136 | 157 | 258 | 528 | 1,088 | 1,188 | 2,644 | 2,952 | 7,604 | 6,192 | 16,775 |
| Replication | NC | 82 | 137 | 157 | 272 | 569 | 1,186 | 1,337 | 3,001 | 3,624 | 9,394 | 8,215 | 22,094 |
| Discovery | NC | 84 | 142 | 172 | 284 | 608 | 1,276 | 1,458 | 3,268 | 3,969 | 10,228 | 8,982 | 24,090 |
| Replication | NC | 89 | 144 | 177 | 291 | 647 | 1,328 | 1,540 | 3,464 | 4,214 | 10,825 | 9,577 | 25,608 |

## Discussion

The genetic architecture of SCZ is complex and spans the genome^2^. Despite evidence for aggregation of implicated genes into certain clusters^43^, characterizing the functional biology of SCZ risk genetics has been a challenge. We applied a tensor decomposition method called sparse decomposition of arrays (SDA) to post-mortem brain gene expression data from three brain regions, i.e., CN, HP, and DLPFC in an effort to identify cohesive biological pathways that are implicated in SCZ illness and risk, which might help delineate plausible routes from SCZ-promoting genetic variation to specific neural circuit functions perturbed in this condition. We discovered a caudate-dominant co-expression gene set (C80) that is enriched for genes differentially co-expressed in individuals with SCZ relative to neurotypical controls and is associated with individual genetic risk for SCZ, features that suggest a role in SCZ pathogenesis. Expanding long-held hypotheses of dopaminergic involvement in psychosis in general and in SCZ more specifically, this gene set showed enrichment for dopamine system genes and embedded SCZ risk variation that specifically tracked with *in vivo* neurochemical and neurofunctional dopamine- and illness-related phenotypes.

The SDA algorithm provided an efficient technique to uncover sparse gene networks that were not only statistically robust but also biologically coherent, capturing, in the case of C80, gene expression covariance showing biological specialization for striatal dopaminergic circuitry implicated in SCZ. A key component of this circuitry, the C80 member gene *DRD2*, is expressed both at the presynaptic terminal, where its protein, the D_2_ dopamine receptor, functions as an autoreceptor regulating dopamine synthesis and release, as well as in the postsynaptic compartment, where its role in indirect pathway medium spiny neurons (MSNs) has been well established^44^. In line with *DRD2*’s autoreceptor role, SDA segregates *DRD2* together with the genes for the primary dopamine biosynthesis enzymes, *TH* and *DDC*, in a single, SCZ-associated component, although isoform transcript analyses did not confirm an isoform preference replicated across LIBD and GTEX data.

This is not surprising in principle, as greater DA biosynthesis should have downstream effects on both receptor isoforms. However, at the total gene expression level, greater C80 (higher SCZ risk) correlated with higher striatal expression of DDC, translating into an increase of dopamine synthesis, and as expected, lower expression of DRD2, with both results replicated in an independent NC cohort. These findings, along with the neurofunctional results, highlight the control of presynaptic DA synthesis and release as a mechanism of DA associated pathogenesis^22^. When looking at transcript level expression, we cannot clearly disentangle replicable effects specific to one isoform over the other. Importantly, while previous work on translating genetic risk into gene expression association has highlighted the presynaptic short isoform as the molecular mechanism of risk, here we are looking at co-expression in a broader biological context than genetic risk alone.

It is notable that follow-up analyses of C80 gene membership additionally identified a preference for genes expressed in the indirect pathway, i.e., D_2_ dopamine receptor bearing MSNs (e.g., *DRD2*, *PENK*) over those expressed in direct pathway associated, i.e., D_1_ dopamine receptor bearing MSNs (e.g., *DRD1*, *PDYN*)^41^. As KEGG pathway analysis revealed a collection of dopaminergic, GABAergic, glutamatergic and cholinergic pathway related genes belonging to this component, it is interesting that among the genes segregated by SDA in both C80 and C18 components is the one encoding for the M4 muscarinic receptor (*CHRM4*), previously associated with the regulation of cholinergic and dopaminergic neurotransmission in SCZ^45–48^ and recently highlighted as a potential therapeutic target for this disorder^49–51^. Besides the relevance of C80 to presynaptic dopaminergic mechanisms, these observations point to a wider biological interpretation of the genes co-expressed within this SCZ-associated component, including cortico- and nigro-striatal terminals closely tethered to the indirect pathway.

Following the identification of these links between C80 and dopaminergic systems, we conducted *in vivo* neuroimaging investigations and found that SCZ genetic risk variation that is mapped to the C80 gene set – and not cumulative risk outside of it – is specifically associated with elevated presynaptic dopamine synthesis in the striatum, which we observed in both NC and SCZ cohorts across three independent PET samples totaling 235 participants. This is consistent with the [^18^F]-FDOPA associated phenotype in SCZ^52^ and further supports the notion that C80, also expressed to a greater degree in SCZ, plays an important role in presynaptic dopamine dynamics. Thus, the present results may provide a molecular mechanism to the risk signature of a central phenotypic pillar of the modern dopamine hypotheses of SCZ^52^.

The specificity of these findings may provide insight for a better understanding of the heterogeneity of SCZ and its pathobiology. These data align with the notion that some routes to clinical illness (e.g., those within the C80 pathway) preferentially perturb presynaptic dopamine systems over others, providing a possible molecular basis for the observation that not all individuals with SCZ show excessive presynaptic dopamine synthesis capacity and not all patients respond well to antidopaminergic medications^19,20^. More generally, the approach we employed may be promising for stratifying patients based on their pathway-specific genetic liability to illness, which, if confirmed to be clinically informative, could provide new avenues for personalized medicine.

The association between these pathway-specific variants and heightened striatal dopamine synthesis, as evidenced by post-mortem and PET data, aligns with findings from a dual PET-fMRI study that demonstrated positive correlations between reward anticipation-related activation and striatal dopamine release in healthy individuals^53^. Accordingly, dopamine depletion attenuates striatal activation during the same task in healthy subjects^54^. Moreover, our results are consistent with past findings of positive correlations between polygenic risk for SCZ and ventral striatal activation during the MID task in a large sample of healthy adolescents^55^. Importantly and again, cumulative risk outside of this filter (i.e., variants not included in C80) did not show a significant relationship with this anticipatory BOLD response; along with the similar pattern observed in our PET results, this specificity suggests that parsing the PRS into co-expression pathways can reveal previously unreported phenotypic association with risk^43^.

The positive correlations between C80-specific SCZ risk burden and reward anticipation BOLD response in neurotypical individuals deviate in direction from prior findings of blunted striatal response to reward anticipation in patients with SCZ^25,26,56–60^. The difference between genetic and clinical findings may have multiple sources, including illness characteristics and pharmacological treatment. Patients with SCZ display abnormal salience attribution patterns^61^, which could lead to reduced contrast between anticipation cues and baseline, ultimately resulting in poorer motivational performance and BOLD contrast during reward anticipation^62–64^. Secondly, the activity of different brain regions involved in the reward system can be affected by the disorder^65^, while being preserved in our fMRI samples only including NC. Third, SCZ patients exhibit elevated striatal dopamine synthesis and release as measured by PET^7,12–15^, suggesting that higher steady-state dopamine levels may cause an apparent blunted response by elevating baseline activation. Consistent with this hypothesis, Knutson et al.^66^ reported that amphetamine administration, which blunts task-based dopamine release while enhancing steady-state availability in the striatum^67^, leads to decreased peak activation but prolonged activation duration during reward anticipation in healthy individuals. Taken together, we suggest that the blunted response to stimuli in a saliency-modulating task observed in SCZ may arise at least in part from both reward devaluation as well as enhanced steady-state baseline activity.

A further and perhaps paramount consideration in studies of reward circuity in patients with schizophrenia is the impact of neuroleptic medications on the sensitivity of the brain’s reward system^68,69^. Previous studies indicate that antipsychotic drugs can blunt reward-related anticipatory striatal activation in individuals with SCZ^56,57^, This effect may be associated with the blockade of D_2_ dopamine receptors^56,57^ or the suppression of dopaminergic neuron firing^70–72^, known as inactivation block. In fact, the recent report by Benjamin et al.^22^ highlights a significant downregulation of the *TH* gene in caudate of SCZ patients receiving antipsychotic drugs, and acute depletion of dopamine has been associated with reduced striatal activation during reward anticipation in both patients^73^ and controls^54^. Notably, atypical antipsychotics, which exhibit a lower level of D_2_ receptor affinity were found to enhance striatal fMRI BOLD signal during reward anticipation relative to first-generation, high-affinity D_2_ blocking agents^57,74,75^. Nonetheless, even atypical antipsychotic medications may increase baseline striatal activity in a dopamine-dependent fashion^76^. Thus, the effects of illness and pharmacological stimulation are not necessarily aligned with the relationship of illness genetic risk and striatal physiological activation in the neurotypical state. In this regard, examining effects of polygenic risk for SCZ in samples of healthy controls provided an important perspective on risk biology while avoiding important illness-associated confounders, such as treatment with D_2_ dopamine receptor antagonists.

Limitations of this study include the relatively limited sample size used in the gene co-expression analysis, which is pivotal for decomposition approaches^37^. To obtain the 3D tensor used as input for SDA, we had to exclude samples without available data in all three brain regions analyzed, a filter which was especially limiting for the GTEX replication dataset. Nonetheless, while a larger sample size would have been ideal and perhaps offered greater precision, we employed the largest gene expression resources available with these tissue types, including the relatively sizeable LIBD resource, and were able to identify replicable gene co-expression findings. Moreover, while including SCZ postmortem data in our analyses provided important insights into illness associations of identified SDA components, antipsychotic treatment and other illness-associated epiphenomena may have introduced confounding effects for the SDA analysis. We tried to address this issue by performing SDA on the same three tissues using an independent dataset consisting of only healthy controls (GTEx) to assess the rate of replication and generalizability of these results. Indeed, we found that 90% of the components identified were replicated, and the one we studied was very well reflected in GTEx. While the total sample size for [^18^F]-FDOPA PET genetic studies in this work is unparalleled, the within-cohort sample sizes are limited, which may explain minor differences in peak findings across different PRS SNP p-value thresholds or anatomically within the striatum.

Nonetheless, the convergent positive association identified between C80 and striatal presynaptic dopamine synthesis in all cohorts studied despite independent, multi-site sampling and diverse methods bolsters confidence in the PET results. Additionally, while the non-invasive reference region graphical linearization method used here for [^18^F]-FDOPA quantification has been well validated, it is possible that alternative kinetic modelling in future studies using arterial data may allow for a more comprehensive view of observed effects. Finally, it is possible that sample size limitations in the fMRI dataset prevented identification of weaker but important BOLD effects, though the replication mitigates this concern.

In sum, these results highlight a DA related striatal gene set that is prominently expressed in SCZ, implicated in SCZ genetic risk, and involved in dopamine synthesis and striatal physiological activity in vivo, suggesting that genetic risk within this pathway differentially affects SCZ-relevant striatal function. These observations provide evidence that polygenic risk for SCZ can be effectively parsed into pathways important for specific systems-level functions that is measurable even in the absence of illness^43^. Furthermore, they suggest a molecular basis of how genetic risk within the C80 pathway might affect illness relevant striatal neurochemistry and neurofunction and may open new possible avenues for studying clinical heterogeneity^19,43^ and drug treatment response^43^.

## Online Methods

### Lieber Institute for Brain Development (LIBD) postmortem data – discovery cohort

We used post-mortem human brain tissue from the LIBD Human Brain Repository. All included neurotypical controls (NC) had minimal age-associated neuropathology (determined from postmortem histopathological examination), had no substance or drug use from toxicology, and were free from any psychiatric or neurological disorder from clinical histories. All tissue donations were made with informed consent from next of kin. Brains in the LIBD Human Brain Repository were transferred from the National Institute of Mental Health Clinical Brain Disorders Branch under a material transfer agreement after having been collected under NIMH Protocol 90-M-0142, as approved by the National Institutes of Health Combined Neurosciences Institutional Review Board. An additional sample of SCZ cases in the LIBD repository were collected from the Office of the Chief Medical Examiner for the State of Maryland under State of Maryland Department of Health and Mental Hygiene Protocol 12-24, and from the Kalamazoo County Michigan Medical Examiners’ Office under Western Institutional Review Board Protocol 20111080. All samples were collected and processed using a standardized protocol specifically developed to minimize sample heterogeneity and technical artifacts as previously described^77,78^. See Table 1 for cohort demographics.

Caudate nucleus samples were derived from the anterior ‘head’ portion, a subregion tightly connected with the prefrontal cortex; HP samples from the mid-hippocampus proper (all dissections included the dentate gyrus, CA3, CA2 and CA1) plus the subicular complex; and DLPFC samples from Brodmann Area 9/46 at the level of the rostrum of corpus callosum^79^.

For all tissues, RNA sequencing was performed via the Illumina Ribozero Kit as previously described^78^. Gene-level mRNA expression was quantified as Reads Per Kilobase per Million mapped reads (RPKMs) and annotated as total gene expression separately for each brain region using GENCODE release 25 (GRCh38.p7).

We included NC and SCZ samples with European or African American ancestry, all with RNA Integrity Number (RIN) ≥ 6 (see Table 1). We used inter-array distance to identify tissue-specific outlier subjects deviating more than three standard deviations from the mean^32^ (CN = 4; HP = 7; DLPFC = 5). We then focused our analyses on mRNA expression measurements that were available for common samples (N=238) and genes (N=58,037) across all three tissues.

### The Genotype-Tissue Expression (GTEx) postmortem data – replication cohort

We used the *recount3 R^80^* package to download already processed GTEx v8 RPKMs for CN, HP, and DLPFC (Frontal Cortex Ba9). Data available for all three tissues consisted of 120 samples and 54,892 genes.

### King’s College London (KCL) PET data – discovery cohorts

Two cohorts, one with 92 NC and one with 47 individuals with SCZ, (see Table 1 for demographics) underwent [^18^F]-FDOPA positron emission tomography (PET) scans to measure dopamine synthesis capacity (indexed as the influx rate constant K_i_) in the striatum as previously described^81–86^. In short, after pretreatment one hour before the scan with fixed doses of carbidopa (150 mg) and entacapone (400 mg) to reduce peripheral tracer metabolism, and immediately following intravenous injection of [^18^F]-FDOPA, a series of dynamically binned emission frames were acquired over 95 minutes. Computed tomography (CT) imaging was performed for attenuation correction. Scans were obtained on one of the following PET scanners: an ECAT HR + 962 (CTI/Siemens, Knoxville, Tennessee), and an ECAT HR+ 966 (CTI/Siemens, Knoxville, Tennessee), and two Siemens Biograph HiRez XVI PET-CT scanners (Siemens Healthcare, Erlangen, Germany). Reconstructed, attenuation corrected emission scans were realigned to correct for interframe head motion. An atlas defining the regions of interest (striatum, its subdivisions, and the reference region (cerebellum) as described in Howes et al 2009^16^ was co-registered to a tracer specific template and transformed to each subject’s PET data series using SPM 12 software (UCL, London, UK). Time-activity curves were extracted for the regions of interest and entered into standard Patlak-Gjedde graphical linear models using the reference region to adjust for non-specific uptake to obtain the influx rate constant K_i_, a measure of specific tracer uptake^87^. The primary analyses focused on the whole striatum. For post-hoc exploratory analyses, the striatum was sub-divided into limbic, associative, and sensorimotor subdivisions based on functional topography of the striatum and its connectivity as previously descrived^88^. All participants provided written, informed consent per KCL IRB approved protocols.

### National Institute of Mental Health (NIMH) PET data – replication cohort

[^18^F]-FDOPA PET scans were acquired for a total of 150 healthy subjects (demographics in Table 1) as previously described^89^. In short, after a required 6-hour fast to prevent competition for tracer transport to the brain, a 4-hour caffeine/nicotine restriction, and pretreatment with fixed doses of carbidopa (200 mg) to reduce peripheral tracer metabolism, and immediately following intravenous injection of [^18^F]-FDOPA, a series of dynamically binned emission frames were acquired over 90 minutes. A transmission scan was performed in the same session for attenuation correction. All scans were obtained on a GE Advance PET tomograph operating in 3D mode with a thermoplastic mask applied to help restrict head movement. Reconstructed, attenuation corrected emission scans were realigned to correct for interframe head motion. Spatial warping of PET data was performed with ANTs software to an MNI space tracer-specific template. A 10 mm Gaussian kernel smoothing was applied to improve voxel-wise signal to noise ratios. Using PMOD software (https://www.pmod.com/), time-activity curves from voxels within the striatum were subjected to standard Patlak-Gjedde graphical linear modelling using cerebellar reference region time-activity data as an input function to yield K_i_ as above^87^. All participants provided informed consent per NIH Combined Neurosciences IRB approved protocols.

### LIBD fMRI data – discovery cohort

An independent sample of 86 NC (demographics in Table 1) participated in a fMRI experiment in which participants performed a modified version of the MID task^90^ based on the expectancy theory of motivation^91^. Participants had no history of any psychiatric or neurological disorders and gave written, informed consent for a protocol approved by the NIH Combined Neurosciences IRB. Participants were told that they would be monetarily compensated based on earnings in the task. Details about the task layout are reported in the Supplementary materials.

fMRI scans were acquired through a 3T GE Signa scanner. Gradient-recall echo-planar imaging was used with the following parameters: TR = 2000 ms; TE = 28 ms; flip angle = 90; 64 × 64 matrix; FOV = 240 mm; and 35 3.5 mm slices acquired with an interleaved order of slice acquisition and first five frames discarded to allow steady-state magnetization. Slice timing correction, six-parameter coregistration to adjust for movement, mean functional-image driven spatial normalization to MNI space, and spatial smoothing with an 8 mm Gaussian kernel were applied and yielded timeseries data with 3 mm isotropic resolution through SPM12 (https://www.fil.ion.ucl.ac.uk/spm/software/spm12/). A separate general linear model (GLM) was specified for each participant, modeling time-locked BOLD responses to high reward vs low reward, i.e., low expectation vs high expectation of reward event onsets, i.e., high motivation vs low motivation, by convolving the onset vectors with a synthetic hemodynamic response function as implemented by SPM12. At the model estimation stage, the data were high-pass filtered with a cutoff of 128 s to remove low-frequency drifts. Global scaling was not applied to the data.

### UNIBA fMRI data – replication cohort

A cohort of 55 NC (demographics in Table 1) participated in a fMRI experiment in which participants performed a version of the MID task^90^ alternative to the one described before. Participants had no history of any psychiatric or neurological disorders and gave informed consent for a protocol approved by the institutional ethics committee of the University of Bari Aldo Moro (UNIBA). Participants were told that they would be compensated with one gift gadget (pen, t-shirt, pin, bag, pouch, notebook) when they earned at least 1700 points, and the chance of choosing between two or three gifts of their choice (when reaching 1900 and 2300 points, respectively) and encouraged to respond as quickly as possible. For details about the task layout see Supplementary materials.

fMRI scans were acquired through a 3T Philips Ingenia scanner. Gradient-recall echo-planar imaging was used with the following parameters: TR = 2000 ms; TE = 38 ms; flip angle = 90; 64 × 64 matrix; FOV = 240 mm; and 38 3.6 mm slices acquired with an interleaved order of slice acquisition. Slice timing correction, six-parameter coregistration to adjust for movement, mean functional-image driven spatial normalization to MNI space, and spatial smoothing with a 9 mm Gaussian kernel were applied and yielded timeseries data with 3 mm isotropic resolution through SPM12. A separate general linear model (GLM) was specified for each participant, modeling time-locked BOLD responses to high reward vs low reward event onsets, i.e., high motivation vs low motivation, by convolving the onset vectors with a synthetic hemodynamic response function as implemented by SPM12. At the model estimation stage, the data were high-pass filtered with a cutoff of 128 s to remove low-frequency drifts. Global scaling was not applied to the data.

### Genotype data processing and polygenic risk score (PRS) calculation

Genotype data for all samples were obtained and processed as previously described^92,93^. Genotype imputation and quality checks as well as calculation of genomic eigenvariates (GEs) for population stratification were performed in each cohort separately. See Supplementary Information for details.

We indexed the whole-genome genetic risk for SCZ by computing the PRS for each sample using the PRSice-2 software^94^. To obtain a highly informative SNP set with as little statistical noise as possible, we excluded uncommon SNPs (MAF < 1%), low-quality variants (imputation INFO < 0.9), indels, and SNPs in the extended MHC region (chr6:25-34 Mb). We used PGC (wave 3; primary autosomal analysis) GWAS^2^ summary statistics that did not include any of the LIBD discovery samples (leave-sample-out) to weight SNPs by the effect size of association with SCZ. We used European samples from the 1000 Genomes Project^95^ (1000G) as external reference panel to improve the linkage disequilibrium (LD) estimation for clumping. Both PGC3 leave-LIBD-out and the reference panel were in reference to human genome Build 37.

To stratify SCZ genetic risk for genes within a specific component we first mapped European 1000G SNPs at 100kbp up- and down-stream of each component-specific gene using MAGMA tool (v1.09b), we then matched component-specific SNPs with PGC3 leave-LIBD-out summary statistics and finally computed the scores for the KCL, NIMH, LIBD- and UNIBA-fMRI cohort separately using PRSset^96^ and again the European 1000G as LD reference panel. As negative control, we computed complementary scores including all PGC3 leave-LIBD-out SNPs not mapping to any of the component-specific genes.

We used PRSs based on 10 SNP sets corresponding to GWAS SNP association p values of p = 5e−8 (PRS1), p = 1e−6 (PRS2), p = 1e−4 (PRS3), p = 0.001 (PRS4), p = 0.01 (PRS5), p = 0.05 (PRS6), p = 0.1 (PRS7), p = 0.2 (PRS8), p = 0.5 (PRS9), and p = 1 (PRS10)^2^. Table 2 shows number of SNPs used for each cohort for each PRS threshold.

### RNA data processing

To analyze LIBD postmortem data with SDA, we first removed mitochondrial genes and genes with RPKM expression median lower than 0.1 or deviating more than 3 standard deviations from the mean in each tissue. We then removed genes with more than 20% zeroes in all three tissues as previously done^35^. We log-transformed RPKM values with an offset of 1, i.e., log2(RPKM+1). After performing quantile normalization to normalize samples based on their gene expression, we rank-normalized gene expression using Blom formula^28,29,97^ to limit the impact of deviations from normality in expression data. We performed all normalization steps for each tissue separately. The final tensor of 22,356 mRNA expression measurements in 238 samples and across CN, HP and DLPFC was used as input for SDA (see Table 1).

### Sparse decomposition of arrays (SDA) computation

We iterated the algorithm 10 times, and for each run we obtained latent components defined by three bidimensional matrices: i) the individual score matrix, which represented the magnitude of the effect of each component across individuals and was used to compute the association with diagnosis; ii) the tissue score matrix, which indicated the activity of the component for each tissue and was used to identify the contribution of each tissue to the components; iii) the gene loading matrix, which indicated the contribution of each gene to components and served to identify genes specific to tissues or shared between them (see *Supplementary Information* for further details about parameters used).

We thus obtained robust components found consistently across multiple iterations, whereas others only occur in one or a few of them. We clustered similar components across different iterations following published procedures^35^ (see *Supplementary Information*). We obtained 126 large clusters containing components from multiple different iterations and combined components within each cluster by taking the mean of the individual scores, tissue scores and gene loadings. We finally used the resulting 126 combined clusters as the basis for further analyses.

### Tissue activity evaluation

We evaluated the tissue loadings across components by column-wise scaling the tissue score matrix (components as columns and tissues as rows) obtained by the SDA decomposition so that the largest score was equal to 1 and the lowest to −1 using a threshold of |0.5| (as previously reported by SDA developers^35^) to infer the tissue specificity of each component and how many components are shared across tissues (Fig 2b and Supplementary Fig. 1a).

### Confounder analysis

Since SDA identifies non-sparse components that might be expected to arise from confounding effects, we expected singular value decomposition to reveal latent confounders. Singular value decomposition in its principal component analysis implementation has been used often to this aim^35,98–100^. To identify components most likely to represent confounding effects, we computed a series of multiple linear regressions using as dependent variable individual scores from the individual score matrix for each of the 126 SDA components and as predictors both biological confounders (age, sex, diagnosis, first 10 GEs) and technical confounders (postmortem interval (PMI), RIN, mitochondrial mapping rate, rRNA rate, gene mapping rate).

We adopted a confounder detection approach consistent with the reference paper^35^ by using the same confounder effect size of 0.274 (p = 10^-10^; sample size = 845). We found that the same effect size corresponded to p < 5×10^-4^ in our sample of 238 individuals (observed power = 80%) and removed the 57 components associated at this threshold with at least one of the technical confounders or GEs. We focused our further analyses on the remaining 69 components.

### Diagnosis and PRS association

To investigate whether the 69 components were differentially co-expressed between NC and SCZ, we tested the association of the component-specific individual scores with diagnosis via ANCOVA while covarying for biological (age and sex) and tissue specific technical confounders (PMI, RIN, mitochondrial mapping rate, rRNA rate, gene mapping rate), taking into account the component tissue activity. Samples with age > 17 were included (N=229) as this was the minimum age in the SCZ sample.

We further evaluated the association of the differentially co-expressed components with PRS via multiple linear regression again covarying for age, sex, diagnosis, and tissue specific confounders and including only samples with European ancestry (N=103) since the summary statistics used are mainly based on European population. For this analysis one-tailed tests were used because of the constraint on effect directionality, i.e., we discarded potential significant results in the opposite direction of diagnosis association. We focused on PGC3 variants with a SCZ association p-value < .05, since this PRS has been shown to have the highest prediction accuracy for diagnostic status in multiple independent samples^2^.We used Benjamini-Hochberg false discovery rate (FDR) correction to correct for multiple comparisons across SDA components and set α_FDR_ < 0.05.

### Biological and functional enrichment analysis

We explored the functional and biological significance of these components through enrichment analyses for multiple psychiatric disorders and immune disorders’ top risk loci genes, i.e., putative causal genes identified by setting a fixed distance around each index GWAS-significant SNP and subsequently integrating genomic functions or chromatin interactions^101,102^ (ADHD - attention deficit hyperactivity disorder^103^; ASD - autism spectrum disorder^104^; BD - bipolar disorder^103^; MDD - major depressive disorder^105^; OCD – obsessive-compulsive disorder^106^; PTSD – post-traumatic stress disorder^107^; SA – suicide attempt^106^; SCZ-schizophrenia^2^; CD and UC – Crohn’s disease and ulcerative colitis^108^).

We also computed enrichments for differentially expressed genes (DEGs) obtained from CN^22^, HP and DLPFC^78^; genes proximal to differentially methylated CpG islands (DMGs) in PFC and blood^77,109–113^ and loss of function intolerant genes^114^. For DEGs, we performed a brain region-specific enrichment using the appropriate gene list of each tissue. Moreover, for DLPFC DEGs and DMGs enrichment, we computed a meta-analysis of the papers from which we retrieved target genes to obtain module-wise enrichment p-values (sum-log Fisher’s method). Considering the overlap between the SDA components obtained, we computed permutation statistics to control for multiple comparisons by first creating for each component a null distribution of 10,000 sets of randomly sampled genes using the 22,356 genes as the universe and then comparing the enrichment hits to the null distribution created from the permuted components (*⍺* =.05).

### MAGMA analysis

We used the MAGMA tool v1.09b, pathology-specific summary statistics as SNP p-value data and 1000G European as the reference data file for a European ancestry population to estimate LD between SNPs. We took the following steps: i) we mapped 1000G SNPs to genes encompassed in each component (a window of 100 kb upstream and downstream of each gene; for H-MAGMA we used Adult brain Hi-C annotation files already computed in the H-MAGMA publication^39^), ii) we calculated gene-wide association statistics based on summary statistic SNPs p-values (MAGMA “mean” method), iii) we performed “competitive” gene-set enrichment analysis where the association statistic for genes in the components is compared to those of all other genes with at least one SNP mapped (universe used consisted of 22,356 genes used for SDA). FDR correction served to control for multiple comparisons (*⍺*_FDR_=0.05).

### Cell-type specificity analysis

We further asked whether SDA components mapped onto specific brain cell-types. We used marker genes already identified by Skene et al. including cell-type specificity indices^115^. They computed specificity indices for each gene ranging between 1 (high specificity for a given cell type) and 0 (low specificity). We used specificity indices derived from single-nuclei RNA-seq of human brains^116^, which discriminated ten different cell types (neuron and glia); and from single-cell RNA-seq of mouse brains^115^ which encompasses 24 different cell types. We used Mean-rank Gene Set Test in the *limma R* package to evaluate the enrichment of our components for the specificity indices of each cell type. This algorithm performs a competitive test comparing the specificity index rank of the co-expressed genes with the remaining genes. FDR correction served to control for multiple comparisons across components and cell-types tested (*⍺*_FDR_=0.05).

### Gene Ontology analysis

Finally, we explored the gene ontology of components via enrichment analysis through the

*clusterProfiler R^117^* package using the Gene Ontology Database (PANTHER, http://pantherdb.org)^118^ and the Kyoto Encyclopedia of Genes and Genomes (KEGG, https://www.genome.jp/kegg/) database and setting the 22,356 genes gave as input to SDA as universe. FDR correction was applied to control for multiple comparisons (*⍺*_FDR_=0.05).

### Association with DRD2, DDC and TH total gene expression

We evaluated the C80 association with *DRD2*, *DDC* and *TH* gene expression via a multiple linear regression analysis using C80 individual scores as dependent variable and *DRD2*, *DDC* and *TH* expression across CN, DLPFC and HP as independent variables. We also included age, sex, diagnosis, postmortem interval (PMI), tissue-specific RIN, mitochondrial mapping rate, rRNA rate and gene mapping rate as covariates. Finally, we added to this model the interaction between diagnosis and each gene expression as well as interaction between diagnosis and age to control for spurious association driven by postnatal samples (Eq.1). The gene expression values used were the ones given as input to SDA for all 238 samples (quantile and rank normalized matrices). We also performed this analysis using only healthy individuals (N = 154; Eq.2).

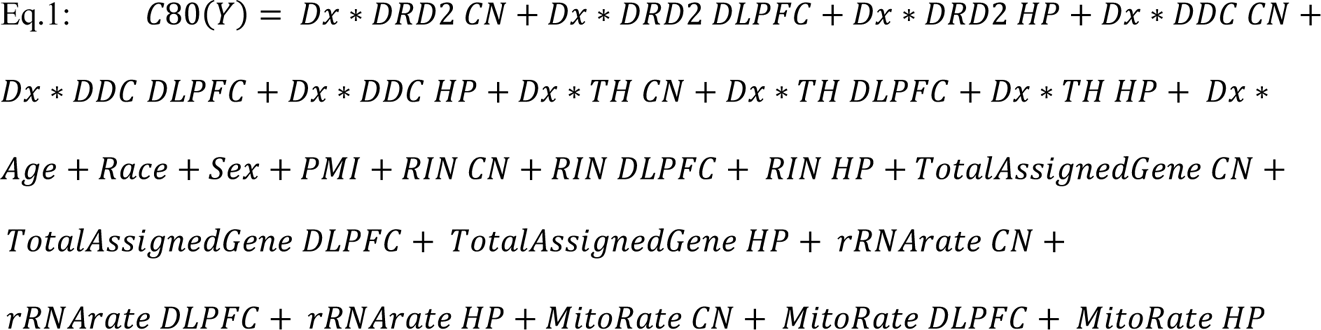

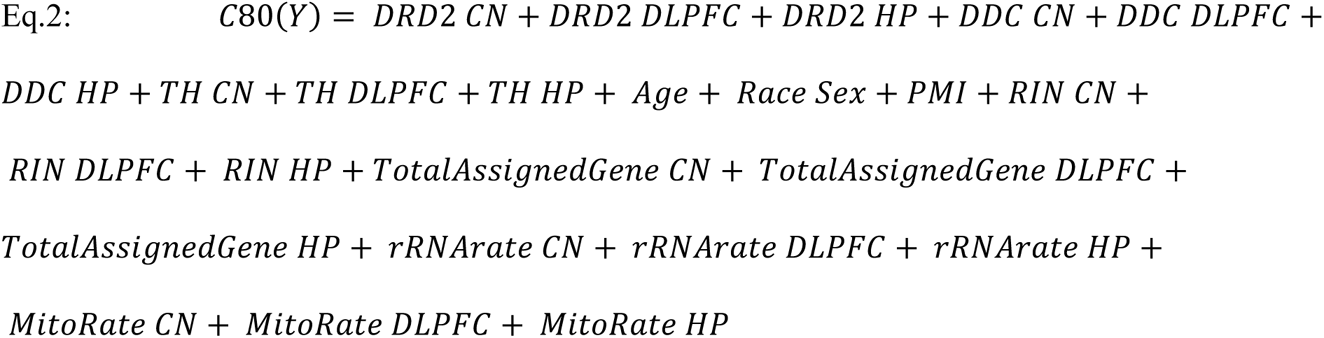

### DRD2 transcript level association analysis

To evaluate contribution of short and long isoform to the *DRD2* expression variance, we substituted the *DRD2* terms in the previous gene level models (Eq.1-2) with the long and short isoform transcript expression values (Eq.3).

Transcript counts were preprocessed and normalized to transcripts per million (TPM) estimates as previously described^22,78^ and were available for 222 out of the 238 samples previously used. After mapping 138,933 transcripts to the 22,356 genes used for previous analyses, we log-transformed TPM values with an offset of 1, i.e., log2(TPM+1) and kept transcripts with a median higher than zero. We then performed quantile and rank normalization in each tissue separately as previously done. *DRD2* short isoform survived filters for all tissues while the long isoform had a median higher than zero only in CN. This analysis was also performed using only healthy individuals (N = 143).

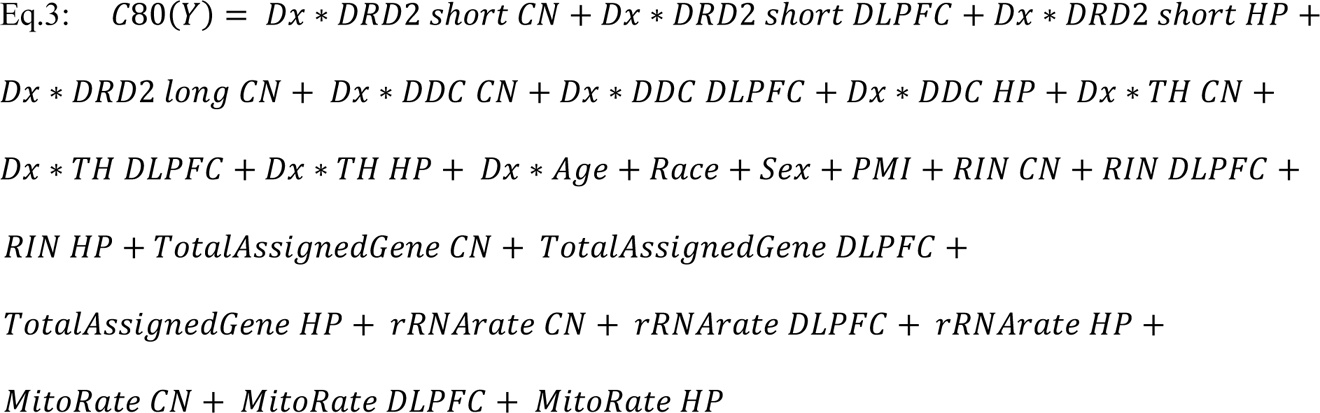

### MSN pathways enrichment

We downloaded the 40 most preferentially expressed genes in each MSN class identified by Tran MN et al.^40^ in the nucleus accumbens and focused on D_1__A and D_2__A clusters as they represented the largest D_1_-MSN (67%) and D_2_-MSN (87%) subclasses, respectively. As 11 genes shared expression for both D_1__A and D_2__A clusters, we considered the intersection without these genes for a total of 29 genes in each class. We permuted 10,000 gene sets matching both C80 component size and GC content, gene length and average expression distributions of C80 genes. The universe from which random genes were pooled consisted of the protein coding genes given as input to SDA for which this info was available (22,282 genes).

We then computed empirical p-values by comparing the enrichment hits with each MSN cluster to the null distribution created from the permuted gene sets (*⍺* =.05).

### GTEx replication analysis

To replicate gene co-expression sets obtained with the LIBD data, we applied SDA on CN, HP, and DLPFC GTEx RNA-seq data using the exact same pipeline previously described. The input matrix for SDA is described in Table 1.

Two replication measures were assessed: correlation between LIBD and GTEx component-specific gene loadings and Jaccard Index (JI) as the intersection/union of the LIBD and GTEx component-specific genes. To identify the LIBD-GTEx pair of replicated components we took for each of the LIBD components the GTEx component with the highest replication measure assessed and iteratively discarded that component to have unique LIBD-GTEx pairs. We then permuted the LIBD components 10,000 times and compared the replication measure previously assessed to the null distribution created from the permuted components to obtain a replication empirical p-value for each pair identified.

### Replication of total gene and transcript level expression in GTEx

To replicate results obtained in the discovery sample, we assessed C18 association with *DRD2* total gene expression via a multiple linear regression analysis using C18 individual scores as dependent variable and *DRD2* expression across CN, DLPFC and HP as independent variables as previously done in the discovery analysis (Eq.2).

*DRD2* long and short isoform transcript association was performed as previously done in the discovery analysis (Eq.3).

We downloaded GTEx v8 transcript TPM values from GTEx portal (https://gtexportal.org/home/datasets) and after mapping 133,788 transcripts to the 20,475 genes used we performed normalization steps as previously done in the discovery analysis.

### Ancestry stratification

As the summary statistics used are mainly based on the European population and PRS association with other ethnicity groups might lead to biases, we evaluated the individual ancestry based on the genotype data rather than only considering the self-reported ethnicity for all cohorts included for the brain function association analysis. To this purpose, we used a procedure developed by the ENIGMA consortium that consists in performing a PCA on target data merged with the HapMap^119^ phase 3 reference dataset (https://enigma.ini.usc.edu/wp-content/uploads/2012/07/ENIGMA2_1KGP_cookbook_v3.pdf). For this analysis we included all samples whose genotype information were available (KCL: 168; NIMH: 169; LIBD: 86; UNIBA: 2,178; see Supplementary Table 1). We then computed an individual ancestry score based on the GE obtained from the PCA analysis. We trained a generalized linear model using the *glmnet R* package; we used the first 20 GE obtained as predictors and the ethnicity (European = 1; Others = 0) as response variable and considered only samples in the reference dataset. Then, we used the trained model to predict the ethnicity of our samples using the first 20 GE. Finally, for each subject we obtained a European ancestry score and considered a threshold of 90% prediction probability to remove individuals with a non-Caucasian ethnicity (KCL: 59; NIMH: 3; UNIBA: 213). Remaining samples whose genotype and PET/fMRI data were available were used in further analyses (demographics in Table 1).

Since the KCL discovery cohort was the most heterogeneous in terms of ethnicities included we decided to also evaluate different ancestry subdivisions based on the visualization of the first two PCA dimensions, i.e. top axes of variation (see details in *Supplementary Information*).

### Parsed-PRS association with PET data

Considering the sample heterogeneity in the KCL discovery cohort in terms of both ethnicity and scanners used and population type at the diagnostic level, we conducted the association analysis separately in the NC and SCZ samples. A multiple linear regression served to associate the C80 stratified PRS (C80-PRS) as well as its complementary score (C80-PRS-complementary) with [^18^F]-FDOPA uptake in the striatum indexed by Ki using age, gender, cannabis use, scanner and first three GEs as covariates. We then combined the effect of the individual studies with a fixed-effect model meta-analysis, as random-effect models require data to be randomly extracted from equivalent populations, an assumption that does not hold for clinical and control cohorts. We converted t-statistics from the regression model into correlation coefficients using the following formula:

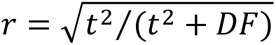

where DF is number of the degrees of freedom for the t-statistic. We finally used Fisher’s r-to-z transformed correlation coefficients as outcome measure in the *metafor*^120^ *R* package.

We focused our analysis on dopamine synthesis in an ROI encompassing the whole striatum and to obtain a more granular view of the relationship between risk and phenotypes corrected for multiple comparisons using Bonferroni method (α = .05/10). We then explored this association also in different striatum subdivisions as well as across different ancestry definitions (see *Supplementary Information*).

As in the KCL analyses described above, we performed separate multiple linear regression analyses for C80-PRS and C80-PRS-complementary predictors in the independent NIMH NC cohort, in each case using the same covariates as described above in the KCL analysis (i.e., age, sex, and the first three GEs; the ‘scanner’ variable was omitted, as all scans were obtained on a GE Advance PET scanner). Comparisons were conducted voxelwise across the whole striatum, using SPM software at a height threshold of p<0.05, voxelwise family-wise error (FWE) corrected for multiple comparisons.

### Parsed-PRS association with fMRI data

Finally, we associated C80-PRS and C80-PRS-complementary to reward anticipation-related fMRI activation in the independent sample of 86 NC from LIBD. We used the data of 55 NC from UNIBA participants to replicate the results.

BOLD responses to events of interest were modelled separately and time-locked to event onset by convolving the onset vectors coinciding with onset of events (including cues by type, button press, successful/unsuccessful outcomes, and error trials) with a synthetic hemodynamic response function as implemented by SPM12. For all analyses, the primary outcome measure was the contrast in BOLD signal of rewarded relative to control cue events, which best reflects reward anticipation responses in this task. Participants additionally completed cognitive testing outside of the scanning environment that assessed full scale IQ, which was included as a nuisance covariate in analyses.

Age, sex, IQ and first three GEs were used as covariates, consistently with previous analyses, whereas MID-related BOLD signal (cue-related anticipatory response during reward versus control trials) was the dependent variable. Cue-related individual contrasts of the 86 NC from LIBD were entered into a group level analysis to identify voxels with a significant effect of C80-PRS and C80-PRS-complementary on reward anticipation through separate multiple regression performed with SPM12. We considered the threshold free cluster enhancement correction^121,122^ p_[TFCE-FDR]_ < 0.05 accounting for multiple comparisons as the number of voxels within the task-related activity mask derived by the one sample t test on cue-related individual contrasts (p_FWE_=0.05). Next, the cue-related individual contrasts in the 55 NC from UNIBA were associated with C80-PRS, using age, sex, IQ and first three GEs as covariates. We considered significance at p_[TFCE-FDR]_ < 0.05 accounting for multiple comparisons as the number of voxels within the task-related activity mask derived by the one sample t test on cue-related individual contrasts (p_FWE_=0.05).

## Supporting information

Compressed zipped directory containing output of SDA computed on the LIBD discovery and GTEx replication datasets

Excel file with LIBD discovery components summary information, association with both biological covariates and technical confounders (Table S1), and c

Excel file of discovery C80 (Table S1) and replication C18 (Table S2) genes as wells as relative GO and KEGG enrichment results (Table S3 and S4)

Compressed zipped directory containing SNPs used to compute the C80-PRS

Compressed zipped directory containing scripts used for this study.

## Data availability

SDA software is publicly available at: https://jmarchini.org/software/#sda.

LIBD post-mortem processed RNA-seq data is available at: https://eqtl.brainseq.org/phase2/ and at: https://erwinpaquolalab.libd.org/caudate_eqtl/.

GTEx post-mortem processed RNA-seq data is available at: https://gtexportal.org/home/datasets. SDA input and output data, components summary information, GO enrichment results, and SNPs used to compute C80-PRSs (Supplementary Data 1-6) are available at: https://doi.org/10.5281/zenodo.8214643

## Code availability

The scripts used for this study are available in Supplementary Data 5 at: https://doi.org/10.5281/zenodo.8214643

## Acknowledgements

We are grateful for the contributions of the Office of the Chief Medical Examiner of the State of Maryland, Office of the Chief Medical Examiner of Kalamazoo County Michigan, Office of the Chief Medical Examiner University of North Dakota School of Medicine, Gift of Life of Michigan, Office of the Chief Medical Examiner of Santa Clara County California, and Medical University of Sofia, Bulgaria in assisting the Lieber Institute for Brain Development in the acquisition and curation of brain tissue donations for this study. All research at the Lieber Institute for Brain Development is made possible by generous gifts from the families of Steve and Connie Lieber and Milton and Tamar Maltz. We would like to thank all the family members of the donors for their exceptional contribution. We would like to acknowledge Dr. Peter Herscovitch and the staff of the NIH PET Center for their data acquisition support. We are appreciative of all the neuroimaging research volunteers for their generous participation.

We are in debt to A. Jaffe, who has contributed to this work by offering data and software. We are grateful to Dr. Christopher Borcuk, Dr. Pasquale Di Carlo, Dr. Piergiuseppe Di Palo, Andrea Gaudio, Dr. Marco Papalino, and Prof. Paolo Taurisano for exploratory analyses and exchanges of ideas that have influenced this work.

## Author information

### Contributions

Conceptualization: L.S., G.P., and D.R.W. Data curation: L.S., D.E., R.P., E.D., L.A.A., J.C., C.F.Z., J.H.S., O.H. Formal analysis: L.S., D.E., R.P., G.P. Funding acquisition: G.P., D.R.W., and A.B. Investigation: L.S., D.E., G.P., A.B., K.B., D.R.W. Methodology: L.S., D.E., R.P., L.A.A., J.H.S., Q.C. Project administration: G.P. Resources: J.C., A.G., L.A.A., M.G., K.G., T.H., J.K., A.F.P., T.P., A.R., M.V., W.U. Software: L.S., D.E. and R.P. Supervision: G.P. and D.R.W. Visualization: L.S., D.E. and R.P. Writing (original draft): L.S, D.E., R.P., G.P., and D.R.W. Writing (review and editing): all authors.

## Supplementary Information

Supplementary Notes, Supplementary Data 1-6, Supplementary Figs 1-5, Supplementary Table 1, Supplementary References.

