## Supplementary Information for "Dopamine and schizophrenia from bench to bedside: Discovery of a striatal co-expression risk gene set that predicts in vivo measures of striatal function"

855 North Wolfe St

21205 Baltimore, MD

**This File includes:**

Supplementary Notes, Supplementary Data 1-6, Supplementary Figs 1-5, Supplementary Table 1, Supplementary References.

### Supplementary Notes

#### fMRI task layout

LIBD cohort. The general layout of a trial in this version of the MID task is divided into three phases: cue (2000-3000 ms), target (up to 1250 ms in the 'difficult condition', while up to 1750 ms in the 'easy' condition), and outcome (450 ms). Immediately prior to scanning, participants were instructed on the task to be performed in the scanner, including being explicitly informed of the meaning of each cue. Cues indicated, via background color, whether the trial would be more likely difficult, more likely easy, i.e., how long the target would likely stay on the screen, and the reward magnitude, i.e., $2 or $0 (control trial). The target was the appearance of paper currency (or blank paper in the control condition) over a money safe. Participants were instructed to respond with a single button press with their right thumb when the target appeared, and if the target was hit, i.e., response was made while the target was on the screen, the target (money bills or blank paper) fell into the money safe (450 ms). On the contrary, a trial was considered an error if the participant: a) did not respond to the target, b) pressed more than once to the target, or c) pressed prior to target appearance. In these cases, the target (money bills or blank paper) disappeared without falling into the money safe and participants did not receive money towards final payment. The outcome phase provided feedback for money earned from that trial. Intertrial intervals (2500-4500 s) were calculated to maintain the same number of trials regardless of reaction time variability. An extensive description of the task is provided by Kholi et al. 2018[1].

UNIBA cohort The general layout of a trial in this version of the MID task is divided into three phases: cue (2000 ms), target (up to 500 ms), and outcome (2000 ms). Immediately prior to scanning, participants were instructed on the task to be performed in the scanner, including being explicitly informed of the meaning of each cue. The target presentation duration was calculated based on individual performance during the training phase to achieve at least 67% success rate. Cue stimuli consisted of three different white geometric shapes: a full circle, representing a chance of gain 100 points (reward condition), an empty circle, representing a chance of gain 0 point (control condition) and a full square, representing a chance of losing 100 points (punishment condition). The target was the appearance of a white triangle. Immediately after the response, feedback appeared for 2000 ms documenting whether the participant had won or lost points as well as their cumulative total at that point. Participants were instructed to respond with a single button press with their right thumb when the target appeared to gain or to not lose points. A trial was considered a hit, when the response was made while the target was on the screen. On the contrary, a trial was considered an error if the participant: a) did not respond to the target, b) pressed prior to target appearance. The outcome phase provided feedback for money earned from that trial and the total amount earned so far.

#### Genotype data processing

LIBD, NIH and UNIBA cohorts. Participants underwent blood withdrawal for subsequent DNA extraction from peripheral blood mononuclear cells. To this aim, approximately 20ml of fresh blood was obtained through a conventional venous blood collection with 10ml EDTA Vacutainer Venous Blood Collection Glass Tubes (Vacutainer ®). Approximately 200 ng DNA was used for genotyping analysis. DNA was concentrated at 50ng/ l (diluted in 10 mM Tris/1mM EDTA) with a Nanodrop Spectrophotometer (ND-1000). Samples were genotyped using variate Illumina Bead Chips including 510K/610K/660K/2.5M.

Quality control was performed on the cohorts separately using PLINK (version 2; http://pngu.mgh.harvard.edu/purcell/plink/)[2] according to standards developed by the Psychiatric Genomics Consortium[3] including SNP missingness < 0.05 (before sample removal); subject missingness < 0.02; autosomal heterozygosity deviation (|Fhet| < 0.2; for NIH cohort |Fhet| within 3.5 standard deviations); SNP missingness < 0.02 (after sample removal), SNP Hardy-Weinberg equilibrium (HWE: P > 10 í6) and minor allele frequency (MAF) > 0.01. Furthermore, the degree of recent shared ancestry, i.e., the identity by descendent (IDB)[4], has been estimated within the cohorts to define the relatedness of all pairs on individuals through the PLINK function '--genome'[5]. The threshold 0.125 (for NIH cohort 0.185) represents the relatedness of 3rd degree[6] that was used as cutoff to exclude possible influence of relatedness within cohorts on dependency between observations[7]. Of each pair of related individuals, the one belonging to the group with greater numerosity within each cohort is dropped from the final datasets.

Genotype imputation was performed using the pre-phasing/imputation stepwise approach implemented in IMPUTE2 / SHAPEIT (default parameters and chunk size of 3 Mb for LIBD and UNIBA - 250 Kbp for NIH) and using Phase 3 1000 genome as reference panel[8,9]. After imputation, imputed dosage data for each SNP with imputation quality (INFO) > 0.9 were used for polygenic risk scores (PRS) calculation.

For LIBD cohort, the first 10 principal components of the whole genome data were calculated using EIGENSOFT v5.01 (EIGENSOFT, http://www.hsph.harvard.edu/alkes-price/software/) and considered as nuisance covariates in genetic analysis while for UNIBA and NIH cohorts the SNPRelate R package[10] and PLINK (version 2) were respectively used..

<u>KCL cohort.</u> DNA was extracted from whole blood samples or cheek swabs using standard procedures[11]. Genotyping was performed at Cardiff University, using HumanCore Exome 1.1 arrays (“Psych-chip”, Illumina, San Diego, California, USA). Genotype quality control (QC) was performed according to standard parameters[12].

PLINK (version 1.9) was used to implement pre-imputation quality control procedures. Individuals were excluded from the sample if estimates of the inbreeding coefficient F indicated ambiguous sex (F = 0.2 - 0.8), or if there was a discrepancy between their genotypic and self-reported sex. Any discrepancies were checked with the data collection site to confirm no errors were made during manual entry of phenotypic data. Samples with a genotype call < 95%, or high pairwise relatedness (pi-hat > 0.1) were excluded. SNPs were removed if MAF < 0.01, if deviation from HWE at p-mid ” 1e-6, or if minimum marker call rate < 95%.

Chromosome files were uploaded to the Michigan Imputation Server (https://imputationserver.sph.umich.edu/index.html#!) where they underwent in house quality control before being passed through imputation pipeline (see:

https://imputationserver.readthedocs.io/en/latest/). Data passing QC were phased using Eagle[13] v2.4 and imputed using minimac4 (https://genome.sph.umich.edu/wiki/Minimac4) with the HRC-r1.1 reference panel[14] (GRCh37/hg19 array build), with population set to mixed. SNPs were excluded if the imputation INFO score was < 0.9, MAF < 0.10, or HWE P < 1e-6, genome = 0.05.

The Principal Components Analysis in Related samples (PC-AiR) method[15] was used in R (GENESIS R/Bioconductor package[16]) on the full set of genotypes to generate the top 10 principal components of the sample.

#### SDA run and post-processing

We run SDA 10 times using the following parameters:

-- N = 238 (number of samples)

-- max_iter = 3000 (number of iterations)

-- num_comps = 238 (number of components)

-- seed = 400 (used for results replication)

Running the method multiple times, we obtained some components that are found consistently across multiple runs, whereas other components only occur in a small number of them. To identify robust components, the authors of the paper implemented a method that clusters similar components across different runs. We then focused on large clusters containing components from multiple different runs and used these as the basis for further analyses. More specifically, at the end of each run we stored the individual and tissue scores, gene loadings and Posterior Inclusion Probabilities (PIPs = a ranking measure to assess whether the data favors the inclusion of a variable in the regression model, i.e., the probability that, after the SDA decomposition, each gene with his loading or weight belongs to a specific component). After the 10 runs finished, following what the authors did in the paper, we calculated the absolute correlation between the individual scores for all pairs of components across the ten runs. Hierarchical clustering was then used to group components into clusters, using one minus the absolute correlation as a dissimilarity measure. The clustering was terminated when no correlations between clusters were above 0.4. Although authors of SDA used 0.6 as threshold, we decided to use a more lenient one to maximise the information considering our significantly smaller sample size. The components within each cluster were then combined. We took the mean of the individual scores, tissue scores and gene loadings and the median PIPs to obtain the most representative value within each cluster (centroid). We finally obtained 126 robust components and we used PIP threshold > .5 to identify genes within each component.

#### Parsed-PRS association with KCL PET data across different ancestry clusters

Since the KCL discovery cohort was the most heterogeneous in terms of ethnicities included (Supplementary Table 1), together with the ancestry stratification reported in the main text for the main analyses, we also decided to evaluate different ancestry subdivisions based on the visualization of the first two PCA dimensions, i.e. top axes of variation. Following what we reported in the main text (see Online Methods), we used a procedure developed by the ENIGMA consortium that consists in performing a PCA on target data merged with the HapMap121 phase 3 reference dataset (https://enigma.ini.usc.edu/wp-content/uploads/2012/07/ENIGMA2_1KGP_cookbook_v3.pdf). For this analysis we included all KCL samples whose genotype information was available (N: 168). We then plotted the first two PCs against each other and defined two different clusters: one excluding individuals belonging to the African (AFR) and Euroasian (EA) ancestry group (PC1 cutoff = .048; PC2 cutoff = .013; N = 102; Supplementary Fig. 4a) and one including only individuals within the European (EUR) ancestry group (PC1 cutoff = 0; PC2 cutoff = -0.049; N = 88; Supplementary Fig. 4a). Finally, after matching samples based on imaging data availability, for each of these clusters as well as for the whole KCL cohort, we evaluated the association of the C80 stratified PRS (C80-PRS) as well as its complementary score (C80-PRS-complementary) with [18F]-FDOPA uptake in the striatum indexed by Ki in the same identical fashion as the one reported in the main text (see Online Methods for details).

We found C80-PRS positively associated with greater striatal dopamine synthesis capacity as measured by [18F]-FDOPA specific uptake in NC and in patients with SCZ in the whole striatum and other subdivisions ROI analysed in both ancestry clusters as well as the whole KCL cohort (Supplementary Fig. 4b). Interestingly, no significant association was found with complementary C80-PRS.

### Supplementary Data

**Supplementary Data 1.**

Compressed tar-zipped directory containing output of SDA computed on the LIBD discovery and GTEx replication datasets: individual score, tissue score, gene loading and posterior inclusion probability (PIP) matrices. The gene.loading_PIP_0.5.txt matrix refers to the gene loading matrix in which loadings for genes with PIP < .5 has been set to 0. This was the matrix used to infer component membership for each gene.

**Supplementary Data 2.**

Excel file with LIBD discovery components summary information, association with both biological covariates and technical confounders (Table S1), and component membership (Table S2).

Table S1: Association between all SDA components and both biological covariates and technical confounders (p-values are reported). The number of genes included in each component is listed, and the 69 components carried forward for further analysis are indicated (column name: removed; value = No).

Table S2: Information about genes assigned to the 69 SDA components. For each gene and for each component a value of 1 indicates that gene is assigned to that specific component.

**Supplementary Data 3.**

Excel file of discovery C80 (Table S1) and replication C18 (Table S2) genes as wells as relative GO and KEGG enrichment results (Table S3 and S4).

Table S1 and Table S2: List of genes belonging to discovery C80 and replication C18 components. Ensembl IDs are relative to GENCODE release 25.

Table S3 and Table S4: GO and KEGG results are reported for terms with p[FDR]<.05.

**Supplementary Data 4.**

Compressed tar-zipped directory containing compressed text files of:

- C80.g1000.gz: 1000 Genome SNPs mapped at 100kbp up- and down-stream of each C80 gene (A1 refers to reference allele)
- C80.g1000.PGC3.gz: PGC3 leave-LIBD-out SNPs matching the 1000 Genome SNPs mapped at C80 genes (A1 refers to effect allele)
- C80.PRS(1-10)_snpList.txt: LD-independent (after clumping by PRSice) PGC3 leave-LIBD-out SNPs used to compute C80-PRS1 to 10.

Chromosome position refers to human genome Build 37. Additionally, an RData object (C80.g1000_snpList.rds) containing the list of C80 genes with the corresponding 1000 Genome SNPs mapped is included in this compressed directory.

**Supplementary Data 5.**

Compressed tar-zipped directory containing scripts used for this study.

**Supplementary Data 6.**

Compressed tar-zipped directory containing LIBD and GTEx post-mortem gene expression data preprocessed for input to SDA.

#### Supplementary Figures

a

b

GWAS MAGMA H-MAGMA

c

**Supplementary Fig. 1: 69 SDA components characterization analyses results and discovery C80 enriched gene ontologies.**

**a**, Gene enrichment analysis results are shown for all 69 robust components. From the left, the first (GWAS), second (MAGMA) and third orange grids (H-MAGMA) show enrichment results for schizophrenia risk genes, other psychiatric illness risk genes, and immune condition risk genes. Enrichment testing results are shown for differentially expressed genes, differentially methylated genes, and loss of function variant intolerant genes in the green grid. The final lightblue grid show tissue specificity as determined by the tissue scores generated during the SDA process and reflects the relative contribution of component gene networks within each of the sampled regions to the overall component. Adjusted p-values shown are empirical p-values obtained from permutation tests. See Fig. 2 caption for abbreviations.

**b,** and **c,** Gene ontology enrichment of C80 for biological processes and molecular function. FDR-adjusted p-values are reported.

a

LIBD GTEx

GTEx LIBD

c

d

18

GNB1 AKT3 KCNJ3
GNAI2 ADCY5 MAPK9 PPP2R1B
DRD2 CACNA1C
GNG2 GNB5 PRKCB PPP1R1B
PRKCG PLCB1 PLCB4

17

b

**Supplementary Fig. 2: GTEx replication analyses results**.

**a,** 69 SDA components generated from the LIBD discovery dataset replicated in the GTEx dataset using JI (orange bars) or gene loading correlation (light blue bars). (*) indicates statistical significance after permutations test < .05; (**) indicates statistical significance after permutations test < .001. JI values on the right and gene loading $R^2$ on the left x axis are shown. Discovery C80 and replication C18 are one of the 4 pairs of components highlighted in red that are consistent with both replication measures.

**b**, Cell-type specificity of replication component C18 using human (left) and mouse (right) single-cell atlases. y-axes show FDR-adjusted p-values. Barplots demonstrates a higher specicifty for GABAergic, medium spiny and dopaminergic neurons. See Fig. 2 caption for abbreviations.

**c,** Gene ontology enrichment of replication component C18 for cellular compartments. FDR-adjusted p-values are reported.

**d,** Venn diagram showing intersection between C80 (orange) and C18 (green) genes enriched for KEGG term dopaminergic synapse with $p_{[FDR]}$<.05.
