## Supplementary material for "Dopamine and schizophrenia from bench to bedside: Discovery of a striatal co-expression risk gene set that predicts in vivo measures of striatal function": Compressed zipped directory containing scripts used for this study.: 1.sda_preprocessing.R

```
###############################################################################################################
############################################### SDA pre_processing ############################################
###############################################################################################################

rm(list = ls())
source("1a.sda_preprocessing_functions.R")

set.seed(123)
HP <- "hippo_brainseq_phase2_hg38_rseGene_merged_n447.rda"
CN <- "caudate_brainseq_phase3_hg38_rseGene_merged_n464.rda"
DLPFC <- "dlpfc_ribozero_brainseq_phase2_hg38_rseGene_merged_n453.rda"
caudate = convertRse(CAUDA,tissue = "CN")
dlpfc = convertRse(DLPFC,tissue = "DLPFC")
hippo = convertRse(HIPPO,tissue = "HP")


### intersection of genes ####
gene_common = geneSub(caudate,dlpfc,hippo)

#####  intersection of samples ####
sample_list = sampleSub(caudate,dlpfc,hippo)

##### matrix of common samples and genes ####
matrix_list = matrixSub(caudate,dlpfc,hippo)

##### median filtering
matrix_list = medianFilter(matrix_list)

##### update gene_common
gene_common = list(ALL = gene_common[match(colnames(matrix_list[[1]]$ALL),gene_common$gencodeID),],
                   CAUC = gene_common[match(colnames(matrix_list[[1]]$CAUC),gene_common$gencodeID),],
                   AA = gene_common[match(colnames(matrix_list[[1]]$AA),gene_common$gencodeID),])

### obtain final matrices to use as input for SDA ###
matrix_SDA = matrixSDA(matrix_list)

##### filter final matrices
gene_common = geneFilter(matrix_SDA,gene_common)
for (i in names(gene_common)){colnames(gene_common[[i]])[c(8,10)] = c("ensembl","hgnc")}
matrix_list = matrixFilter(matrix_SDA)

##### normalize final matrices
matrix_SDA_final = matrixSDAnorm(matrix_list)

### write .txt file to use as input
write.table(matrix_SDA_final$matrix_SDA_final$ALL,file = "./caud_dlpfc_hippo_ALL.txt",row.names = F,col.names = F,quote = F,sep= "\t")

### save 
save(caudate,dlpfc,hippo,matrix_list,sample_list,matrix_SDA,matrix_SDA_final,gene_common, file = paste0("./sda_preprocessing.RData"))

#### RUN SDA on cluster ####
```
