## Supplementary material for "Dopamine and schizophrenia from bench to bedside: Discovery of a striatal co-expression risk gene set that predicts in vivo measures of striatal function": Compressed zipped directory containing scripts used for this study.: 1a.sda_preprocessing_functions.R

```
###############################################################################################################
############################################### SDA pre_processing ############################################
###############################################################################################################

###### preprocessing functions #####
library(SummarizedExperiment)
library("recount")
library("jaffelab")
library(readr)
library (plyr)
library(car)
library(dplyr)
library(preprocessCore)
library(RNOmni)

convertRse <- function(filename, minAge = -1, maxAge = 150, minRin = 6,tissue) {
          if(tissue == "DENTATE"){
                    rseName <- paste0(substring(filename, 1, 5), "rse")
                    load(filename)
                    rse_merged = rse_gene
                    filtered = rse_merged[,rse_merged$Age >= minAge &
                                                    rse_merged$Age <= maxAge &
                                                    rse_merged$Race %in% c('AA', 'CAUC') &
                                                    rse_merged$Dx %in% c("Control")]
                    expMat <- getRPKM(filtered, length_var = "Length", mapped_var = "numMapped")
                    indVar = data.frame(colData(filtered))
                    map = data.frame(rowRanges(filtered)) # this is not filtered by exp
                    map <- map[which(map$gencodeID %in% row.names(expMat)),]
                    stopifnot(identical(map$gencodeID, row.names(expMat)))
                    row.names(map) = row.names(expMat)
                    return(list(expMat, indVar, map))
          } else {
                    rseName <- paste0(substring(filename, 1, 5), "rse")
                    load(filename)
                    rse_merged <- merge_rse_metrics(rse_gene)
                    filtered = rse_merged[,rse_merged$Age >= minAge &
                                                    rse_merged$Age <= maxAge &
                                                    rse_merged$Race %in% c('AA', 'CAUC') &
                                                    rse_merged$Dx %in% c("Control") &
                                                    sapply(rse_merged$RIN,mean) >= minRin]
                    expMat <- getRPKM(filtered, length_var = "Length", mapped_var = "numMapped")
                    indVar = data.frame(colData(filtered))
                    map = data.frame(rowRanges(filtered)) # this is not filtered by exp
                    map <- map[which(map$gencodeID %in% row.names(expMat)),]
                    stopifnot(identical(map$gencodeID, row.names(expMat)))
                    row.names(map) = row.names(expMat)
                    return(list(expMat, indVar, map))
          }
}

geneSub = function(caudate,dlpfc,hippo){
          
          gene_caud = row.names(caudate[[1]])
          gene_dlpfc = row.names(dlpfc[[1]])
          gene_hippo = row.names(hippo[[1]])
          
          gene_common = unique(Reduce(intersect,list(gene_caud,gene_dlpfc,gene_hippo)))
          gene_common = caudate[[3]][which(caudate[[3]]$gencodeID %in% gene_common),1:14]
          #### removal of Mitochondrial genes 
          mitho = gene_common$gencodeID[grep("MT-",gene_common$Symbol)]
          gene_common = gene_common[!(gene_common$gencodeID %in% mitho),]
          
          return(gene_common)
} ### function to obtain common genes between tissues

sampleSub = function(caudate,dlpfc,hippo) {
          
          sample_caud = caudate[[2]][order(caudate[[2]]$BrNum),c("BrNum","RNum","Region","RIN","Age","Sex","Race","Dx","pH","PMI","mitoRate","totalAssignedGene","rRNA_rate")]
          sample_dlpfc = dlpfc[[2]][order(dlpfc[[2]]$BrNum),c("BrNum","RNum","Region","RIN","Age","Sex","Race","Dx","mitoRate","totalAssignedGene","rRNA_rate")]
          sample_hippo = hippo[[2]][order(hippo[[2]]$BrNum),c("BrNum","RNum","Region","RIN","Age","Sex","Race","Dx","mitoRate","totalAssignedGene","rRNA_rate")]
          sample_dlpfc$RIN = as.numeric(sample_dlpfc$RIN)
          sample_hippo$RIN = as.numeric(sample_hippo$RIN)
          
          sample_common = Reduce(intersect,list(sample_caud$BrNum,sample_dlpfc$BrNum,sample_hippo$BrNum))
          #### outliers removal
          load("C:/Users/lsporte1/Desktop/sda/script_3D/outliers_sample_removed_BrNum.RData")
          outliers = as.character(unlist(outlier_samples_removed[1:3]))
          samp_out = intersect(sample_common,outliers)
          sample_common = sample_common[!(sample_common %in% outliers)]
          ####
          sample_common = sort(sample_common)
          
          sample_caud = sample_caud[which(sample_caud$BrNum %in% sample_common),]
          sample_dlpfc = sample_dlpfc[which(sample_dlpfc$BrNum %in% sample_common),]
          sample_hippo = sample_hippo[which(sample_hippo$BrNum %in% sample_common),]
          
          if(identical(sample_dlpfc$BrNum,sample_caud$BrNum) & identical(sample_caud$BrNum,sample_hippo$BrNum)){
                    
                    
                    #### order by Dx and Age ####
                    sample_caud = sample_caud[order(sample_caud$Dx,sample_caud$Age),]
                    
                    sample_dlpfc = sample_dlpfc[order(sample_dlpfc$Dx,sample_dlpfc$Age),]
                    
                    sample_hippo = sample_hippo[order(sample_hippo$Dx,sample_hippo$Age),]
                    
                    ##### subset by Race ####
                    
                    sample_caud_CAUC = sample_caud[which(sample_caud$Race == "CAUC"),]
                    sample_caud_AA = sample_caud[which(sample_caud$Race == "AA"),]
                    
                    sample_dlpfc_CAUC = sample_dlpfc[which(sample_dlpfc$Race == "CAUC"),]
                    sample_dlpfc_AA = sample_dlpfc[which(sample_dlpfc$Race == "AA"),]
                    
                    sample_hippo_CAUC = sample_hippo[which(sample_hippo$Race == "CAUC"),]
                    sample_hippo_AA = sample_hippo[which(sample_hippo$Race == "AA"),]
                    
                    sample_list = list(caudate = list(ALL = sample_caud,CAUC = sample_caud_CAUC,AA = sample_caud_AA),
                                       dlpfc = list(ALL = sample_dlpfc,CAUC = sample_dlpfc_CAUC,AA = sample_dlpfc_AA),
                                       hippo = list(ALL = sample_hippo,CAUC = sample_hippo_CAUC,AA = sample_hippo_AA))
                    
                    
          } else stop("samples are not identical")
          
          return(sample_list)
} ### function to obtain common samples between tissues

matrixSub = function(caudate,dlpfc,hippo){
          
          matrix_list = list(caudate = list() ,dlpfc = list() ,hippo = list() )
          
          
          for (tissue in c("caudate","dlpfc","hippo")) { 
                    
                    for (race in c("ALL","CAUC","AA")) {
                              
                              
                              df = get(tissue)[[1]][which(row.names(get(tissue)[[1]]) %in% gene_common$gencodeID),
                                                    which(colnames(get(tissue)[[1]]) %in% row.names(sample_list[[tissue]][[race]]))]
                              
                              matrix_list[[tissue]][[race]] = t(df[,order(match(colnames(df),
                                                                                row.names(sample_list[[tissue]][[race]])))])
                              
                    }
          }
          
          if (identical(row.names(matrix_list$caudate$ALL),row.names(sample_list$caudate$ALL)) &
              identical(row.names(matrix_list$dlpfc$ALL),row.names(sample_list$dlpfc$ALL)) &
              identical(row.names(matrix_list$hippo$ALL),row.names(sample_list$hippo$ALL))) 
                    
          {return(matrix_list)}
          
          else stop("samples are not identical")
} ### function to obtain common samples and genes matrices

medianFilter = function(matrix_list){
          
          median_fltered = lapply(matrix_list,function(x){
                    lapply(x,function(y){
                              new = apply(y,2,median)
                              new = names(new[which(new < .1)])
                              matrix = y[,-which(colnames(y) %in% new)]
                              new_m = colnames(matrix)[which(abs(scale(apply(matrix,2,median))) > 3)]
                              return(union(new,new_m))
                    })                  
          })
          out_genes_ALL = Reduce(intersect,list(median_fltered$caudate$ALL,median_fltered$dlpfc$ALL,median_fltered$hippo$ALL))
          out_genes_CAUC = Reduce(intersect,list(median_fltered$caudate$CAUC,median_fltered$dlpfc$CAUC,median_fltered$hippo$CAUC))
          out_genes_AA = Reduce(intersect,list(median_fltered$caudate$AA,median_fltered$dlpfc$AA,median_fltered$hippo$AA))
          for(m in names(matrix_list)){
                    matrix_list[[m]]$ALL = matrix_list[[m]]$ALL[,-which(colnames(matrix_list[[m]]$ALL) %in% out_genes_ALL)]
                    matrix_list[[m]]$CAUC = matrix_list[[m]]$CAUC[,-which(colnames(matrix_list[[m]]$CAUC) %in% out_genes_CAUC)]
                    matrix_list[[m]]$AA = matrix_list[[m]]$AA[,-which(colnames(matrix_list[[m]]$AA) %in% out_genes_AA)]
          }
          return(matrix_list)
} #### function to filter common genes with median < 0.1 in each tissue

matrixSDA = function(matrix_list,check_dim = TRUE) {
          
          matrix_ALL = rbind(matrix_list$caudate$ALL,matrix_list$dlpfc$ALL,matrix_list$hippo$ALL)
          matrix_CAUC = rbind(matrix_list$caudate$CAUC,matrix_list$dlpfc$CAUC,matrix_list$hippo$CAUC)
          matrix_AA = rbind(matrix_list$caudate$AA,matrix_list$dlpfc$AA,matrix_list$hippo$AA)
          
          if (check_dim){
                    if (dim(matrix_ALL)[[2]] == dim(matrix_CAUC)[[2]] & dim(matrix_ALL)[[2]] == dim(matrix_AA)[[2]] &
                        dim(matrix_ALL)[[1]] == (dim(matrix_CAUC)[[1]] + dim(matrix_AA)[[1]])){
                              
                              row.names(matrix_ALL) = rep(sample_list$caudate$ALL$BrNum,3)
                              row.names(matrix_CAUC) = rep(sample_list$caudate$CAUC$BrNum,3)
                              row.names(matrix_AA) = rep(sample_list$caudate$AA$BrNum,3)
                              
                              colnames(matrix_ALL) = caudate[[3]][which(caudate[[3]]$gencodeID %in% gene_common$gencodeID),]$ensemblID
                              colnames(matrix_CAUC) = caudate[[3]][which(caudate[[3]]$gencodeID %in% gene_common$gencodeID),]$ensemblID
                              colnames(matrix_AA) = caudate[[3]][which(caudate[[3]]$gencodeID %in% gene_common$gencodeID),]$ensemblID
                              
                              return(list(ALL = matrix_ALL,CAUC = matrix_CAUC,AA = matrix_AA)) 
                    } else stop("matrices don't have correct dimensions")}
          
          else if (!check_dim){
                    row.names(matrix_ALL) = rep(sample_list$caudate$ALL$BrNum,3)
                    row.names(matrix_CAUC) = rep(sample_list$caudate$CAUC$BrNum,3)
                    row.names(matrix_AA) = rep(sample_list$caudate$AA$BrNum,3)
                    
                    colnames(matrix_ALL) = caudate[[3]][which(caudate[[3]]$gencodeID %in% gene_common[["ALL"]]$gencodeID),]$ensemblID
                    colnames(matrix_CAUC) = caudate[[3]][which(caudate[[3]]$gencodeID %in% gene_common[["CAUC"]]$gencodeID),]$ensemblID
                    colnames(matrix_AA) = caudate[[3]][which(caudate[[3]]$gencodeID %in% gene_common[["AA"]]$gencodeID),]$ensemblID
                    
                    return(list(ALL = matrix_ALL,CAUC = matrix_CAUC,AA = matrix_AA))}
} ### function to obtain input SDA matrices

geneFilter = function(matrix_SDA,gene_common){
          
          #### removal of genes with > 20% of 0 in ALL tissues
        
          for (pop in names(matrix_SDA)){
                    
                    th = as.integer((nrow(matrix_SDA[[pop]])*20)/100)
                    
                    genes_th = c()
                    for (i in colnames(matrix_SDA[[pop]])){
                              
                              if (length(matrix_SDA[[pop]][which(matrix_SDA[[pop]][,i] == 0),i]) > th) genes_th = c(genes_th,i) else {next}
                              
                    }
                    
                    gene_common[[pop]] = gene_common[[pop]][!(gene_common[[pop]]$ensemblID %in% genes_th),]
          }
          return(gene_common)
          
} ### function to filter out genes with > 20% of 0s in ALL 3 tissue

matrixFilter = function(matrix_SDA){
          
          for (pop in names(matrix_SDA)){
                    
                    for (tissue in names(matrix_list)){
                              matrix_list[[tissue]][[pop]]=matrix_list[[tissue]][[pop]][,gene_common[[pop]]$gencodeID]}
          }
          return(matrix_list)
 
} ### function to filter matrices according to new filtered genes

matrixSDAnorm = function(matrix_list){
          
          
          #### log2 transformation
          matrix_list_log = lapply(matrix_list,function(x)
                    lapply(x,function(y)
                              log2(y + 1)))
          #### quantile normalization in each tissue
          matrix_list_quantile = lapply(matrix_list_log,function(x)
          lapply(x,function(y)
          t(normalize.quantiles(t(y)))))
          #### rank transformation (Blom)
          matrix_list_blom = lapply(matrix_list_quantile,function(x)
                    lapply(x,function(y)
                    apply(y,2,rankNorm)))
          matrix_SDA_norm <- matrixSDA(matrix_list_blom,check_dim = F)
          
          return(list(matrix_log2 = matrix_list_log,
                      matrix_quantile = matrix_list_quantile,
                      matrix_Blom = matrix_list_blom,
                      matrix_SDA_final = matrix_SDA_norm))
          
          
} ### function to quantile and rank normalize in each tissue
```
