## Supplementary material for "Dopamine and schizophrenia from bench to bedside: Discovery of a striatal co-expression risk gene set that predicts in vivo measures of striatal function": Compressed zipped directory containing scripts used for this study.: 2a.sda_postProcess.R

```
###############################################################################################################
############################################### SDA post_processing ############################################
###############################################################################################################

postProcess = function(data_name,
                       samples,
                       tissues,
                       genes,
                       seed = NULL,
                       threshold,
                       data_dir)
{
    source("2b.sda_postprocessing_functions.R")         
  #browser()
  #### reading SDA output ####
  setwd(paste0(data_dir)) ### directory where output files are
  
  output_SDA = read_output_SDA(file_name = data_name)
  
  #### reset every matrix according to non-missing-values components
  output_SDA_subset = sapply(1:length(output_SDA), function(x){
    if(length(tissues) > 1){
      colnames(output_SDA[[x]]$A) = 
        colnames(output_SDA[[x]]$B) = 
        row.names(output_SDA[[x]]$X) = 
        row.names(output_SDA[[x]]$S) = 
        paste0(colnames(output_SDA[[x]]$A),".",x)
    } else {
      colnames(output_SDA[[x]]$A) = 
        row.names(output_SDA[[x]]$X) = 
        row.names(output_SDA[[x]]$S) = 
        paste0(colnames(output_SDA[[x]]$A),".",x)
    }
      output_SDA[[x]]$A = output_SDA[[x]]$A[,!apply(output_SDA[[x]]$A,2,function(y) all(y == 0))]
      output_SDA[[x]]
    },simplify = F)
  names(output_SDA_subset) = names(output_SDA)
  
  ##### clustering components across 10 runs using 1-|cor| as dissimilarity measure and 0.4,0.5 or 0.6 as clustering threshold ####
  abs.corr.matrix = abscorrmatrix(output_SDA_subset = output_SDA_subset)
  if(threshold == 0.4) {th = 0.6} else if (threshold == 0.6) {th = 0.4} else th = threshold
  components.clusters = hclust(as.dist(1-abs.corr.matrix))
  pdf(paste0("SDA_cluster_dendogram_",th,"corr.pdf"))
  plot(components.clusters)
  rect.hclust(components.clusters,h = threshold)
  dev.off()
  
  #### obtaining final SDA matrices with robust components ####
  if(length(tissues) > 1){
    final_SDA = clusterCombine(clusters = components.clusters,
                               output_SDA_subset = output_SDA_subset,
                               th = threshold,file_name = data_name,
                               nsubj = samples,ntissue = length(tissues),ngene = genes)
  } else final_SDA = clusterCombine_2D(clusters = components.clusters,
                                       output_SDA_subset = output_SDA_subset,
                                       th = threshold,file_name = data_name,
                                       nsubj = samples,ntissue = length(tissues),ngene = genes)
  }
  
rename = function(SDA_out,
                  samples,
                  tissues,
                  seed = NULL,
                  population = NULL,
                  preProcess_dir,
                  threshold,
                  output_dir)
{
  ### renaming samples and genes of each matrix
  load(paste0(preProcess_dir,"/sda_preprocessing.RData"))
  #browser()
  data = matrix_SDA_final$matrix_log2$caudate$all
  row.names(final_SDA$A) = row.names(data)
  if(!is.null(tissues)){ row.names(final_SDA$B) = tissues }
  colnames(final_SDA$X) = limma::strsplit2(colnames(data),"\\.")[,1]
  colnames(final_SDA$S) = limma::strsplit2(colnames(data),"\\.")[,1]
  
  ### obtaining network of genes with different PIP threshold for each component
  final_SDA[["X_sig_0.5"]] = (final_SDA$X * as.matrix(final_SDA$S > 0.5))     
  final_SDA[["X_sig_1"]] = (final_SDA$X * as.matrix(final_SDA$S == 1 ))
  colnames(final_SDA$X_sig_0.5) = colnames(final_SDA$X_sig_1) = colnames(final_SDA$X)
  
  return(final_SDA)
}
```
