## Supplementary material for "Dopamine and schizophrenia from bench to bedside: Discovery of a striatal co-expression risk gene set that predicts in vivo measures of striatal function": Compressed zipped directory containing scripts used for this study.: 3.sda_metadata.R

```
###############################################################################################################
######################## SDA metadata  ####################################
###############################################################################################################

load("sda_preprocessing.RData")
rm(list = setdiff(ls(),"sample_list"))
source("3a.sda_metadata_functions.R")

data = metadata_LIBD(data_dir = "",
              tissues = c("caudate","dlpfc","hippo"),common_metadata = c("Dx","Race","Sex","Age"),
              tissue_specific_metadata = c("RIN","totalAssignedGene","rRNA_rate","mitoRate"),
              data_name = "final_SDA",output_dir = "",
              sample_list = sample_list,pop = "ALL")

save(data,file = paste0("data.SDA.RData"))
```
