## Supplementary material for "Dopamine and schizophrenia from bench to bedside: Discovery of a striatal co-expression risk gene set that predicts in vivo measures of striatal function": Compressed zipped directory containing scripts used for this study.: 3a.sda_metadata_functions.R

```
###############################################################################################################
######################## SDA metadata  ####################################
###############################################################################################################

metadata_LIBD = function(data_dir,data_name,
                         tissues,pop,
                         common_metadata,
                         tissue_specific_metadata,
                         sample_list,
                         output_dir)
{
          
          #browser()
          require(dplyr)
          load(paste0(data_dir,data_name,".RData"))
          sample_list = lapply(sample_list,function(x)
          {
                    x[[pop]]$RIN = as.numeric(x[[pop]]$RIN)
                    x
                    
          })
          metadata <- sample_list[[1]][[pop]][,c("BrNum",common_metadata)]
          for (i in c("Dx","Race","Sex")){metadata[[i]] = as.factor(metadata[[i]])}
          ##### technical confounders
          batch_data = lapply(tissues,t=tissue_specific_metadata,function(x,t)
          {
                    new = sample_list[[x]][[pop]][,t]
                    colnames(new) = paste0(t,"_",x)
                    new
                    
          })
          batch_data = as.data.frame(batch_data)
          batch_data = batch_data[match(row.names(metadata),row.names(batch_data)),]
          ##### Genomic eigenvariates
          load("Genomic_eigenvariates.RData")
          GE = Genomic_eigenvariates[match(metadata$BrNum,row.names(Genomic_eigenvariates)),]
          ##### PRS6 for PGC3
          load("PRS.PGC3.RData")
          PRS.PGC3 = PRS_CAUDATE.DLPFC.HIPPO.DG[match(metadata$BrNum,row.names(PRS_CAUDATE.DLPFC.HIPPO.DG)),c("PRS_6")]
          colnames(PRS.PGC3) = c("PRS6.PGC3")
          ##### SDA components
          final_SDA = final_SDA$A[match(metadata$BrNum,row.names(final_SDA$A)),]
          ##### combine data
          data = cbind(metadata,batch_data,GE,PRS.PGC2,PRS.PGC3,final_SDA)
          
}
```
