## Supplementary material for "Dopamine and schizophrenia from bench to bedside: Discovery of a striatal co-expression risk gene set that predicts in vivo measures of striatal function": Compressed zipped directory containing scripts used for this study.: 4.sda_covariates.R

```
###############################################################################################################
######################## SDA component association with covariates ########################
###############################################################################################################

rm(list = ls())
LMdetect <- function(myComp, myVar, myData, value) {
  
  LMmod <- as.formula(paste0(myComp,"~", paste0(myVar, collapse = "+")))
  if(value == "p")results <- summary(lm(LMmod, data = myData))$coefficients[-1,4]
  if(value == "t")results <- summary(lm(LMmod, data = myData))$coefficients[-1,3]
  return(results)
}

load(paste0("data.SDA.RData"))

### correlation analysis
myVar = c("Age","Sex", "Dx","PMI",
          outer(c("RIN","totalAssignedGene","mitoRate","rRNA_rate"),c("_caudate","_dlpfc","_hippo"),FUN = "paste0"),
          paste0("GE_",1:10))
myData = data
myComps = grep("^V",colnames(myData),value = T)
test_result_p = t(sapply(myComps,function(x)LMdetect(x,myVar,myData,value = "p")))
test_result_t = t(sapply(myComps,function(x)LMdetect(x,myVar,myData,value = "t")))

save(test_result_t,test_result_p, file = "confounders.RData")
```
