## Supplementary material for "Dopamine and schizophrenia from bench to bedside: Discovery of a striatal co-expression risk gene set that predicts in vivo measures of striatal function": Compressed zipped directory containing scripts used for this study.: 5.sda_components_pruning.R

```
###############################################################################################################
######################## SDA components pruning ##################################
###############################################################################################################

load(paste0("confounders.RData"))
load(paste0("final_SDA.RData"))

technical_association = test_result_p[,-grep("Age|Sex|Dx",colnames(test_result_p))]
technical_threshold = technical_association < 5.1e-04
technical_threshold = apply(technical_threshold,1,any)
index.out = names(technical_threshold[technical_threshold])
index.filtered = setdiff(row.names(test_result_p),index.out)
##########################################  

rm(list = setdiff(ls(),c("index.filtered","index.out","final_SDA")))

final_SDA_filtered = final_SDA
final_SDA_filtered$A = final_SDA$A[,colnames(final_SDA$A) %in% index.filtered]
final_SDA_filtered$B = final_SDA$B[,colnames(final_SDA$B) %in% index.filtered]
final_SDA_filtered$X = final_SDA$X[row.names(final_SDA$X) %in% index.filtered,]
final_SDA_filtered$S = final_SDA$S[row.names(final_SDA$S) %in% index.filtered,]
final_SDA_filtered$X_sig_0.5 = final_SDA$X_sig_0.5[row.names(final_SDA$X_sig_0.5) %in% index.filtered,]
final_SDA_filtered$X_sig_1 = final_SDA$X_sig_1[row.names(final_SDA$X_sig_1) %in% index.filtered,]
colnames(final_SDA_filtered$X_sig_0.5) = colnames(final_SDA_filtered$X_sig_1) = colnames(final_SDA_filtered$X)

final_SDA_out = final_SDA
final_SDA_out$A = final_SDA$A[,which(colnames(final_SDA$A) %in% index.out)]
final_SDA_out$B = final_SDA$B[,which(colnames(final_SDA$B) %in% index.out)]
final_SDA_out$X = final_SDA$X[which(row.names(final_SDA$X) %in% index.out),]
final_SDA_out$S = final_SDA$S[which(row.names(final_SDA$S) %in% index.out),]
final_SDA_out$X_sig_0.5 = final_SDA$X_sig_0.5[which(row.names(final_SDA$X_sig_0.5) %in% index.out),]
final_SDA_out$X_sig_1 = final_SDA$X_sig_1[which(row.names(final_SDA$X_sig_1) %in% index.out),]
colnames(final_SDA_out$X_sig_0.5) = colnames(final_SDA_out$X_sig_1) = colnames(final_SDA_out$X)

save.image(file = paste0("final_SDA.RData"))
```
