## Supplementary material for "Dopamine and schizophrenia from bench to bedside: Discovery of a striatal co-expression risk gene set that predicts in vivo measures of striatal function": Compressed zipped directory containing scripts used for this study.: 8.Dx_PRS_association.R

```
###############################################################################################################
######################## SDA components association with Dx and PRS ##################################
###############################################################################################################

###### stats to detect PRS and Dx effects #####
LMdetect <- function(myComp, myData, myVar, myMod,summary = F) {
  
  myName <- tissue.score.c[myComp,]   
  active <- colnames(tissue.score.c)[ which(myName==1) ]
  mod = c(sapply(active,function(x){grep(x, names(myData), value = T)}),myMod)
  LMmod <- as.formula(paste0(myComp,"~", paste0(mod, collapse = "+")))
  fit <- lm(LMmod, data = myData)  
  print(summary(fit))
  # lmp <- function (modelobject) {
  #           if (class(modelobject) != "lm") stop("Not an object of class 'lm' ")
  #           f <- summary(modelobject)$fstatistic
  #           p <- pf(f[1],f[2],f[3],lower.tail=F)
  #           attributes(p) <- NULL
  #           return(p)
  # }
  if(summary){   results = fit      } else results = summary(fit)$coefficients[myVar,3:4]
  
}

###### load and set data #####
load("data.SDA.RData")
load("tissue.score.RData")

data = data[!data$BrNum %in% data$BrNum[is.na(data$GE_1)],]
to_filter_out = c(grep("^PC",colnames(data),value = T))
data = data[,!colnames(data)%in% to_filter_out]
adults <- data[data$Age > 17,]
row.names(adults) = adults$BrNum
adults$BrNum <- NULL
comps <- row.names(active_components_filtered)

#### Dx association ####
testVars <- c(grep(paste0("Age|Sex|PMI"),names(adults),value = T),"Dx")
Dx_assoc = sapply(comps,function(x)LMdetect(myComp = x,
                                            myData = adults,
                                            myVar = "DxSchizo",
                                            myMod = testVars,summary = T),simplify = F)
Dx_t = sapply(Dx_assoc,function(x) summary(x)$coefficients["DxSchizo",3])
Dx_anova = sapply(Dx_assoc, function(x) anova(x),simplify = F)
Dx_p = sapply(Dx_anova,function(x) x["Dx",5])

#### PRS association ####
adults_CAUC = adults[adults$Race %in% "CAUC",]
adults_CAUC[,paste0("PRS6.PGC3")] = PRS_CAUDATE.DLPFC.HIPPO.DG[match(row.names(adults_CAUC),row.names(PRS_CAUDATE.DLPFC.HIPPO.DG)),
                                                                      1:10]

PRS_assoc = sapply(comps,function(x)
  {
    
    testVars <- grep(paste0("Dx|Age|Sex|PMI|PRS6.PGC3"),
                     names(adults_CAUC),value = T)
    LMdetect(myComp = x,
             myData = adults_CAUC,
             myVar = prs,
             myMod = testVars,summary = T) 
  },simplify = F)

PRS_t = t(sapply(PRS_assoc,function(x) sapply(x,function(y) summary(y)$coefficients["PRS6.PGC3",3]  )))
PRS_p = t(sapply(PRS_assoc,function(x) sapply(x,function(y) summary(y)$coefficients["PRS6.PGC3",4]  )))

save(Dx_anova, PRS_assoc, file = "Dx.PRS_assoc.RData")
```
