## Supplementary material for "Dopamine and schizophrenia from bench to bedside: Discovery of a striatal co-expression risk gene set that predicts in vivo measures of striatal function": Compressed zipped directory containing scripts used for this study.: 1.sda_preprocessing.R

```
###############################################################################################################
############################################### SDA pre_processing ############################################
###############################################################################################################

rm(list = ls())
source("1a.sda_preprocessing_functions.R")

set.seed(123)
load("gtex_CAUD_DLPFC_HIPPO.RData")
gtex_CAUDATE_DLPFC_HIPPO = lapply(gtex_CAUDATE_DLPFC_HIPPO,function(x) x[c(1,3,2)])

#### RIN filtering 6 ####
gtex_CAUDATE_DLPFC_HIPPO = lapply(names(gtex_CAUDATE_DLPFC_HIPPO),function(x)
{
        gtex_CAUDATE_DLPFC_HIPPO[[x]][[2]] =
                  gtex_CAUDATE_DLPFC_HIPPO[[x]][[2]][gtex_CAUDATE_DLPFC_HIPPO[[x]][[2]]$RIN >= 6 ,]
        gtex_CAUDATE_DLPFC_HIPPO[[x]]
})
names(gtex_CAUDATE_DLPFC_HIPPO) = c("CAUDATE","DLPFC","HIPPO")

### intersection of genes ####
gene_common = geneSub(gtex_CAUDATE_DLPFC_HIPPO$CAUDATE,
                      gtex_CAUDATE_DLPFC_HIPPO$DLPFC,
                      gtex_CAUDATE_DLPFC_HIPPO$HIPPO)

#####  intersection of samples ####
sample_list = sampleSub(gtex_CAUDATE_DLPFC_HIPPO$CAUDATE,
                        gtex_CAUDATE_DLPFC_HIPPO$DLPFC,
                        gtex_CAUDATE_DLPFC_HIPPO$HIPPO)

##### matrix of common samples and genes ####
matrix_list = matrixSub(gtex_CAUDATE_DLPFC_HIPPO$CAUDATE,
                        gtex_CAUDATE_DLPFC_HIPPO$DLPFC,
                        gtex_CAUDATE_DLPFC_HIPPO$HIPPO)

##### median filtering
matrix_list = medianFilter(matrix_list)

##### update gene_common
gene_common = gene_common[match(colnames(matrix_list$caudate),gene_common$gencodeID),]

### obtain final matrices to use as input for SDA ###
matrix_SDA = matrixSDA(matrix_list)

##### filter final matrices
gene_common = geneFilter(matrix_SDA,gene_common)
colnames(gene_common)[c(8,10)] = c("ensembl","hgnc")
matrix_list = matrixFilter(matrix_list)

##### normalize final matrices
matrix_SDA_final = matrixSDAnorm(matrix_list)

### write .txt file to use as input
write.table(matrix_SDA_final$matrix_SDA_final,file = "caudate_dlpfc_hippo_GTEx.txt",row.names = F,col.names = F,quote = F,sep= "\t")

### save 
save(gtex_CAUDATE_DLPFC_HIPPO,matrix_list,sample_list,matrix_SDA,matrix_SDA_final,gene_common, file =paste0("sda_preprocessing.RData"))

#### RUN SDA on cluster ####
```
