## Supplementary material for "Dopamine and schizophrenia from bench to bedside: Discovery of a striatal co-expression risk gene set that predicts in vivo measures of striatal function": Compressed zipped directory containing scripts used for this study.: 1a.sda_preprocessing_functions.R

```
###############################################################################################################
############################################### SDA pre_processing ############################################
###############################################################################################################

###### preprocessing functions #####
require(readr)
require (plyr)
require(car)
require(dplyr)
require(preprocessCore)
require(RNOmni)

geneSub = function(caudate,dlpfc,hippo){
          
          gene_caud = row.names(caudate[[1]])
          gene_dlpfc = row.names(dlpfc[[1]])
          gene_hippo = row.names(hippo[[1]])
          
          gene_common = unique(Reduce(intersect,list(gene_caud,gene_dlpfc,gene_hippo)))
          gene_common = caudate[[3]][which(caudate[[3]]$gencodeID %in% gene_common),1:14]
          #### removal of Mitochondrial genes 
          mitho = gene_common$gencodeID[grep("MT-",gene_common$Symbol)]
          gene_common = gene_common[!(gene_common$gencodeID %in% mitho),]
          
          return(gene_common)
} ### function to obtain common genes between tissues

sampleSub = function(caudate,dlpfc,hippo) {
          
          sample_common = Reduce(intersect,list(caudate[[2]]$SAMPID,dlpfc[[2]]$SAMPID,hippo[[2]]$SAMPID))
          sample_common = sort(sample_common)
          sample_caud = caudate[[2]][caudate[[2]]$SAMPID %in% sample_common,]
          sample_dlpfc = dlpfc[[2]][dlpfc[[2]]$SAMPID %in% sample_common,]
          sample_hippo = hippo[[2]][hippo[[2]]$SAMPID %in% sample_common,]
          sample_caud=sample_caud[order(sample_caud$SAMPID),];sample_dlpfc=sample_dlpfc[order(sample_dlpfc$SAMPID),];sample_hippo=sample_hippo[order(sample_hippo$SAMPID),]
          
          stopifnot(all(sapply(list(sample_dlpfc$SAMPID,sample_hippo$SAMPID), FUN = identical, sample_caud$SAMPID)))
          sample_list = list(caudate = sample_caud,
                             dlpfc = sample_dlpfc,
                             hippo = sample_hippo)
          return(sample_list)
} ### function to obtain common samples between tissues

matrixSub = function(caudate,dlpfc,hippo){
          
          #browser()
          matrix_list = list(caudate = matrix() ,dlpfc = matrix(),hippo = matrix())
          
          
          for (tissue in c("caudate","dlpfc","hippo")) { 
                    
                    matrix_list[[tissue]] = t(get(tissue)[[1]][match(gene_common$gencodeID,row.names(get(tissue)[[1]])),
                                          match(row.names(sample_list[[tissue]]),colnames(get(tissue)[[1]]))])
                    
                    # matrix_list[[tissue]] = t(df[,order(match(colnames(df),
                    #                                                   row.names(sample_list[[tissue]])))])
                    #stopifnot(identical(row.names(sample_list[[tissue]]),)))
                    row.names(matrix_list[[tissue]]) = sample_list[[tissue]]$SAMPID[match(row.names(matrix_list[[tissue]]),
                                                                                          row.names(sample_list[[tissue]]))]
                    
                    
          }
          stopifnot(all(sapply(list(sample_list$caudate$SAMPID,
                                    sample_list$dlpfc$SAMPID,
                                    sample_list$hippo$SAMPID,
                                    row.names(matrix_list$caudate),
                                    row.names(matrix_list$dlpfc)),identical,row.names(matrix_list$hippo))))
          return(matrix_list)
          
          
} ### function to obtain common samples and genes matrices

medianFilter = function(matrix_list){
          
          median_fltered = lapply(matrix_list,function(x){
                    
                    new = apply(x,2,median)
                    new = names(new[which(new < .1)])
                    matrix = x[,!colnames(x) %in% new]
                    new_m = colnames(matrix)[abs(scale(apply(matrix,2,median))) > 3]
                    return(union(new,new_m))
                    
          })
          out_genes = Reduce(intersect,list(median_fltered$caudate,median_fltered$dlpfc,median_fltered$hippo))
          matrix_list = lapply(matrix_list,function(m) m[,!colnames(m) %in% out_genes])
          return(matrix_list)
          
} #### function to filter common genes with median < 0.1 in each tissue

matrixSDA = function(matrix_list) {
          
          matrix = rbind(matrix_list$caudate,matrix_list$dlpfc,matrix_list$hippo)
          colnames(matrix) = gene_common$ensemblID[match(colnames(matrix),gene_common$gencodeID)]
          
          
          return(matrix)
} ### function to obtain input SDA matrices

geneFilter = function(matrix_SDA,gene_common){
          
          #### removal of genes with > 20% of 0 in ALL tissues
          th = as.integer((nrow(matrix_SDA)*20)/100)
          
          genes_th = sapply(colnames(matrix_SDA),function(x){sum(matrix_SDA[,x]==0)})
          genes_th = names(genes_th[genes_th >= th])
          
          gene_common = gene_common[!gene_common$ensemblID %in% genes_th,]
          
          return(gene_common)
          
} ### function to filter out genes with > 20% of 0s in ALL 3 tissue

matrixFilter = function(matrix_list){
          
          m = lapply(matrix_list,function(x){x[,gene_common$gencodeID]})
                    
          
          return(m)
          
} ### function to filter matrices according to new filtered genes

matrixSDAnorm = function(matrix_list){
          
          
          #### log2 transformation
          matrix_list_log = lapply(matrix_list,function(x)
                    log2(x + 1))
          #### quantile normalization in each tissue
          matrix_list_quantile = lapply(matrix_list_log,function(x)
                    t(normalize.quantiles(t(x))))
          #### rank transformation (Blom)
          matrix_list_blom = lapply(matrix_list_quantile,function(x)
                    apply(x,2,rankNorm))
          matrix_SDA_norm <- rbind(matrix_list_blom$caudate,
                                   matrix_list_blom$dlpfc,matrix_list_blom$hippo)
          row.names(matrix_SDA_norm) = rep(sample_list$caudate$SAMPID,3)
          colnames(matrix_SDA_norm) = gene_common$ensembl
          
          return(list(matrix_log2 = matrix_list_log,
                      matrix_quantile = matrix_list_quantile,
                      matrix_Blom = matrix_list_blom,
                      matrix_SDA_final = matrix_SDA_norm))
          
          
} ### function to quantile and rank normalize in each tissue
```
