## Supplementary material for "Dopamine and schizophrenia from bench to bedside: Discovery of a striatal co-expression risk gene set that predicts in vivo measures of striatal function": Compressed zipped directory containing scripts used for this study.: 2.sda_postprocessing.R

```
###############################################################################################################
############################################### SDA post_processing ############################################
###############################################################################################################

rm(list = ls())

### set parameters and folder used
samples = 120
tissues = c("CN","DLPFC","HP")
genes = 20475

source("2a.sda_postProcess.R")         

#### if th=0.6 than correlation > 0.4 
threshold = 0.6 

final_SDA = postProcess(data_name = "caud_dlpfc_hippo_GTEx",
                        samples = samples,tissues = tissues,genes = genes,
                        threshold = threshold,
                        data_dir = "")

final_SDA = rename(final_SDA,samples = samples,tissues = NULL,
                   threshold = threshold,
                   preProcess_dir = "",
                   output_dir = "")

### save 
save(final_SDA, file = paste0("final_SDA.RData"))
```
