## Supplementary material for "Dopamine and schizophrenia from bench to bedside: Discovery of a striatal co-expression risk gene set that predicts in vivo measures of striatal function": Compressed zipped directory containing scripts used for this study.: 2b.sda_postprocessing_functions.R

```
###############################################################################################################
############################################### SDA post_processing ############################################
###############################################################################################################

### function to read SDA 10-runs output 
read_output_SDA = function (file_name) {
  
  folders = c()
  output_SDA = list()
  
  library(data.table) 
  
  read_output <- function(folder) {
    # reads all estimates 
    # e.g folder <- "my_output_folder/it2000"
    # if you are in the folder containing the estimates, folder <- "../it2000"
    out <- list()
    #folder_exists <- system(paste0("[ -d ", folder, " ] && echo true || echo false"), intern=TRUE)
    
    #stopifnot(folder_exists=='true')
    
    # get list of all files in folder
    files <- list.files(folder)
    # read all files
    for (file in files) {
      out[[file]] <- as.matrix(fread(paste0(folder, "/", file)))
    }
    
    out1 <- reformat_data(out) # from the estimates, get N, L, P, ncomps and reformat data
    for (i in names(out1)) {
      out[[i]] <- out1[[i]]
    }
    out$X1 <- NULL; out$S1 <- NULL; out$B1 <- NULL 
    
    return(out)
  }
  
  reformat_data <- function(out) {
    nam <- names(out)
    
    est <- list()
    est$A <- out$A
    
    est$N <- nrow(est$A) # number individuals
    est$C <- ncol(est$A) # number components
    
    num_X_mats <- length(nam[grep("X[0-9]?", nam)])
    stopifnot(length(nam[grep("S[0-9]?", nam)])==num_X_mats)
    stopifnot(num_X_mats != 0)
    num_B_mats <- length(nam[grep("B[0-9]?", nam)])
    
    if (num_B_mats == 0 && num_X_mats == 1) { # 2D matrix factorisation
      est$X <- out$X1
      est$S <- out$S1
      
    } else if (num_B_mats == 1 && num_X_mats == 1) { # 3D matrix factorisation
      est$X <- out$X1; est$S <- out$S1
      est$B <- out$B1
      est$P <- nrow(est$B)
      
    } else {
      print("problem with estimates")
    }
    
    return(est)
  }
  
  for (i in 1:10) {
    
    dir = paste0(file_name,"_",i,".run/it3000")
    folders = c(folders,dir)
    
  }
  
  for (out in folders){
    
    
    out1 = read_output(out)
    
    output_SDA[[out]] = out1
    
    
  }
  
  ### give same components names to gene_loadings and PIP matrix as the ones in individual and tissue matrix
  
  for (i in 1:10){
    
    row.names(output_SDA[[paste0(file_name,"_",i,".run/it3000")]]$X) = colnames(output_SDA[[paste0(file_name,"_",i,".run/it3000")]]$A)
    row.names(output_SDA[[paste0(file_name,"_",i,".run/it3000")]]$S) = colnames(output_SDA[[paste0(file_name,"_",i,".run/it3000")]]$A)
    
  }
  
  rm(list = setdiff(ls(),"output_SDA"))
  return(output_SDA)
  
} 

### function to obtain the components absolute correlation matrix across 10 runs
abscorrmatrix = function(output_SDA_subset){
  
  components.matrix = Reduce(cbind,
                             lapply(output_SDA_subset,function(x) x$A))
  components.matrix = abs(cor(components.matrix,method = "pearson"))
  weird = colnames(components.matrix)[ apply(components.matrix,2,function(x) sum(is.na(x))) == nrow(components.matrix)-1 ]
  components.matrix = components.matrix[!row.names(components.matrix) %in% weird,
                                        !colnames(components.matrix) %in% weird]
  return(components.matrix)
  
} 

### function to obtain individual,tissue,gene and PIP matrices 
### with robust components (clusters with a minimum membership size of 5)
clusterCombine = function(clusters,output_SDA_subset,nsubj = samples,ntissue = length(tissues),ngene = genes,th = 0.5,file_name){
  
  clusterCombine_indiv = function(cluster,f.name = file_name){
    
    cluster_combined = sapply(cluster,function(comp){
      
      path = paste0(f.name,"_",strsplit(comp,"\\.")[[1]][[2]],".run/it3000")               
      # comp_name = strsplit(comp,"\\.")[[1]][[1]]
      output_SDA_subset[[path]]$A[1:nsubj,comp] 
      
    },USE.NAMES = T)
    cluster_combined = rowMeans(cluster_combined)
    return(cluster_combined)
    
    
  }    #####################################################
  clusterCombine_tissue = function(cluster,f.name = file_name){
    
    cluster_combined = sapply(cluster,function(comp){
      
      path = paste0(f.name,"_",strsplit(comp,"\\.")[[1]][[2]],".run/it3000")               
      # comp_name = strsplit(comp,"\\.")[[1]][[1]]
      output_SDA_subset[[path]]$B[1:ntissue,comp] 
      
    },USE.NAMES = T)
    cluster_combined = rowMeans(cluster_combined)
    return(cluster_combined)
    
    
  }   ######### functions to combine components ###########
  clusterCombine_genes = function(cluster,f.name = file_name){
    
    cluster_combined = sapply(cluster,function(comp){
      
      path = paste0(f.name,"_",strsplit(comp,"\\.")[[1]][[2]],".run/it3000")               
      # comp_name = strsplit(comp,"\\.")[[1]][[1]]
      t(output_SDA_subset[[path]]$X)[1:ngene,comp] 
      
    },USE.NAMES = T)
    cluster_combined = rowMeans(cluster_combined)
    return(cluster_combined)
    
    
  }    ######### in each cluster for each specific matrix ##
  clusterCombine_PIP =   function(cluster,f.name = file_name){
    
    cluster_combined = sapply(cluster,function(comp){
      
      path = paste0(f.name,"_",strsplit(comp,"\\.")[[1]][[2]],".run/it3000")               
      # comp_name = strsplit(comp,"\\.")[[1]][[1]]
      t(output_SDA_subset[[path]]$S)[1:ngene,comp] 
      
    },USE.NAMES = T)
    cluster_combined = apply(cluster_combined,1,median)
    return(cluster_combined)
    
    
  }    #####################################################
  
  ### obtaining clusters with minimum membership size of 5
  cluster.cut = cutree(clusters,h = th)
  cluster.df = data.frame(cluster = 1:length(table(cluster.cut)),freq = as.vector(table(cluster.cut)))
  clusters = list()
  for (i in 1:nrow(cluster.df)){
    
    cluster.i = names(cluster.cut[cluster.cut %in% cluster.df$cluster[[i]]])
    clusters[[paste0("V",i)]] = cluster.i
    
  }
  cluster.df$freq.unique = sapply(clusters,function(clust){
    
    new = sapply(clust,function(x)strsplit(x,"\\.")[[1]][[2]])
    length(unique(new))
    
    
  },USE.NAMES = T)
  cluster.df = cluster.df[which(cluster.df$freq.unique >= 5),]
  
  ### obtaining names of components to be combined across 10 runs 
  clusters = list()
  for (i in 1:nrow(cluster.df)){
    
    cluster.i = names(cluster.cut[cluster.cut %in% cluster.df$cluster[[i]]])
    clusters[[paste0("V",i)]] = cluster.i
    
  }
  
  ### obtaining robust components for each specific matrix
  cluster_indiv = sapply(clusters,clusterCombine_indiv)
  cluster_tissue = sapply(clusters,clusterCombine_tissue)
  cluster_gene = sapply(clusters,clusterCombine_genes)
  cluster_PIP = sapply(clusters,clusterCombine_PIP)
  
  ### obtaining final SDA matrices
  A = cluster_indiv
  B = cluster_tissue
  X = t(cluster_gene)
  S = t(cluster_PIP)
  
  return(list(A = A,
              B = B,
              X = X,
              S = S,
              N = samples,
              C = length(clusters)))
  
  
  
} 
```
