## Supplementary material for "Dopamine and schizophrenia from bench to bedside: Discovery of a striatal co-expression risk gene set that predicts in vivo measures of striatal function": Compressed zipped directory containing scripts used for this study.: 3a.sda_metadata_functions.R

```
###############################################################################################################
######################## SDA metadata  ####################################
###############################################################################################################

metadata_GTEx = function(data_dir,data_name,
                         tissues,
                         common_metadata,
                         tissue_specific_metadata,
                         sample_list,
                         threshold,
                         output_dir)
{
          #browser()
          load(paste0(data_dir,data_name,".RData"))
          sample_list = lapply(sample_list,function(x)
          {
                    x$RIN = as.numeric(x$RIN)
                    row.names(x) = x$SAMPID
                    x
                    
          })
          metadata <- sample_list[[1]][,common_metadata]
          for (i in c("Sex")){metadata[[i]] = as.factor(metadata[[i]])}
          batch_data = lapply(tissues,t=tissue_specific_metadata,function(x,t)
          {
                    new = sample_list[[x]][,t]
                    colnames(new) = paste0(t,"_",x)
                    new
                    
          })
          batch_data = as.data.frame(batch_data)
          data = cbind(metadata,batch_data[match(row.names(metadata),row.names(batch_data)),])
          stopifnot(identical(row.names(final_SDA$A),data$SAMPID))
          data = cbind(data,final_SDA$A)
          
}
```
