## Supplementary material for "Dopamine and schizophrenia from bench to bedside: Discovery of a striatal co-expression risk gene set that predicts in vivo measures of striatal function": Compressed zipped directory containing scripts used for this study.: 6.sda_genes_components.R

```
###############################################################################################################
############################################### SDA genes/components ##################################
###############################################################################################################

load(paste0("final_SDA.RData"))
final_SDA = final_SDA_filtered

### list of genes with loading != 0 in each component
geneMatrix = t(final_SDA[[paste0("X_sig_0.5")]])

geneList = sapply(colnames(geneMatrix),function(x){
  
  row.names(geneMatrix[which(geneMatrix[,x] != 0),])
  
  
})

### save 
save(geneList, file = "genes_components_overlap.RData")
```
