## Supplementary material for "Dopamine and schizophrenia from bench to bedside: Discovery of a striatal co-expression risk gene set that predicts in vivo measures of striatal function": Compressed zipped directory containing scripts used for this study.: 7.sda_tissue_score.R

```
###############################################################################################################
######################## SDA tissue specificity ##################################
###############################################################################################################

my_scale = function(x){return(x/max(abs(x)))}

load(paste0("final_SDA.RData"))
tissue.score = as.data.frame(apply(final_SDA$B,2,my_scale))

save(tissue.score, file = paste0("tissue.score.RData"))
```
