## Supplementary material for "Dopamine and schizophrenia from bench to bedside: Discovery of a striatal co-expression risk gene set that predicts in vivo measures of striatal function": Compressed zipped directory containing scripts used for this study.: 1.Run_enrichment.R

```
####################################################################################################
#                                                                                                  #
#                    Script to run various enrichments on gene_component_list:                     #
#                   GWAS risk genes, DEGs, DMGs, TWAS, GO, MAGMA, Cell Specificity etc.            #
#                                                                                                  #
####################################################################################################

rm(list=ls())
require(limma)
source('Enrich.R')
load('sda_preprocessing.RData')
load(paste0('genes_components_overlap.RData'))
geneMap = gene_common

#### Run ####
Enrichment(Net = geneList, Net_type = 'SDA', Net_name = paste0("LIBD_discovery"),
           Map = geneMap,
           enrichment_type = c("Top_GWAS","MAGMA","H-MAGMA",
                               "DEGs","DMGs","pleiotropic_Lof_Denovo_CNVs",
                                "Cell Specificity", "GO"),
           Pathology = c('SCZ','MDD','BIP','SA','ASD','ADHD','OCD','PTSD','CD','UC'),
           MAGMA_pathology = c('SCZ','MDD','BIP','SA','ASD','ADHD','OCD','PTSD','CD','UC'),
           magma_directory = '~/Desktop/magma_v1/',
           directory_target = 'script.data.enrichment/RData/',
           directory_script = 'script.data.enrichment/script/',
           directory_output = paste0('output/'))
```
