## Supplementary material for "Dopamine and schizophrenia from bench to bedside: Discovery of a striatal co-expression risk gene set that predicts in vivo measures of striatal function": Compressed zipped directory containing scripts used for this study.: Cell_specificity_enrich.R

```
##################################################################################################
### CELL-SPECIFICITY geneset enrichment analysis with limma ######################################
##################################################################################################

Cell_Specificity = function(Net, network.name, genemap, target.directory, output.directory){
          
          library(limma)
          heatmapL = function(cor_mat, col, x_name, y_name, title) {
                    require(ggplot2)
                    require(reshape2)
                    melt_mat <- melt(cor_mat)
                    ifelse(max(results.log10)>=50,{increase = 50},{increase = 10})
                    p =  ggplot(melt_mat, aes(Var1, Var2)) + geom_tile(aes(fill = value) , colour = "white") +
                              geom_text(data = melt_label, label = melt_label$label, size = 2.5, col='red') +
                              scale_fill_gradient(limits=c(0, max(results.log10)+1), breaks=seq(0, max(results.log10)+1,by=increase), 
                                                  low = "white", high = col, guide = guide_colorbar(title="-log10(fdr-adj p)")) +
                              theme(axis.text.y = element_text(angle=0, hjust=0.5, size=15)) +
                              theme(axis.text.x = element_text(angle=90, vjust=0.5, size=10)) +
                              theme(axis.title = element_text(size = 30)) +
                              xlab(x_name) + ylab(y_name) + ggtitle(title)
                    return(p)
          }
          ### load target
          load(paste0(target.directory,"Cell_specificity.RData"))
          ### load network data
          candidate.list = sapply(names(Net),function(x){
                    
                    geneMap_new = genemap[which(genemap$ensembl %in% Net[[x]]),]
                    Net[[x]] = geneMap_new$hgnc
                    
          })
          load(paste0(target.directory,"m_to_h_homologs.RData"))
          mouse_to_human_homologs = m_to_h_homologs
          candidate.list.mouse = lapply(candidate.list, function(x) 
                    mouse_to_human_homologs$MGI.symbol[mouse_to_human_homologs$HGNC.symbol %in% x] )
          ######  human ######
          condition_h = names(Cell_specificity$human) %in% c("DRONC_human","DRONC_sub_human","AIBS")
          for(h in 1:length(Cell_specificity$human)){ 
                    
                    ### GSEA
                    ifelse(condition_h[h],{
                              ### load target
                              namePSI = names(Cell_specificity$human[h])
                              specificity = Cell_specificity$human[[h]] 
                              results = sapply(colnames(specificity), function(v) 
                                        sapply(candidate.list, function(m) 
                                                  geneSetTest(index = row.names(specificity) %in% m, statistics = specificity[,v], 
                                                              alternative = "up", type= "t", ranks.only = T, nsim=9999) ))
                              results[results == 0] = 1e-200
                              results.fdr = sapply(colnames(results), function(x) p.adjust(results[,x],method = "fdr", n = (nrow(results)*ncol(results))))
                    },{
                              ### load target
                              namePSI = paste0(names(Cell_specificity$human[h]),"_human")
                              specificity = Cell_specificity$human[[h]]   
                              ### GSEA
                              results = sapply(names(specificity), function(v) 
                                        sapply(candidate.list, function(m) 
                                                  geneSetTest(index = specificity[[v]][,"gene"] %in% m, statistics = specificity[[v]][,1], 
                                                              alternative = "up", type= "t", ranks.only = T, nsim=9999) ))
                              results[results == 0] = 1e-200
                              results.fdr = sapply(colnames(results), function(x) p.adjust(results[,x],method = "fdr"))
                    })
                    
                    results.log10 = -log10(results.fdr)
                    ### threshold
                    th.fdr = -log10(0.05)
                    ### melt p-value
                    require(reshape2)
                    melt_label = melt(results.log10)
                    melt_label$label = ifelse(melt_label$value >= th.fdr, "#", "")
                    ### save
                    save(results,results.fdr,results.log10, th.fdr, file = paste0(output.directory,"/GSEA_", network.name, "_", namePSI, ".RData"))
                    ####################################################################################
                    ### HEATMAP with labels labels EMPIRICAL P-VALUE
                    pdf(paste0(output.directory,"/heatmap_", network.name, "_", namePSI, ".pdf"), width = 12,  height = 4, useDingbats = FALSE)
                    print(heatmapL(results.log10, col = 'steelblue', x_name = "Robust components", y_name = namePSI, title = ''))
                    dev.off()
          }
          
          ######  mouse ######
          condition_m = names(Cell_specificity$mouse) %in% c("DRONC_mouse","DRONC_sub_mouse","allKI_mouse","allKI_sub_mouse") 
          for(m in 1:length(Cell_specificity$mouse)){ 
                    
                    ### GSEA
                    ifelse(condition_m[m],{
                              ### load target
                              namePSI = names(Cell_specificity$mouse[m])
                              specificity = Cell_specificity$mouse[[m]] 
                              results = sapply(colnames(specificity), function(v) 
                                        sapply(candidate.list.mouse, function(y) 
                                                  geneSetTest(index = row.names(specificity) %in% y, statistics = specificity[,v], 
                                                              alternative = "up", type= "t", ranks.only = T, nsim=9999) ))
                              results[results == 0] = 1e-200
                              results.fdr = sapply(colnames(results), function(x) p.adjust(results[,x],method = "fdr", n = (nrow(results)*ncol(results))))
                    },{
                              ### load target
                              namePSI = paste0(names(Cell_specificity$mouse[m]),"_mouse")
                              specificity = Cell_specificity$mouse[[m]]   
                              ### GSEA
                              results = sapply(names(specificity), function(v) 
                                        sapply(candidate.list, function(y) 
                                                  geneSetTest(index = specificity[[v]][,"gene"] %in% y, statistics = specificity[[v]][,1], 
                                                              alternative = "up", type= "t", ranks.only = T, nsim=9999) ))
                              results[results == 0] = 1e-200
                              results.fdr = sapply(colnames(results), function(x) p.adjust(results[,x],method = "fdr"))
                    })
                    results.log10 = -log10(results.fdr)
                    ### threshold
                    th.fdr = -log10(0.05)
                    ### melt p-value
                    require(reshape2)
                    melt_label = melt(results.log10)
                    melt_label$label = ifelse(melt_label$value >= th.fdr, "#", "")
                    ### save
                    save(results,results.fdr,results.log10, th.fdr, file = paste0(output.directory,"/GSEA_", network.name, "_", namePSI, ".RData"))
                    ####################################################################################
                    ### HEATMAP with labels labels EMPIRICAL P-VALUE
                    if(m == which(names(Cell_specificity$mouse) == "allKI_sub_mouse")){
                              pdf(paste0(output.directory,"/heatmap_", network.name, "_", namePSI, ".pdf"), width = 20,  height = 30, useDingbats = FALSE)
                              print(heatmapL(results.log10, col = 'navajowhite4', x_name = "Robust components", y_name = namePSI, title = ''))
                              dev.off()}else{
                                        pdf(paste0(output.directory,"/heatmap_", network.name, "_", namePSI, ".pdf"), width = 16,  height = 12, useDingbats = FALSE)
                                        print(heatmapL(results.log10, col = 'navajowhite4', x_name = "Robust components", y_name = namePSI, title = ''))
                                        dev.off()  
                                        
                              }
          }

######  human_mouse ######
namePSI = "Zhang_Darmanis_Zeisel_Tasic_mouse_human"
specificity = Cell_specificity$human_mouse   
### GSEA
results = sapply(names(specificity), function(v) 
          sapply(candidate.list, function(y) 
                    geneSetTest(index = specificity[[v]][,"gene"] %in% y, statistics = specificity[[v]][,1], 
                                alternative = "up", type= "t", ranks.only = T, nsim=9999) ))
results[results == 0] = 1e-200
results.fdr = sapply(colnames(results), function(x) p.adjust(results[,x],method = "fdr"))
results.log10 = -log10(results.fdr)
### threshold
th.fdr = -log10(0.05)
### melt p-value
require(reshape2)
melt_label = melt(results.log10)
melt_label$label = ifelse(melt_label$value >= th.fdr, "#", "")
### save
save(results,results.fdr,results.log10, th.fdr, file = paste0(output.directory,"/GSEA_", network.name, "_", namePSI, ".RData"))
####################################################################################
### HEATMAP with labels labels EMPIRICAL P-VALUE
pdf(paste0(output.directory,"/heatmap_", network.name, "_", namePSI, ".pdf"), width = 14,  height = 7, useDingbats = FALSE)
print(heatmapL(results.log10, col = 'darkorange', x_name = "Robust components", y_name = namePSI, title = ''))
dev.off()
}          
```
