## Supplementary material for "Dopamine and schizophrenia from bench to bedside: Discovery of a striatal co-expression risk gene set that predicts in vivo measures of striatal function": Compressed zipped directory containing scripts used for this study.: empirical_enrichment.R

```
Enrich = function(ll.target, ll.components, geneMap,perm ,network_name, biotype, output.directory){
          
          #source("C:/Users/lsporte1/Desktop/script.data.enrichment/script/SDA/myPermutation.R")
          #myPermutation = cmpfun(myPermutation)
          condition = grep(biotype,c("all_biotypes","protein_coding"))
          switch(condition,{
                    
                    compList = lapply(geneMap$ensembl,components = ll.components,function(gene,components){
                              comp =  sapply(components,intersect,gene)
                              comp = names(Filter(length,comp))
                    })
                    names(compList) = geneMap$ensembl
                    compList[sapply(compList,length)==0] = "grey"
                    grey = setdiff(geneMap$ensembl,unique(unlist(ll.components)))
          },{
                    ll.components = sapply(ll.components,function(x){
                              with(geneMap,ensembl[ensembl %in% x & gene_type == "protein_coding"])
                    })
                    prot.ensembl = with(geneMap,ensembl[gene_type == "protein_coding"])
                    compList = lapply(prot.ensembl,components = ll.components,function(gene,components){
                              comp =  sapply(components,intersect,gene)
                              comp = names(Filter(length,comp))
                    })
                    names(compList) = prot.ensembl
                    compList[sapply(compList,length)==0] = "grey"
                    grey = setdiff(prot.ensembl,unique(unlist(ll.components)))
          })
          
          p = sapply(compList,length)/(length(unlist(ll.components)) + length(grey)) 
          all_genes = union(unique(unlist(ll.components)),grey)
          p = p[match(all_genes,names(p))]
          set.seed(123)
          result = myPermutation(target = ll.target,components = ll.components,
                                 all_genes = all_genes,p = p, perm = perm)
          for(target_name in names(ll.target)){
                    
                    enrich.matrix.pval = result$enrich.matrix[[target_name]]
                    real.enrich = result$real.enrich[[target_name]]
                    
                    ## assign and save
                    assign("enrich.matrix.pval"                              , enrich.matrix.pval)
                    assign("Modules"                                   , ll.components)
                    assign(paste("enrich.list", network_name,  sep=".")                   , real.enrich)
                    
                    save(list = c(paste("enrich.matrix.pval"),
                                  paste("Modules"),
                                  paste("enrich.list", network_name, sep=".")),
                         file = paste0(output.directory,"/",target_name, ".", network_name, ".RData"))
                    
                    ### plot GENES
                    require(grDevices)
                    #browser()
                    pdf(paste0(output.directory,"/",paste("enrich", target_name, network_name, "pdf", sep=".")),width = 18, useDingbats = FALSE)
                    par(mar = c(8,5,5,2), lwd = 96/72, ps = 8, las = 2)
                    #browser()
                    for(i in colnames(enrich.matrix.pval)) {
                              
                              bp = barplot(height = -log10(enrich.matrix.pval[,i]),
                                           col = "grey", names.arg = row.names(enrich.matrix.pval), space = 0,
                                           main = paste0(target_name, ' enrichment of ', network_name,
                                                         '\n',' list: ', i),
                                           ylab = '-log10(p-value)', cex.lab = 3, cex.axis = 2, ylim = c(0,max(-log10(enrich.matrix.pval))+2),las = 2)
                              
                              p.line = -log10(0.05)
                              p.text = -log10(0.05)+0.1
                              
                              abline(h = p.line, lwd = 2, lty = 2, col = c('darkred'))
                              text(x = 5, y = p.text, labels = c('pvalue=0.05'), col = c('darkred'))
                              
                              signif = which(enrich.matrix.pval[,i] < 0.05)
                              signif.hit = real.enrich[signif,i]
                              
                              if (length(signif) > 0){
                                        text(x = bp[signif], y = -log10(enrich.matrix.pval[signif, i])+1.5,
                                             labels = as.numeric(signif.hit), cex = 1.5)}
                              
                    }
                    dev.off()
          }
          
          
}
```
