## Supplementary material for "Dopamine and schizophrenia from bench to bedside: Discovery of a striatal co-expression risk gene set that predicts in vivo measures of striatal function": Compressed zipped directory containing scripts used for this study.: Enrich.R

```
###### Multi-Enrichment function ######
Enrichment = function(Net = NULL, Net_type = NULL, Net_name = NULL, list_enrich = FALSE,
                      Map = NULL,
                      Loci = NULL,
                      gene_PIP = NULL,
                      modules_WGCNA = NULL,
                      gene_ID = NULL, 
                      enrichment_type = c("Top_GWAS","Loci_enrichment","MAGMA","H-MAGMA", "eMAGMA", "TWAS","DMGs","pleiotropic_Lof_Denovo_CNVs",
                                          "DEGs", "Druggable_genes","DEGs_MDD_Single.cell",
                                          "Cell Specificity", "GO", "WGCNA"),
                      Pathology = c("SCZ","MDD","BIP","ASD","ADHD","OCD","PTSD","SA",
                                    "AD","PD","ALS","MS","RA","CD","UC"),
                      MAGMA_pathology = c("SCZ","MDD","BIP","ASD","ADHD","OCD","PTSD","SA",
                                           "AD","PD","ALS","CUD","RA","CD","UC"),
                      magma_directory = NULL,
                      directory_target = "/script.data.enrichment/RData/",
                      directory_script = "/script.data.enrichment/script/",
                      directory_output = NULL){
          
          if(isTRUE(list_enrich)){
                    print(paste("Type of enrichments ------> ",paste(enrichment_type,collapse = ", ")))
                    print(paste("Pathologies available for Top_GWAS ------> ",paste(Pathology,collapse = ",")))
                    print(paste("Pathologies available for MAGMA/H-MAGMA ------> ",paste(MAGMA_pathology,collapse = ",")))}
          else if(Net_type == "WGCNA"){
                    
                    source(paste0(directory_script,Net_type,"/Enrich_",Net_type,".R"))
                    #if(!any(colnames(Net) %in% c("Gene","Symbol"))){stop("Set Network Ensembl ID and HGNC Symbol as 'Gene' and 'Symbol'")}
                    Enrich_WGCNA(Network = Net,
                                 Network_type = Net_type,
                                 Network_name = Net_name,
                                 GeneMap = Map,
                                 LociMap = Loci,
                                 ID = gene_ID,
                                 enrich_type = enrichment_type,
                                 pathology = Pathology,
                                 magma_pathology = MAGMA_pathology,
                                 MAGMA_directory = magma_directory,
                                 Target_directory = directory_target,
                                 Script_directory = directory_script,
                                 Output_directory = directory_output)
          } 
          else if(Net_type == "SDA"){
                    
                    source(paste0(directory_script,Net_type,"/Enrich_",Net_type,".R"))
                    Enrich_SDA(Network = Net,
                               Network_type = Net_type,
                               Network_name = Net_name,
                               GeneMap = Map,
                               LociMap = Loci,
                               PIP = gene_PIP,
                               WGCNA_modules = modules_WGCNA,
                               ID = gene_ID,
                               enrich_type = enrichment_type,
                               pathology = Pathology,
                               magma_pathology = MAGMA_pathology,
                               MAGMA_directory = magma_directory,
                               Target_directory = directory_target,
                               Script_directory = directory_script,
                               Output_directory = directory_output)
          }
} 
```
