## Supplementary material for "Dopamine and schizophrenia from bench to bedside: Discovery of a striatal co-expression risk gene set that predicts in vivo measures of striatal function": Compressed zipped directory containing scripts used for this study.: Enrich_SDA.R

```
############################   SDA enrichments ###########################
##########################################################################
Enrich_SDA = function(Network,
                      Network_type,
                      Network_name,
                      GeneMap,
                      LociMap,
                      PIP,
                      WGCNA_modules,
                      ID,
                      enrich_type,
                      pathology = c("SCZ","MDD","BIP","ASD","ADHD","OCD","PTSD","SA",
                                    "AD","PD","ALS","MS","RA","CD","UC"),
                      magma_pathology = c("SCZ","MDD","BIP","ASD","ADHD","OCD","PTSD","SA",
                                          "AD","PD","ALS","CUD","RA","CD","UC"),
                      Target_directory,
                      Script_directory,
                      Output_directory,
                      MAGMA_directory){
  
  require(compiler)
  require(future);require(furrr);require(purrr)
  #### computing enrichment with permutations
  if("Top_GWAS" %in% enrich_type){full_condition = c(enrich_type,pathology)} else full_condition = enrich_type
  condition = full_condition %in% c("SCZ","MDD","BIP","ASD","ADHD","OCD","PTSD","SA",
                                    "AD","PD","ALS","MS","RA","CD","UC",
                                    "TWAS","DEGs","DEGs_MDD_Single.cell","Druggable_genes","DMGs","pleiotropic_Lof_Denovo_CNVs")
  if(any(condition)){
    
    dir = "TopGWAS.TWAS.DEGs.DMGs.Druggable.LoF.Denovo.pleiotropic.CNVs"
    if(!dir %in% list.files(Output_directory)){dir.create(paste0(Output_directory,dir))}
    source(paste0(Script_directory,Network_type,"/myPermutation.R"))
    source(paste0(Script_directory,Network_type,"/empirical_enrichment.R"))
    # Enrich = cmpfun(Enrich)
    # myPermutation = cmpfun(myPermutation)
    ll.target = list()
    for (en in full_condition[condition]){
      
      new_env = environment()
      
      ###### creating list of target genes ######
      if(en %in% c("SCZ","MDD","BIP","ASD","ADHD","OCD","PTSD","SA",
                   "AD","PD","ALS","MS","RA","CD","UC"))
      {load(paste0(Target_directory,tolower(en),"/PGC.gene.lists.RData"),envir = new_env)
      }else load(paste0(Target_directory,en,".RData"),envir = new_env)
      target = grep("protein\\.coding|all\\.biotypes|TWAS|DEGs|Druggable|DMGs|pleiotropic",ls(new_env),value = T)
      for (list in target){ll.target[[list]] = get(list,envir = new_env)}
      
    }
    rm(list = grep("protein\\.coding|all\\.biotypes|TWAS|DEGs|DMGs|Druggable|pleiotropic|^new_env",ls(),value = T))
    ## load and set components list
    ll.components = Network
    ## load and set target list
    ll.target.all = ll.target[!names(ll.target) %in% grep("protein\\.coding",names(ll.target),value = T)]
    ll.target.pc = ll.target[names(ll.target) %in% grep("protein\\.coding",names(ll.target),value = T)]
    ## enrichment computation
    Enrich(ll.target.all,ll.components,perm = 10000,biotype = "all_biotypes",geneMap = GeneMap,
           network_name = Network_name,output.directory = paste0(Output_directory,dir))
    Enrich(ll.target.pc,ll.components,perm = 10000,biotype = "protein_coding",geneMap = GeneMap,
           network_name = Network_name,output.directory = paste0(Output_directory,dir))
    print(paste0(full_condition[condition]," enrichment computed with success"))
    
  }
  rm(list = grep("PGC|TWAS|DEGs|Druggable|^new_env
                         |ll.target|ll.components",ls(),value = T))
  
  #### computing Loci enrichment with permutations
  if("Loci_enrichment" %in% enrich_type){full_condition = c(enrich_type,pathology)} else full_condition = enrich_type
  condition = full_condition %in% c("SCZ")
  if(any(condition)){
    
    dir = "Loci_enrichment"
    if(!dir %in% list.files(Output_directory)){dir.create(paste0(Output_directory,dir))}
    source(paste0(Script_directory,Network_type,"/myLociPermutation.R"))
    source(paste0(Script_directory,Network_type,"/empirical_Loci_enrichment.R"))
    Enrich = cmpfun(Enrich)
    myLociPermutation = cmpfun(myLociPermutation)
    ll.target = list()
    for (en in full_condition[condition]){
      
      new_env = environment()
      ###### creating list of target genes ######
      load(paste0(Target_directory,tolower(en),"/PGC.loci.lists.RData"),envir = new_env)
      target = grep("protein\\.coding|all\\.biotypes",ls(new_env),value = T)
      for (list in target){ll.target[[list]] = get(list,envir = new_env)}
      
      
    }
    rm(list = grep("protein\\.coding|all\\.biotypes|TWAS|DEGs|DMGs|Druggable|pleiotropic|^new_env",ls(),value = T))
    ## load and set components list
    ll.components = lapply(Network,function(x){
      intersect(x,LociMap$ensembl)
    })
    ## load and set target list
    ll.target.all = ll.target[!names(ll.target) %in% grep("protein\\.coding",names(ll.target),value = T)]
    ll.target.pc = ll.target[names(ll.target) %in% grep("protein\\.coding",names(ll.target),value = T)]
    ## enrichment computation
    # Enrich(ll.target.all,ll.components,perm = 10000,biotype = "all_biotypes",lociMap = LociMap,
    #        network_name = Network_name,output.directory = paste0(Output_directory,dir))
    Enrich(ll.target.pc,ll.components,perm = 10000,biotype = "protein_coding",lociMap = LociMap,
           network_name = Network_name,output.directory = paste0(Output_directory,dir))
    print(paste0(full_condition[condition]," Loci enrichment computed with success"))
    
  }
  rm(list = grep("PGC|TWAS|DEGs|Druggable|^new_env
                         |ll.target|ll.components",ls(),value = T))
  
  if("Cell Specificity" %in% enrich_type){
    
    source(paste0(Script_directory,Network_type,"/Cell_specificity_enrich.R"))
    Cell_Specificity = cmpfun(Cell_Specificity)
    #if(!paste0("PIP_",PIP) %in% list.files(Output_directory)){dir.create(paste0(Output_directory,"PIP_",PIP))}
    if(!"Cell_specificity" %in% list.files(Output_directory)){dir.create(paste0(Output_directory,"Cell_specificity"))}
    Cell_Specificity(Net = Network,
                     network.name = Network_name,
                     genemap = GeneMap,
                     target.directory = Target_directory,
                     output.directory = paste0(Output_directory,"Cell_specificity"))
    print("Cell Specificity enrichment computed with success")
  }
  
  if("GO" %in% enrich_type){
    
    #browser()
    source(paste0(Script_directory,Network_type,"/GO_enrich.R"))
    GO_enrich = cmpfun(GO_enrich)
    #if(!paste0("PIP_",PIP) %in% list.files(Output_directory)){dir.create(paste0(Output_directory,"PIP_",PIP))}
    if(!"GO" %in% list.files(Output_directory)){dir.create(paste0(Output_directory,"GO"))}
    GO_enrich(Net = Network,
              network_name = Network_name,Net_type = Network_type,
              geneMap = GeneMap,
              target_name = c("GO","KEGG","Reactome"),
              script.directory = Script_directory,
              target.directory = Target_directory,
              pvalue = c(1),
              output.directory = paste0(Output_directory,"GO"))
    print("GO enrichment computed with success")
  }
  
  if("WGCNA" %in% enrich_type){
    
    source(paste0(Script_directory,Network_type,"/WGCNA_modules_enrich.R"))
    source(paste0(Script_directory,Network_type,"/hypergeo_enrichment.R"))
    #if(!paste0("PIP_",PIP) %in% list.files(Output_directory)){dir.create(paste0(Output_directory,"PIP_",PIP))}
    if(!"WGCNA_modules" %in% list.files(Output_directory)){dir.create(paste0(Output_directory,"WGCNA_modules"))}
    WGCNA_modules_enrich(Net = Network,
                         network_name = Network_name,
                         genemap = GeneMap,
                         WGCNA = WGCNA_modules,
                         ID = ID,
                         target_name = "WGCNA_modules",
                         target.directory = Target_directory,
                         output.directory = paste0(Output_directory,"PIP_",PIP,"/WGCNA_modules"))
    print("WGCNA modules enrichment computed with success")
  }
  
  if("MAGMA" %in% enrich_type){
    
    source(paste0(Script_directory,Network_type,"/MAGMA_enrich.R"))
    # source(paste0(Script_directory,Network_type,"/MAGMA_enrich_preprocess.R"))
    MAGMA_enrich = cmpfun(MAGMA_enrich)
    # MAGMA_enrich_preprocess = cmpfun(MAGMA_enrich_preprocess)
    pathologies = magma_pathology[magma_pathology %in% c("SCZ","MDD","BIP","ASD","ADHD","OCD","PTSD","SA",
                                                                 "AD","PD","ALS","CUD","RA","CD","UC")]
    if(!"MAGMA" %in% list.files(Output_directory)){dir.create(paste0(Output_directory,"MAGMA"))}
    purrr::walk(pathologies,~{
      patho = .x
      MAGMA_enrich(Net = Network,
                   Net_name = Network_name,
                   genemap = GeneMap,
                   pathology = patho,
                   target.directory = paste0(Target_directory,"MAGMA/"),
                   output.directory = paste0(Output_directory),
                   magma.directory = MAGMA_directory)
      print(paste0("MAGMA enrichment for ",patho," computed with success"))
    })
  }
  if("H-MAGMA" %in% enrich_type){
    
    source(paste0(Script_directory,Network_type,"/H-MAGMA_enrich.R"))
    # source(paste0(Script_directory,Network_type,"/MAGMA_enrich_preprocess.R"))
    H.MAGMA_enrich = cmpfun(H.MAGMA_enrich)
    # MAGMA_enrich_preprocess = cmpfun(MAGMA_enrich_preprocess)
    pathologies = magma_pathology[magma_pathology %in% c("SCZ","MDD","BIP","ASD","ADHD","OCD","PTSD","SA",
                                                                 "AD","PD","ALS","CUD","RA","CD","UC")]
    if(!"H-MAGMA" %in% list.files(Output_directory)){dir.create(paste0(Output_directory,"H-MAGMA"))}
    purrr::walk(pathologies,~{
      patho = .x
      H.MAGMA_enrich(Net = Network,
                     Net_name = Network_name,
                     genemap = GeneMap,
                     pathology = patho,
                     target.directory = paste0(Target_directory,"H-MAGMA/"),
                     output.directory = paste0(Output_directory),
                     magma.directory = MAGMA_directory)
      print(paste0("H-MAGMA enrichment for ",patho," computed with success"))
    })
  }
  if("eMAGMA" %in% enrich_type){
    
    source(paste0(Script_directory,Network_type,"/eMAGMA_enrich.R"))
    # source(paste0(Script_directory,Network_type,"/MAGMA_enrich_preprocess.R"))
    eMAGMA_enrich = cmpfun(eMAGMA_enrich)
    # MAGMA_enrich_preprocess = cmpfun(MAGMA_enrich_preprocess)
    pathologies = magma_pathology[magma_pathology %in% c("SCZ","MDD","BIP","ASD","ADHD","OCD","PTSD","SA",
                                                                 "AD","PD","ALS","CUD","RA","CD","UC")]
    if(!"eMAGMA" %in% list.files(Output_directory)){dir.create(paste0(Output_directory,"eMAGMA"))}
    purrr::walk(pathologies,~{
      patho = .x
      eMAGMA_enrich(Net = Network,
                    Net_name = Network_name,
                    genemap = GeneMap,
                    pathology = patho,
                    target.directory = paste0(Target_directory,"eMAGMA/GTEx/"),
                    output.directory = paste0(Output_directory),
                    magma.directory = MAGMA_directory)
      print(paste0("eMAGMA enrichment for ",patho," computed with success"))
    })
  }
}
```
