## Supplementary material for "Dopamine and schizophrenia from bench to bedside: Discovery of a striatal co-expression risk gene set that predicts in vivo measures of striatal function": Compressed zipped directory containing scripts used for this study.: EnrichGO_custom.R

```
#### GO enrichment ####
EnrichGO_custom = function (gene, OrgDb, keyType = "ENTREZID", ont = "MF", pvalueCutoff = 0.05, 
                            pAdjustMethod = "BH", universe, qvalueCutoff = 0.2, minGSSize = 10, 
                            maxGSSize = 500, readable = FALSE, pool = FALSE) 
{
  
  # browser()
          require(clusterProfiler)
          require(org.Hs.eg.db)
          require(DOSE)
          require(enrichplot)
          require(ggplot2)
          require(magrittr)
  # Function modified from clusterProfiler::EnrichGO to reuse GO_DATA from memory instead of recalculating 
          
          ont %<>% toupper
          ont <- match.arg(ont, c("BP", "MF", "CC", "ALL"))
          #GO_DATA <- get_GO_data(OrgDb, ont, keyType)
          G = get(paste0("GO_DATA_ENSEMBL_",ont))
          assign("GO_DATA",G)
          if (missing(universe)) 
                    universe <- NULL
          if (ont == "ALL" && !pool) {
                    lres <- lapply(c("BP", "CC", "MF"), function(ont) suppressMessages(enrichGO(gene, 
                                                                                                OrgDb, keyType, ont, pvalueCutoff, pAdjustMethod, 
                                                                                                universe, qvalueCutoff, minGSSize, maxGSSize)))
                    lres <- lres[!sapply(lres, is.null)]
                    if (length(lres) == 0) 
                              return(NULL)
                    df <- do.call("rbind", lapply(lres, as.data.frame))
                    geneSets <- lres[[1]]@geneSets
                    if (length(lres) > 1) {
                              for (i in 2:length(lres)) {
                                        geneSets <- append(geneSets, lres[[i]]@geneSets)
                              }
                    }
                    res <- lres[[1]]
                    res@result <- df
                    res@geneSets <- geneSets
          }
          else {
                    res <- clusterProfiler:::enricher_internal(gene, pvalueCutoff = pvalueCutoff, 
                                                               pAdjustMethod = pAdjustMethod, universe = universe, 
                                                               qvalueCutoff = qvalueCutoff, minGSSize = minGSSize, 
                                                               maxGSSize = maxGSSize, USER_DATA = GO_DATA)
                    if (is.null(res)) 
                              return(res)
          }
          res@keytype <- keyType
          res@organism <- clusterProfiler:::get_organism(OrgDb)
          if (readable) {
                    res <- clusterProfiler:::setReadable(res, OrgDb)
          }
          res@ontology <- ont
          if (ont == "ALL") {
                    res <- clusterProfiler:::add_GO_Ontology(res, GO_DATA)
          }
          return(res)
}
```
