## Supplementary material for "Dopamine and schizophrenia from bench to bedside: Discovery of a striatal co-expression risk gene set that predicts in vivo measures of striatal function": Compressed zipped directory containing scripts used for this study.: enrichPathway_custom.R

```
enrichPathway_custom = function (gene, organism = "human", 
                                 pvalueCutoff = 0.05, pAdjustMethod = "BH", qvalueCutoff = 0.2, 
                                 universe, minGSSize = 10, maxGSSize = 500, readable = FALSE) 
{
  Reactome_DATA <- get("Reactome_DATA")
  #Reactome_DATA <- get_Reactome_DATA(organism)
  
  res <- ReactomePA:::enricher_internal(gene, pvalueCutoff = pvalueCutoff, 
                           pAdjustMethod = pAdjustMethod, qvalueCutoff = qvalueCutoff, 
                           universe = universe, minGSSize = minGSSize, maxGSSize = maxGSSize, 
                           USER_DATA = Reactome_DATA)
  if (is.null(res)) 
    return(res)
  res@keytype <- "ENTREZID"
  res@organism <- organism
  OrgDb <- ReactomePA:::getDb(organism)
  if (readable) {
    res <- setReadable(res, OrgDb)
  }
  res@ontology <- "Reactome"
  return(res)
}
```
