## Supplementary material for "Dopamine and schizophrenia from bench to bedside: Discovery of a striatal co-expression risk gene set that predicts in vivo measures of striatal function": Compressed zipped directory containing scripts used for this study.: GO_enrich.R

```
###############
###  Main  ####
GO_enrich = function(Net,Net_type,network_name,target_name,geneMap,pvalue,method.adjust = "fdr",script.directory,target.directory,output.directory){
  #browser()
  require(clusterProfiler)
  require(org.Hs.eg.db)
  require(DOSE)
  require(enrichplot)
  require(ggplot2)
  require(magrittr)
  require(purrr)
  require(limma)
  require(dplyr)
  # require(ReactomePA)
  
  source(paste0(script.directory,Net_type,"/EnrichGO_custom.R"))
  source(paste0(script.directory,Net_type,"/enrichPathway_custom.R"))
  load(paste0(target.directory,"GO_DATA_ENSEMBL.RData"),envir = .GlobalEnv)  #Load saved GO_DATA list
  # load(paste0(target.directory,"Reactome_DATA_human.RData"),envir = .GlobalEnv)  #Load saved Reactome_human_DATA list
  
  ll.modules = Net
  universe = unique(geneMap$ensembl)
  
  ll.modules.ENTREZ = map(ll.modules,~{clusterProfiler::bitr(.,fromType = "ENSEMBL",toType = "ENTREZID",OrgDb = org.Hs.eg.db)$ENTREZID})
  universe.ENTREZ = clusterProfiler::bitr(universe,fromType = "ENSEMBL",toType = "ENTREZID",OrgDb = org.Hs.eg.db)$ENTREZID
  
  plot.fn = function(df,ont, module){
    df$Description = factor(df$Description,levels = rev(df$Description))
    print(ggplot(df, aes(x= GeneRatio, y=Description, size = Count, color = p.adjust)) +
            geom_point() + xlab("GeneRatio") +
            scale_color_continuous(low = "red", high = "blue", name = df$p.adjust, guide= guide_colorbar(title = "p.adjust", reverse = T)) +
            ggtitle(module, subtitle = ont) + theme_dose(12) + scale_size(range = c(3, 8), guide = guide_legend(title = "Count"))  + theme(axis.text.y = element_text(size = 10)) + coord_equal(ratio = 0.005))
  }
  
  for (p in pvalue){
    
    ############ Gene Ontology #########
    result_all_modules_BP     = compareCluster(geneClusters = ll.modules, fun = "EnrichGO_custom", universe = universe, ont = "BP", OrgDb = org.Hs.eg.db, keyType = "ENSEMBL", pvalueCutoff = p, qvalueCutoff = p, pAdjustMethod = method.adjust)
    df_all_modules_BP         = fortify(result_all_modules_BP, showCategory = 30, includeAll = F)
    df_all_modules_BP$Cluster = strsplit2(df_all_modules_BP$Cluster, "\\n")[,1]
    missing = c(setdiff(names(Net),unique(df_all_modules_BP$Cluster)))
    
    result_all_modules_MF     = compareCluster(geneClusters = ll.modules, fun = "EnrichGO_custom", universe = universe, ont = "MF", OrgDb = org.Hs.eg.db, keyType = "ENSEMBL", pvalueCutoff = p, qvalueCutoff = p, pAdjustMethod = method.adjust)
    df_all_modules_MF         = fortify(result_all_modules_MF, showCategory = 30, includeAll = F)
    df_all_modules_MF$Cluster = strsplit2(df_all_modules_MF$Cluster, "\\n")[,1]
    missing = c(missing,setdiff(names(Net),unique(df_all_modules_MF$Cluster)))
    
    result_all_modules_CC     = compareCluster(geneClusters = ll.modules, fun = "EnrichGO_custom", universe = universe, ont = "CC", OrgDb = org.Hs.eg.db, keyType = "ENSEMBL", pvalueCutoff = p, qvalueCutoff = p, pAdjustMethod = method.adjust)
    df_all_modules_CC         = fortify(result_all_modules_CC, showCategory = 30, includeAll = F)
    df_all_modules_CC$Cluster = strsplit2(df_all_modules_CC$Cluster, "\\n")[,1]
    missing = unique(c(missing,setdiff(names(Net),unique(df_all_modules_CC$Cluster))))
    
    ll.GO = list(BP = split.data.frame(df_all_modules_BP, df_all_modules_BP$Cluster),
                 MF = split.data.frame(df_all_modules_MF, df_all_modules_MF$Cluster),
                 CC = split.data.frame(df_all_modules_CC, df_all_modules_CC$Cluster))
    if(length(missing)!=0){
      for(i in missing){
        
        ll.GO$BP[[i]] = ll.GO$BP[[1]];ll.GO$MF[[i]] = ll.GO$MF[[1]];ll.GO$CC[[i]] = ll.GO$CC[[1]]
        ll.GO$BP[[i]]$Cluster = rep(i,nrow(ll.GO$BP[[i]]))
        ll.GO$MF[[i]]$Cluster = rep(i,nrow(ll.GO$MF[[i]]))
        ll.GO$CC[[i]]$Cluster = rep(i,nrow(ll.GO$CC[[i]]))
        ll.GO$BP[[i]]$pvalue = ll.GO$BP[[i]]$p.adjust = rep(1,nrow(ll.GO$BP[[i]]))
        ll.GO$MF[[i]]$pvalue = ll.GO$MF[[i]]$p.adjust = rep(1,nrow(ll.GO$MF[[i]]))
        ll.GO$CC[[i]]$pvalue = ll.GO$CC[[i]]$p.adjust = rep(1,nrow(ll.GO$CC[[i]]))
      }}
    ll.GO = lapply(ll.GO,function(x)x[names(Net)])
    ll.GO[["modules"]] = names(Net)
    
    pdf(paste0(output.directory,"/enrich", "_", target_name[1], "_", network_name, ".pdf"),width = 15,  height = 10, useDingbats = FALSE)
    
    pwalk(ll.GO, ~{
      plot.fn(..1, ont = "BP", module = ..4)
      plot.fn(..2, ont = "MF", module = ..4)
      plot.fn(..3, ont = "CC", module = ..4)
    })
    
    # dotplot(result_all_modules_BP, includeAll= T)
    # dotplot(result_all_modules_MF, includeAll= T)
    # dotplot(result_all_modules_CC, includeAll= T)
    
    dev.off()
    
    GO = dplyr::lst(result_all_modules_BP, result_all_modules_MF, result_all_modules_CC)
    save(GO,ll.GO, file = paste0(output.directory,"/enrich", "_", target_name[1], "_", network_name, ".RData"))
    
    ############ KEGG Ontology #########
    result_all_modules_KEGG         = compareCluster(geneClusters = ll.modules.ENTREZ, fun = "enrichKEGG"   , organism = "hsa" , keyType = "kegg", pvalueCutoff = p, qvalueCutoff = p, pAdjustMethod = method.adjust, universe = universe.ENTREZ, minGSSize = 10, maxGSSize = 500)
    df_all_modules_KEGG             = fortify(result_all_modules_KEGG, showCategory = 30, includeAll = F)
    df_all_modules_KEGG$Cluster     = strsplit2(df_all_modules_KEGG$Cluster, "\\n")[,1]
    missing = unique(setdiff(names(Net),df_all_modules_KEGG$Cluster))
    
    ll.KEGG = list(KEGG = split.data.frame(df_all_modules_KEGG, df_all_modules_KEGG$Cluster))
    if(length(missing)!=0){
      for(i in missing){
        
        ll.KEGG$KEGG[[i]] = ll.KEGG$KEGG[[1]]
        ll.KEGG$KEGG[[i]]$Cluster = rep(i,nrow(ll.KEGG$KEGG[[i]]))
        ll.KEGG$KEGG[[i]]$pvalue = ll.KEGG$KEGG[[i]]$p.adjust = rep(1,nrow(ll.KEGG$KEGG[[i]]))
      }}
    ll.KEGG = lapply(ll.KEGG,function(x)x[names(Net)])
    ll.KEGG[["modules"]] = names(Net)
    
    pdf(paste0(output.directory,"/enrich", "_", target_name[2], "_", network_name, ".pdf"),width = 15,  height = 10, useDingbats = FALSE)
    
    pwalk(ll.KEGG, ~{plot.fn(..1, ont = "KEGG", module = ..2)})
    
    #dotplot(result_all_modules_KEGG, includeAll= T)
    
    dev.off()
    
    save(result_all_modules_KEGG,ll.KEGG, file = paste0(output.directory,"/enrich", "_", target_name[2], "_", network_name, ".RData"))
    
    # ############ REACTOME Ontology #########
    # result_all_modules_reactome     = compareCluster(geneClusters = ll.modules.ENTREZ, fun = "enrichPathway_custom",organism = "human"           , pvalueCutoff = p, qvalueCutoff = p, pAdjustMethod = method.adjust, universe = universe.ENTREZ, minGSSize = 10, maxGSSize = 500)
    # df_all_modules_reactome             = fortify(result_all_modules_reactome, showCategory = 30, includeAll = F)
    # df_all_modules_reactome$Cluster     = strsplit2(df_all_modules_reactome$Cluster, "\\n")[,1]
    # missing = unique(setdiff(names(Net),df_all_modules_reactome$Cluster))
    # 
    # ll.Reactome = list(Reactome = split.data.frame(df_all_modules_reactome, df_all_modules_reactome$Cluster))
    # if(length(missing)!=0){
    #           for(i in missing){
    #           
    #           ll.Reactome$Reactome[[i]] = ll.Reactome$Reactome[[1]]
    #           ll.Reactome$Reactome[[i]]$Cluster = rep(i,nrow(ll.Reactome$Reactome[[i]]))
    #           ll.Reactome$Reactome[[i]]$pvalue = ll.Reactome$Reactome[[i]]$p.adjust = rep(1,nrow(ll.Reactome$Reactome[[i]]))
    # }}
    # ll.Reactome = lapply(ll.Reactome,function(x)x[names(Net)])
    # ll.Reactome[["modules"]] = names(Net)
    # 
    # pdf(paste0(output.directory,"/enrich", "_", target_name[3], "_", network_name, ".pdf"),width = 15,  height = 10, useDingbats = FALSE)
    # 
    # pwalk(ll.Reactome, ~{plot.fn(..1, ont = "REACTOME", module = ..2)})
    # 
    # #dotplot(result_all_modules_reactome, includeAll= T)
    # 
    # dev.off()
    # 
    # save(result_all_modules_reactome,ll.Reactome, file = paste0(output.directory,"/enrich", "_", target_name[3], "_", network_name, ".RData"))
    
  }
}
```
