## Supplementary material for "Dopamine and schizophrenia from bench to bedside: Discovery of a striatal co-expression risk gene set that predicts in vivo measures of striatal function": Compressed zipped directory containing scripts used for this study.: H-MAGMA_enrich.R

```
#### H-MAGMA enrich ####
H.MAGMA_enrich = function(Net,Net_name,genemap,target.directory,output.directory,pathology,magma.directory = NULL){
  
  require(future);require(furrr);require(purrr)
  universe = grep("universe.txt",list.files(output.directory),value = T)
  if(pathology == "SCZ") {pathology.file = "PGC3"} else pathology.file = pathology
  patho.file = grep(paste0(pathology.file),list.files(target.directory),value = T)
  patho.file = patho.file[grepl("genes.raw",patho.file)]
  future::plan("multisession")
  ######## Gene-level analysis step ####
  furrr::future_walk(universe,~{
    biotype = .x
    bio = gsub("_universe.txt","",biotype)
    purrr::walk(patho.file,~{
      ### run Magma
      pgc = .x
      annot.type = gsub(paste0("_gene.analysis.genes.raw"),"",pgc)
      magma = "magma"
      gene_results = paste0(target.directory,pgc) ### output of gene analysis already performed
      gene_set = paste0(output.directory,Net_name,"_gene.set") ### gene set info file previously created 
      gene_include = paste0(output.directory,biotype) ### universe file previously created
      out = paste0(output.directory,"H-MAGMA/",Net_name,"_",bio,"_",annot.type,"_gene-level.analysis")
      cmd = paste0(magma.directory,magma," --gene-results ",gene_results," --set-annot ",gene_set," col=2,1 --settings gene-include=",gene_include," --out ",out)
      ##### run analysis 
      system(cmd) 
    })
  })
  future::plan("sequential")
  # rm(list = setdiff(ls(),c("Net","Net_name","genemap","target.directory","output.directory","sumstats")))
  
  ##### organize output file #####
  result.name = grep(".gsa.out",list.files(paste0(output.directory,"H-MAGMA/")),value = T)
  result.name = grep(pathology.file,result.name,value = T)
  PGC_MAGMA_out = sapply(universe,function(biotype){
    
    bio = gsub("_universe.txt","",biotype)
    result = grep(bio,result.name,value = T)
    result = grep(Net_name,result,value = T)
    res.name = gsub(paste0(c("_gene-level.analysis.gsa.out",paste0(Net_name,"_",bio,"_")),collapse = "|"),"",result)
    res = purrr::map(result,function(x){
      
      ### load output files
      PGC_out <- read.table(paste0(output.directory,"H-MAGMA/",x), 
                            header=TRUE, quote="\"", stringsAsFactors=FALSE)
      ### set pvalue = 1 for missing components
      miss_component = setdiff(names(Net),PGC_out$VARIABLE)
      if(length(miss_component)!=0){
        for (i in miss_component){
          miss_comp = PGC_out[1,]
          miss_comp$VARIABLE = i
          miss_comp$P = 1
          PGC_out = rbind(PGC_out,miss_comp)
        }
      } 
      ### adjust with fdr and bonf correction
      PGC_out$P_fdr = p.adjust(PGC_out$P,method = "fdr")
      PGC_out$P_bonf = p.adjust(PGC_out$P,method = "bonferroni")
      ### order dataframe based on components name
      PGC_out$VARIABLE = factor(PGC_out$VARIABLE,levels = names(Net))
      PGC_out = PGC_out[order(PGC_out$VARIABLE),]
    })
    names(res) = res.name
    res[order(names(res))]
  },simplify = F)
  names(PGC_MAGMA_out) = gsub("_universe.txt","",universe)
  ### save ###
  save(PGC_MAGMA_out,file = paste0(output.directory,"H-MAGMA/",Net_name,"_",pathology,".RData"))
  
  ##### plot ####
  require(grDevices)
  purrr::iwalk(PGC_MAGMA_out,function(x,y){
    pdf(paste0(output.directory,"H-MAGMA/",Net_name,"_",y,"_",pathology,"_MAGMA.pdf"),width = 18, useDingbats = FALSE)
    par(mar = c(8,5,5,2), lwd = 96/72, ps = 8, las = 2)
    
    purrr::iwalk(x,function(z,w){
      signif = which(z[,"P_fdr"] < 0.05)
      
      if (length(signif) == 0){
        
        barplot(height = -log10(z[,"P_fdr"]), 
                col = "grey", names.arg = z[,"VARIABLE"], space = 0,
                main = paste0('H-MAGMA enrichment of ', Net_name, 
                              '\n',w),
                ylab = '-log10(fdr-adjusted p-value)', cex.lab = 3, cex.axis = 2, ylim = c(0,max(-log10(z[,"P_fdr"]))+1),las = 2)
        abline(h = -log10(0.05), lwd = 2, lty = 2, col = 'darkred')
        text(x = 5, y = -log10(.05) +.3, labels = 'FDR=0.05', col = 'darkred')
      } else {
        
        bp = barplot(height = -log10(z[,"P_fdr"]), 
                     col = "grey", names.arg = z[,"VARIABLE"], space = 0,
                     main = paste0('H-MAGMA enrichment of ', Net_name, 
                                   '\n',w),
                     ylab = '-log10(fdr-adjusted p-value)', cex.lab = 3, cex.axis = 2, ylim = c(0,max(-log10(z[,"P_fdr"]))+2),las = 2)
        abline(h = -log10(0.05), lwd = 2, lty = 2, col = 'darkred')
        text(x = 5, y = -log10(.05) +.3, labels = 'FDR=0.05', col = 'darkred')      
        text(x = bp[signif], y = -log10(z[signif,"P_fdr"])+1.5, 
             labels = z[signif,"NGENES"], cex = 1.5)
        
      }         
      
    })
    dev.off()
  })
  
  
}
```
