## Supplementary material for "Dopamine and schizophrenia from bench to bedside: Discovery of a striatal co-expression risk gene set that predicts in vivo measures of striatal function": Compressed zipped directory containing scripts used for this study.: H.MAGMA_preprocess.R

```
#### H-MAGMA preprocess ####
require(future);require(furrr);require(purrr)

pathology = c("SA")
windows = c("Adult_brain","Fetal_brain")
target.dir = "script.data.enrichment_new/RData"
purrr::walk(windows,~{
  kbp = .x
  purrr::walk(pathology,function(patho){
    sumstats = grep("check|clumped|grch38|\\.version|\\.RData|PGC2|PGC2\\.clozuk",
                    list.files(paste0("script.data.enrichment_new/RData/",tolower(patho),"/")),value = T,invert = T)   
    purrr::walk(sumstats,~{
      pgc = .x
      ### run Magma
      magma      = "~/Desktop/magma_v1/magma"
      gene_annot = paste0("script.data.enrichment_new/RData/H-MAGMA/",kbp,".genes.annot") ### output of annotation step
      pval       = paste0(target.dir,"/",tolower(patho),"/",pgc," ncol=N") ### summary statistic of PGC
      out        = paste0("script.data.enrichment_new/RData/H-MAGMA/",kbp,"_",pgc,"_gene.analysis")
      
      future::plan("multicore")
      dir.file = furrr::future_map_chr(1:22,~{
        chr = .x
        
        bfile = paste0(target.dir,"/g1000_eur.chr",chr) ### 1000 genome genotype file for LD estimation
        cmd = paste0(magma," --bfile ",bfile," --gene-annot ",gene_annot," --pval ",pval," --batch ",chr," chr --out ",out)
        
        if ((patho == "RA" & chr == 6)|(patho == "UC" & chr == 6)) {
          cmd = paste0(cmd, " --gene-settings adap-permp=10000,10")
        }
        
        
        ##### run analysis
        raw.file = paste0(out, ".batch", chr, "_chr.genes.raw")
        # out.file = paste0(out, ".batch", chr, "_chr.genes.out")
        
        system(cmd)
        return(gsub(".genes.raw$","",raw.file))
        
      })
      future::plan("sequential")
      
      ### merge MAGMA batch output
      cmd.1 = paste0(paste0("{ cat ",dir.file[1],".genes.raw;"),paste0(" sed '1,2d' ",dir.file[-1],collapse = ".genes.raw;"),".genes.raw; } > ",
                     out,".genes.raw")
      cmd.2 = paste0(paste0("{ cat ",dir.file[1],".genes.out;"),paste0(" sed '1d' ",dir.file[-1],collapse = ".genes.out;"),".genes.out; } > ",
                     out,".genes.out")
      
      system(cmd.1)
      print(paste0("Writing file: ",out,".genes.raw"))
      system(cmd.2)
      print(paste0("Writing file: ",out,".genes.out"))
      
    })
  })
})
```
