## Supplementary material for "Dopamine and schizophrenia from bench to bedside: Discovery of a striatal co-expression risk gene set that predicts in vivo measures of striatal function": Compressed zipped directory containing scripts used for this study.: myPermutation.R

```
myPermutation = function(target,components,all_genes,p,perm = 10000){

          require(purrr)
          require(future)
          require(furrr)
          require(Rcpp)
          require(RcppArmadillo)
          #require(sampleCpp)
          future::plan("multicore")
          result = lapply(components,function(x){
                    
                    #### compute real enrichment
                    real.enrich = lapply(target,function(t){
                              purrr::map2(t,list(x),~{
                                        tt = .x ; real = .y
                                        length(intersect(tt,real))})
                    })
                    #### compute null enrichment
                    
                    null.enrich = furrr::future_map(1:perm,~{
                              
                              null.comp = list(sample(all_genes,size = length(x),prob = p,replace = F))
                              lapply(target,function(t){
                                        purrr::map2(t,null.comp,~{
                                                  tt = .x ; null = .y
                                                  length(intersect(tt,null))})
                              })
                    },.options = furrr_options(seed = T))
                    
                    null.enrich = purrr::transpose(null.enrich)
                    null.enrich = lapply(null.enrich,function(n){
                              join = function(list1,list2,...){purrr::map2(list1,list2,c)}
                              Reduce(join,n)
                    })
                    #### obtain empirical pvalue
                    enrich.matrix = purrr::map2(real.enrich,null.enrich,~{
                              
                              real = .x ; null = .y
                              purrr::map2(real,null,~{
                                        rr = .x ; nn = .y
                                        (sum(nn >= rr) + 1)/(perm + 1)
                              })
                              
                    })
                    return(dplyr::lst(real.enrich,enrich.matrix))
          })
          future::plan("sequential")
          result = purrr::transpose(result)
          result = lapply(result,function(x){
                    new = purrr::transpose(x)
                    lapply(new,function(y)t(sapply(y,unlist)))
          })
          return(result)
          
}
```
