## Supplementary material for "Dopamine and schizophrenia from bench to bedside: Discovery of a striatal co-expression risk gene set that predicts in vivo measures of striatal function": Compressed zipped directory containing scripts used for this study.: C18_assoc.R

```
###############################################################################################################
############################################### GTEx C18 gene expression association ##########################
###############################################################################################################

load("data.SDA.RData")
load("sda_preprocessing.RData")
CN = matrix_SDA_final$matrix_Blom$caudate
colnames(CN) = gene_common$hgnc[match(colnames(matrix_SDA_final$matrix_log2$caudate),gene_common$gencodeID)]
row.names(CN) = sample_list$caudate$SAMPID[match(row.names(matrix_SDA_final$matrix_log2$caudate),sample_list$caudate$SAMPID)]
DLPFC = matrix_SDA_final$matrix_Blom$dlpfc
colnames(DLPFC) = gene_common$hgnc[match(colnames(matrix_SDA_final$matrix_log2$dlpfc),gene_common$gencodeID)]
row.names(DLPFC) = sample_list$dlpfc$SAMPID[match(row.names(matrix_SDA_final$matrix_log2$dlpfc),sample_list$dlpfc$SAMPID)]
HP = matrix_SDA_final$matrix_Blom$hippo
colnames(HP) = gene_common$hgnc[match(colnames(matrix_SDA_final$matrix_log2$hippo),gene_common$gencodeID)]
row.names(HP) = sample_list$hippo$SAMPID[match(row.names(matrix_SDA_final$matrix_log2$hippo),sample_list$hippo$SAMPID)]

CN = CN[,c("DRD1","DRD2","TH","DDC")]
DLPFC = DLPFC[,c("DRD1","DRD2","TH","DDC")]
HP = HP[,c("DRD1","DRD2","TH","DDC")]

data[,paste0(c("DRD1","DRD2","TH","DDC"),"_CN")] = CN[match(data$SAMPID,row.names(CN)),1:4]
data[,paste0(c("DRD1","DRD2","TH","DDC"),"_DLPFC")] = DLPFC[match(data$SAMPID,row.names(DLPFC)),1:4]
data[,paste0(c("DRD1","DRD2","TH","DDC"),"_HP")] = HP[match(data$SAMPID,row.names(HP)),1:4]
data = data[complete.cases(data),]

mod = as.formula(paste0("V18~DRD2_CN+DRD2_DLPFC+DRD2_HP+DDC_CN+DDC_DLPFC+DDC_HP+TH_CN+TH_DLPFC+TH_HP+Age+Sex+PMI+
RIN_caudate+TotalAssignedGene_caudate+rRNA_rate_caudate+
                        RIN_dlpfc+TotalAssignedGene_dlpfc+rRNA_rate_dlpfc+
                        RIN_hippo+TotalAssignedGene_hippo+rRNA_rate_hippo"))

############# Tx 
require(data.table)
require(SummarizedExperiment)
library(preprocessCore)
library(RNOmni)
load("caudate_brainseq_phase3_hg38_rseTx_merged_n464.rda")
Tx_map = data.frame(rowData(rse_tx))
load("transcriptsID.RData")
load("GTEx_transcripts.RData")

############### Tx caudate
Tx.sample = sample_list$caudate
stopifnot(identical(Tx.sample$SAMPID,row.names(caudate.Tx)))
caudate.Tx = caudate.Tx[,colnames(caudate.Tx) %in% transcriptsID$transcript_id[transcriptsID$gene_id %in% gene_common$gencodeID]]
############# Tx dlpfc
Tx.sample = sample_list$dlpfc
stopifnot(identical(Tx.sample$SAMPID,row.names(dlpfc.Tx)))
dlpfc.Tx = dlpfc.Tx[,colnames(dlpfc.Tx) %in% transcriptsID$transcript_id[transcriptsID$gene_id %in% gene_common$gencodeID]]
############# Tx hippo
Tx.sample = sample_list$hippo
stopifnot(identical(Tx.sample$SAMPID,row.names(hippo.Tx)))
hippo.Tx = hippo.Tx[,colnames(hippo.Tx) %in% transcriptsID$transcript_id[transcriptsID$gene_id %in% gene_common$gencodeID]]

matrix.final = sapply(c("caudate","dlpfc","hippo"),function(x){
  df = get(paste0(x,".Tx"))
  df = df[match(data$SAMPID,row.names(df)),]
  median_filtered = apply(df,2,median)
  check = names(median_filtered)[median_filtered == 0]
  # check = setdiff(check,"DRD2-001")
  matrix = df[,!colnames(df) %in% check]
  
},USE.NAMES = T,simplify = F)

# common = Reduce(intersect,sapply(matrix.final,colnames))

matrix.final = lapply(matrix.final,function(x){
  # x = x[,common]
  #### log2 transformation
  matrix_log =  log2(x + 1)
  #### quantile normalization in each tissue
  matrix_quantile = t(normalize.quantiles(t(matrix_log)))
  #### rank transformation (Blom)
  matrix_blom =  apply(matrix_quantile,2,RankNorm)
  colnames(matrix_blom) = colnames(matrix_quantile) = colnames(matrix_log)
  row.names(matrix_blom) = row.names(matrix_quantile) = row.names(matrix_log)
  return(dplyr::lst(matrix_log,matrix_quantile,matrix_blom))
  
})

matrix.final = lapply(matrix.final,function(x){
  lapply(x,function(y){
    y = y[,colnames(y) %in% Tx_map$transcript_id[Tx_map$gene_name %in% c("DRD2","TH","DDC")]]
    colnames(y) = Tx_map$transcript_name[match(colnames(y),Tx_map$transcript_id)]
    y
  })
})

data[,paste0(grep("^TH\\-|DRD2|^DDC\\-",colnames(matrix.final$caudate$matrix_blom),value = T),"_CN")] = 
  matrix.final$caudate$matrix_blom[match(data$SAMPID,row.names(matrix.final$caudate$matrix_blom)),
                                   grep("^TH\\-|DRD2|^DDC\\-",colnames(matrix.final$caudate$matrix_blom))]

data[,paste0(grep("^TH\\-|DRD2|^DDC\\-",colnames(matrix.final$dlpfc$matrix_blom),value = T),"_DLPFC")] = 
  matrix.final$dlpfc$matrix_blom[match(data$SAMPID,row.names(matrix.final$dlpfc$matrix_blom)),
                                 grep("^TH\\-|DRD2|^DDC\\-",colnames(matrix.final$dlpfc$matrix_blom))]

data[,paste0(grep("^TH\\-|DRD2|^DDC\\-",colnames(matrix.final$hippo$matrix_blom),value = T),"_HP")] = 
  matrix.final$hippo$matrix_blom[match(data$SAMPID,row.names(matrix.final$hippo$matrix_blom)),
                                 grep("^TH\\-|DRD2|^DDC\\-",colnames(matrix.final$hippo$matrix_blom))]


grep("^TH|DRD2|^DDC",colnames(data),value = T)

############
colnames(data) = gsub("\\-","_",colnames(data))

mod = as.formula(paste0("V18~DRD2_003_CN+DRD2_003_DLPFC+DRD2_003_HP+DRD2_001_CN+
DDC_CN+DDC_DLPFC+DDC_HP+
TH_CN+TH_DLPFC+TH_HP+
Age+Sex+PMI+
RIN_caudate+TotalAssignedGene_caudate+rRNA_rate_caudate+
                        RIN_dlpfc+TotalAssignedGene_dlpfc+rRNA_rate_dlpfc+
                        RIN_hippo+TotalAssignedGene_hippo+rRNA_rate_hippo"))
fit = lm(mod,data = data)
```
