## Supplementary material for "Dopamine and schizophrenia from bench to bedside: Discovery of a striatal co-expression risk gene set that predicts in vivo measures of striatal function": Compressed zipped directory containing scripts used for this study.: C80_assoc.R

```
###############################################################################################################
############################################### LIBD C80 gene expression association ##########################
###############################################################################################################

load("data.SDA.RData")
load("sda_preprocessing.RData")
CN = matrix_SDA_final$matrix_Blom$caudate$all
colnames(CN) = gene_common$ALL$hgnc[match(colnames(matrix_SDA_final$matrix_log2$caudate$all),gene_common$ALL$gencodeID)]
row.names(CN) = sample_list$caudate$all$BrNum[match(row.names(matrix_SDA_final$matrix_log2$caudate$all),sample_list$caudate$all$RNum)]
DLPFC = matrix_SDA_final$matrix_Blom$dlpfc$all
colnames(DLPFC) = gene_common$ALL$hgnc[match(colnames(matrix_SDA_final$matrix_log2$dlpfc$all),gene_common$ALL$gencodeID)]
row.names(DLPFC) = sample_list$dlpfc$all$BrNum[match(row.names(matrix_SDA_final$matrix_log2$dlpfc$all),sample_list$dlpfc$all$RNum)]
HP = matrix_SDA_final$matrix_Blom$hippo$all
colnames(HP) = gene_common$ALL$hgnc[match(colnames(matrix_SDA_final$matrix_log2$hippo$all),gene_common$ALL$gencodeID)]
row.names(HP) = sample_list$hippo$all$BrNum[match(row.names(matrix_SDA_final$matrix_log2$hippo$all),sample_list$hippo$all$RNum)]

CN = CN[,c("DRD1","DRD2","TH","DDC")]
DLPFC = DLPFC[,c("DRD1","DRD2","TH","DDC")]
HP = HP[,c("DRD1","DRD2","TH","DDC")]

data[,paste0(c("DRD1","DRD2","TH","DDC"),"_CN")] = CN[match(data$BrNum,row.names(CN)),1:4]
data[,paste0(c("DRD1","DRD2","TH","DDC"),"_DLPFC")] = DLPFC[match(data$BrNum,row.names(DLPFC)),1:4]
data[,paste0(c("DRD1","DRD2","TH","DDC"),"_HP")] = HP[match(data$BrNum,row.names(HP)),1:4]
data = data[complete.cases(data),]
adults <- data[data$Dx %in% "Control",]
adults <- data[data$Age >=20 & data$Dx %in% "Control",]

mod = as.formula(paste0("V80~Dx*DRD2_CN+Dx*DRD2_DLPFC+Dx*DRD2_HP+Dx*DDC_CN+Dx*DDC_DLPFC+Dx*DDC_HP+Dx*TH_CN+Dx*TH_DLPFC+Dx*TH_HP+Dx*Age+Race+Sex+PMI+
RIN_caudate+totalAssignedGene_caudate+rRNA_rate_caudate+mitoRate_caudate+
                        RIN_dlpfc+totalAssignedGene_dlpfc+rRNA_rate_dlpfc+mitoRate_dlpfc+
                        RIN_hippo+totalAssignedGene_hippo+rRNA_rate_hippo+mitoRate_hippo"))
fit = lm(mod,data = data)

mod = as.formula(paste0("V80~DRD2_CN+DRD2_DLPFC+DRD2_HP+DDC_CN+DDC_DLPFC+DDC_HP+TH_CN+TH_DLPFC+TH_HP+Age+Race+Sex+PMI+
RIN_caudate+totalAssignedGene_caudate+rRNA_rate_caudate+mitoRate_caudate+
                        RIN_dlpfc+totalAssignedGene_dlpfc+rRNA_rate_dlpfc+mitoRate_dlpfc+
                        RIN_hippo+totalAssignedGene_hippo+rRNA_rate_hippo+mitoRate_hippo"))
fit = lm(mod,data = adults)


############# Tx 
library(qs)
library(SummarizedExperiment)
library(preprocessCore)
library(RNOmni)
Tx = qread("rse.TX.qs")
Tx.sample = data.frame(colData(Tx))
Tx.map = data.frame(rowData(Tx))
Tx.matrix = assays(Tx)$tpm
stopifnot(identical(row.names(Tx.matrix),row.names(Tx.map)))
stopifnot(identical(colnames(Tx.matrix),row.names(Tx.sample)))
Tx.map = Tx.map[Tx.map$gene_id %in% gene_common$ALL$gencodeID,]
rm(Tx)

Tx.sample.caudate = Tx.sample[Tx.sample$Region %in% "Caudate" & Tx.sample$BrNum %in% data$BrNum,]
Tx.sample.dlpfc.phase1 = Tx.sample[Tx.sample$Region %in% "DLPFC" & Tx.sample$BrNum %in% data$BrNum & Tx.sample$Dataset %in% "BrainSeq_Phase1",]
Tx.sample.dlpfc.phase2 = Tx.sample[Tx.sample$Region %in% "DLPFC" & Tx.sample$BrNum %in% data$BrNum & Tx.sample$Dataset %in% "BrainSeq_Phase2_DLPFC",]
Tx.sample.hippo = Tx.sample[Tx.sample$Region %in% "HIPPO" & Tx.sample$BrNum %in% data$BrNum,]
common = Reduce(intersect,list(Tx.sample.caudate$BrNum,Tx.sample.dlpfc.phase2$BrNum,Tx.sample.hippo$BrNum))

##### caudate
caudate_Tx = Tx.matrix[row.names(Tx.matrix) %in% row.names(Tx.map),
                       colnames(Tx.matrix) %in% row.names(Tx.sample.caudate)]
row.names(caudate_Tx) = Tx.map$transcript_name
colnames(caudate_Tx) = Tx.sample.caudate$BrNum[match(colnames(caudate_Tx),row.names(Tx.sample.caudate))]
caudate_Tx = t(caudate_Tx[,common])

##### dlpfc
dlpfc_Tx = Tx.matrix[row.names(Tx.matrix) %in% row.names(Tx.map),
                       colnames(Tx.matrix) %in% row.names(Tx.sample.dlpfc.phase2)]
row.names(dlpfc_Tx) = Tx.map$transcript_name
colnames(dlpfc_Tx) = Tx.sample.dlpfc.phase2$BrNum[match(colnames(dlpfc_Tx),row.names(Tx.sample.dlpfc.phase2))]
dlpfc_Tx = t(dlpfc_Tx[,common])

##### hippo
hippo_Tx = Tx.matrix[row.names(Tx.matrix) %in% row.names(Tx.map),
                       colnames(Tx.matrix) %in% row.names(Tx.sample.hippo)]
row.names(hippo_Tx) = Tx.map$transcript_name
colnames(hippo_Tx) = Tx.sample.hippo$BrNum[match(colnames(hippo_Tx),row.names(Tx.sample.hippo))]
hippo_Tx = t(hippo_Tx[,common])

all(common %in% Reduce(intersect,list(row.names(caudate_Tx),row.names(hippo_Tx),row.names(dlpfc_Tx))))

matrix.final = sapply(c("caudate","dlpfc","hippo"),function(x){
  df = get(paste0(x,"_Tx"))
  # df = df[match(data$BrNum,row.names(df)),]
  median_filtered = apply(df,2,median)
  check = names(median_filtered)[median_filtered == 0]
  # check = setdiff(check,"DRD2-001")
  matrix = df[,!colnames(df) %in% check]
  
},USE.NAMES = T,simplify = F)

# common = Reduce(intersect,sapply(matrix.final,colnames))

matrix.final = lapply(matrix.final,function(x){
  # x = x[,common]
  #### log2 transformation
  matrix_log =  log2(x + 1)
  #### quantile normalization in each tissue
  matrix_quantile = t(normalize.quantiles(t(matrix_log)))
  #### rank transformation (Blom)
  matrix_blom =  apply(matrix_quantile,2,RankNorm)
  colnames(matrix_blom) = colnames(matrix_quantile) = colnames(matrix_log)
  row.names(matrix_blom) = row.names(matrix_quantile) = row.names(matrix_log)
  return(dplyr::lst(matrix_log,matrix_quantile,matrix_blom))
  
})

data = data[match(common,data$BrNum),]
data[,paste0(grep("^TH\\-|DRD2|^DDC\\-",colnames(matrix.final$caudate$matrix_blom),value = T),"_CN")] = 
  matrix.final$caudate$matrix_blom[match(data$BrNum,row.names(matrix.final$caudate$matrix_blom)),
                                   grep("^TH\\-|DRD2|^DDC\\-",colnames(matrix.final$caudate$matrix_blom))]

data[,paste0(grep("^TH\\-|DRD2|^DDC\\-",colnames(matrix.final$dlpfc$matrix_blom),value = T),"_DLPFC")] = 
  matrix.final$dlpfc$matrix_blom[match(data$BrNum,row.names(matrix.final$dlpfc$matrix_blom)),
                                   grep("^TH\\-|DRD2|^DDC\\-",colnames(matrix.final$dlpfc$matrix_blom))]

data[,paste0(grep("^TH\\-|DRD2|^DDC\\-",colnames(matrix.final$hippo$matrix_blom),value = T),"_HP")] = 
  matrix.final$hippo$matrix_blom[match(data$BrNum,row.names(matrix.final$hippo$matrix_blom)),
                                   grep("^TH\\-|DRD2|^DDC\\-",colnames(matrix.final$hippo$matrix_blom))]


grep("^TH|DRD2|^DDC",colnames(data),value = T)

###############
colnames(data) = gsub("\\-","_",colnames(data))
data$DRD2s.DRD2l_CN = data$DRD2_003_CN/data$DRD2_001_CN
adults <- data[data$Dx %in% "Control",]
adults <- data[data$Age >=20 & data$Dx %in% "Control",]

mod = as.formula(paste0("V80~Dx*DRD2_003_CN+Dx*DRD2_003_DLPFC+Dx*DRD2_003_HP+Dx*DRD2_001_CN+
Dx*DDC_CN+Dx*DDC_DLPFC+Dx*DDC_HP+
Dx*TH_CN+Dx*TH_DLPFC+Dx*TH_HP+
Dx*Age+Race+Sex+PMI+
RIN_caudate+totalAssignedGene_caudate+rRNA_rate_caudate+mitoRate_caudate+
                        RIN_dlpfc+totalAssignedGene_dlpfc+rRNA_rate_dlpfc+mitoRate_dlpfc+
                        RIN_hippo+totalAssignedGene_hippo+rRNA_rate_hippo+mitoRate_hippo"))
fit = lm(mod,data = data)

mod = as.formula(paste0("V80~DRD2_003_CN+DRD2_003_DLPFC+DRD2_003_HP+DRD2_001_CN+
DDC_CN+DDC_DLPFC+DDC_HP+
TH_CN+TH_DLPFC+TH_HP+
Age+Race+Sex+PMI+
RIN_caudate+totalAssignedGene_caudate+rRNA_rate_caudate+mitoRate_caudate+
                        RIN_dlpfc+totalAssignedGene_dlpfc+rRNA_rate_dlpfc+mitoRate_dlpfc+
                        RIN_hippo+totalAssignedGene_hippo+rRNA_rate_hippo+mitoRate_hippo"))
fit = lm(mod,data = adults)
```
