## Supplementary material for "Dopamine and schizophrenia from bench to bedside: Discovery of a striatal co-expression risk gene set that predicts in vivo measures of striatal function": Compressed zipped directory containing scripts used for this study.: MDS_ancestry_preprocess.R

```
###############################################################################################################
############################################### Ancestry score pre-processing ##################################
###############################################################################################################

HM3_b37 = read.delim("HM3_b37.bim", header=FALSE)
HM3_b37$chr.pos = paste0(HM3_b37$V1,":",HM3_b37$V4)
HM3_b37 = HM3_b37[!duplicated(HM3_b37$chr.pos),]

datasets = c("KCL")
purrr::walk(datasets,function(dname){
  print(dname)
  bim = grep("\\.bim",list.files(paste0("ENIGMA_GE/",dname)),value = T)
  plink = "~/Desktop/plink/plink2"
  data.dir = paste0("ENIGMA_GE/",dname,"/")
  data.name = gsub("\\.bim","",bim)
  out.dir = data.dir
  out.name = paste0(dname,".maf0.01.geno0.05")
  cmd = paste0(plink," --bfile ",data.dir,data.name, " --chr 1-22 --snps-only --rm-dup force-first --maf 0.01 --geno 0.05 --hwe 1e-06 --make-bed --out ",out.dir,out.name)
  system(cmd)
  system(paste0("rm ",data.dir,"*~"))
  ############################################################################ filter MAF 0.01 and geno 0.05 and hwe 1e-06
  bim = grep("\\maf0.01\\.geno0.05\\.bim",list.files(paste0("ENIGMA_GE/",dname)),value = T)
  df = read.delim(paste0("ENIGMA_GE/",dname,"/",bim), header=FALSE)
  df$chr.pos = paste0(df$V1,":",df$V4)
  df = df[!duplicated(df$chr.pos),]
  ambig = which((df$V5 == "T" & df$V6 == "A") |
                  (df$V5=="A" & df$V6=="T")|
                  (df$V5=="C" &  df$V6=="G")|
                  (df$V5=="G" & df$V6=="C"))
  if(length(ambig)!=0) df = df[-ambig,]
  
  common = intersect(df$chr.pos,HM3_b37$chr.pos)
  df = df[match(common,df$chr.pos),]
  df$new.rs = HM3_b37$V2[match(common,HM3_b37$chr.pos)]
  
  cat(df$V2,sep = "\n",file = paste0("ENIGMA_GE/",dname,"/to.extract.",dname))
  write.table(df[,c("V2","new.rs")],row.names = F,col.names = F,quote = F,sep = "\t", file = paste0("ENIGMA_GE/",dname,"/to.update.",dname))
  cat(df$new.rs,sep = "\n",file = paste0("ENIGMA_GE/",dname,"/to.extract.HM3.",dname))
  
  ################################ plink steps ###############################
  plink = "~/Desktop/plink/plink2"
  data.dir = paste0("ENIGMA_GE/",dname,"/")
  data.name = paste0(dname,".maf0.01.geno0.05")
  to.extract = paste0("to.extract.",dname)
  out.dir = data.dir
  out.name = paste0(dname,"_updated")
  cmd = paste0(plink," --bfile ",data.dir,data.name, " --extract ",data.dir,to.extract, " --make-bed --out ",out.dir,out.name)
  system(cmd)
  system(paste0("rm ",data.dir,"*~"))
  ############################################################################ extract common SNP from target
  data.name = out.name
  to.update = paste0("to.update.",dname)
  cmd = paste0(plink," --bfile ",data.dir,data.name, " --update-name ",data.dir,to.update, " --make-bed --out ",out.dir,out.name)
  system(cmd)
  system(paste0("rm ",data.dir,"*~"))
  ############################################################################ update correct SNP names
  data.dir = "ENIGMA_GE/HM3/"
  data.name = "HM3_b37"
  to.extract = paste0("ENIGMA_GE/",dname,"/to.extract.HM3.",dname)
  out.dir = paste0("ENIGMA_GE/",dname,"/")
  out.name = paste0(data.name,".",dname)
  cmd = paste0(plink," --bfile ",data.dir,data.name, " --extract ",to.extract, " --make-bed --out ",out.dir,out.name)
  system(cmd)
  ############################################################################ extract common SNP from reference
  plink = "~/Desktop/plink/plink"
  data.dir = paste0("ENIGMA_GE/",dname,"/")
  data.name = paste0(dname,"_updated")
  to.merge = paste0(data.dir,"HM3_b37.",dname,c(".bed",".bim",".fam"),collapse = " ")
  out.dir = "ENIGMA_GE/HM3.merged/"
  out.name = paste0("HM3.",dname,".merged")
  cmd = paste0(plink, " --bfile ",data.dir,data.name, " --bmerge ",to.merge, " --keep-allele-order --allow-no-sex --make-bed --out ",out.dir,out.name )
  system(cmd)
  ############################################################################ merge target and reference
  if(sum(grepl(paste0("HM3\\.",dname,"\\.merged\\-merge\\.missnp"),list.files(out.dir))) != 0){
    out.dir = paste0("ENIGMA_GE/",dname,"/")
    out.name = paste0(dname,"_updated.flipped")
    to.flip = paste0("ENIGMA_GE/HM3.merged/HM3.",dname,".merged-merge.missnp")
    cmd = paste0(plink, " --bfile ",data.dir,data.name, " --flip ",to.flip, " --keep-allele-order --allow-no-sex --make-bed --out ",out.dir,out.name )
    system(cmd)
    system(paste0("rm ",to.flip))
    ########################################################################## flip mismatching SNPs
    data.dir = paste0("ENIGMA_GE/",dname,"/")
    data.name = paste0(dname,"_updated.flipped")
    to.merge = paste0(data.dir,"HM3_b37.",dname,c(".bed",".bim",".fam"),collapse = " ")
    out.dir = "ENIGMA_GE/HM3.merged/"
    out.name = paste0("HM3.",dname,".merged")
    cmd = paste0(plink, " --bfile ",data.dir,data.name, " --bmerge ",to.merge, " --keep-allele-order --allow-no-sex --make-bed --out ",out.dir,out.name )
    system(cmd)
  }
  ############################################################################ redo merging
  if(sum(grepl(paste0("HM3\\.",dname,"\\.merged\\-merge\\.missnp"),list.files(out.dir))) != 0){  
    for(data in c(paste0(dname,"_updated.flipped"),paste0("HM3_b37.",dname))){
      
      data.name = data
      to.exclude = paste0("ENIGMA_GE/HM3.merged/HM3.",dname,".merged-merge.missnp")
      out.dir = data.dir
      out.name = data
      cmd = paste0(plink," --bfile ",data.dir,data.name, " --keep-allele-order --exclude ",to.exclude, " --make-bed --out ",out.dir,out.name)
      system(cmd)
      system(paste0("rm ",data.dir,"*~"))
    }
    ############################################################################ remove mismatch SNPs
    data.dir = paste0("ENIGMA_GE/",dname,"/")
    data.name = paste0(dname,"_updated.flipped")
    to.merge = paste0(data.dir,"HM3_b37.",dname,c(".bed",".bim",".fam"),collapse = " ")
    out.dir = "ENIGMA_GE/HM3.merged/"
    out.name = paste0("HM3.",dname,".merged")
    cmd = paste0(plink, " --bfile ",data.dir,data.name, " --bmerge ",to.merge, " --keep-allele-order --allow-no-sex --make-bed --out ",out.dir,out.name )
    system(cmd)
  }
  ############################################################################ redo merging
  plink = "~/Desktop/plink/plink2"
  data.dir = paste0("ENIGMA_GE/",dname,"/")
  data.name = paste0(dname,"_updated.flipped")
  inversion = "ENIGMA_GE/inversion.txt"
  out.dir =  paste0("ENIGMA_GE/",dname,"/")
  out.name = data.name
  cmd = paste0(plink, " --bfile ",data.dir,data.name, " --exclude range ",inversion, " --indep-pairwise 200 100 0.2 --out ",out.dir,out.name )
  system(cmd)
  ############################################################################ perform pruning
  plink = "~/Desktop/plink/plink"
  fam = read.delim(paste0("ENIGMA_GE/HM3.merged/HM3.",dname,".merged.fam"), header=FALSE,sep = " ")
  eigen = nrow(fam)
  data.dir = "ENIGMA_GE/HM3.merged/"
  data.name = paste0("HM3.",dname,".merged")
  prune.in = paste0("ENIGMA_GE/",dname,"/",dname,"_updated.flipped.prune.in")
  out.dir = "ENIGMA_GE/PCA/"
  out.name = paste0("HM3.",dname,".mds")
  cmd = paste0(plink, " --bfile ",data.dir,data.name, " --extract ",prune.in, " --cluster --mds-plot ",eigen," eigvals --out ",out.dir,out.name )
  system(cmd)
  ############################################################################ perform PCA
})
```
