## Supplementary material for "Dopamine and schizophrenia from bench to bedside: Discovery of a striatal co-expression risk gene set that predicts in vivo measures of striatal function": Compressed zipped directory containing scripts used for this study.: MDS_ancestry_score.R

```
###############################################################################################################
############################################### Ancestry score computation ####################################
###############################################################################################################

library(glmnet)
require(ggplot2)
samples = c("KCL")
purrr::walk(samples,function(sample){
  mds.cluster <- read.csv(paste0("ENIGMA_GE/PCA/HM3.",sample,".mds.mds"), sep="")
  mds.cluster = mds.cluster[,c(1:3,grep(paste0("C",1:20,"$",collapse = "|"),colnames(mds.cluster)))]
  populations = c("CEU",
                  "CHB",
                  "YRI",
                  "TSI",
                  "JPT",
                  "CHD",
                  "MEX",
                  "GIH",
                  "ASW","LWK",
                  "MKK")
  super.populations = c("AFR","EAS","EUR","SAS","AMR")
  mds.cluster$Ethnicity = factor(sapply(mds.cluster$FID,function(x) {
    
    if(x %in% populations){ populations[populations == x]
      
    } else sample
    
    
  }),levels = c(sample,populations))
  mds.cluster$populations = factor(sapply(mds.cluster$FID,function(x) {
    
    if(x %in% c("YRI","LWK","ASW","MKK")){ super.populations[1]
    } else if(x %in% c("CHB","JPT","CHD")){ super.populations[2]
    } else if(x %in% c("CEU","TSI")){ super.populations[3]
    } else if(x %in% c("GIH")){ super.populations[4]
    } else if(x %in% c("MEX")){ super.populations[5]
    } else sample
    
    
  }),levels = c(sample,super.populations))
  mds.cluster$alpha = sapply(mds.cluster$Ethnicity,function(x) {
    
    if(x %in% sample) 1 else 0.5 
    
  })
  
  HM3 = mds.cluster[!mds.cluster$populations %in% sample,]
  mySample = mds.cluster[mds.cluster$populations %in% sample,]
  
  pop.score = sapply(super.populations,function(super_pop){
    
    HM3$score = as.factor(ifelse(HM3$populations %in% super_pop,1,0))
    fit = cv.glmnet(as.matrix(HM3[, paste0("C", 1:20)]), HM3$score, family="binomial");
    score = predict(fit, newx=as.matrix(mySample[, paste0("C", 1:20)]), type="response")
    assign(paste0(super_pop,"_score"),score)
    return(get(paste0(super_pop,"_score")))
  })
  pop.score = as.data.frame(pop.score)
  pop.score[,c("FID","IID")] = mySample[,1:2]
  to.remove = mySample[signif(pop.score$EUR,2) < 0.9,1:2]
  
  save(pop.score,file = paste0(sample,".pop.score.RData"))
  save(to.remove,file = paste0("to.remove.",sample,".RData"))

})
```
