## Supplementary material for "Dopamine and schizophrenia from bench to bedside: Discovery of a striatal co-expression risk gene set that predicts in vivo measures of striatal function": Compressed zipped directory containing scripts used for this study.: C80.heatmap_plot.R

```
library(limma)
library(gtools)
library(stringr)
library(ggplot2)
library(WGCNA)
library(dplyr)
library(purrr)
library(tidyr)
library(ggrepel)
library(patchwork)
library(ComplexHeatmap)
library(circlize)
library(gridtext)
library(magrittr)
library(metap)
library(simplifyEnrichment)

load("sda_quantile/ALL/seed_400/PIP_0.5/filtered_new_method.0.4corr/genes_components_overlap.RData")
comps = names(geneList)
rm(list=setdiff(ls(),"comps"))
dir = "sda_quantile/Heatmap_data/"

DEG.DLPFC_bins    = c("Fromer_nosva"                 ,"Fromer_sva"    ,
                      "Jaffe_DLPFC_sczd"             ,"Jaffe_sczd_2018" )
DEG.HIPPO_bins    = c("Jaffe_HIPPO_sczd")
DEG.CAUDATE_bins  = c("Apua_CAUDATE_sczd" )
DMG_bins      = c("DMGs_Hannon","DMGs_Jaffe","DMGs_Kinoshita",
                  "DMGs_Montano","DMGs_Numata","DMGs_Wockner")
LOF_bins          = "LoF"

all.files = list.files(dir)
patho.files      = all.files[grep("SCZ|MDD|ASD|ADHD|BIP|PTSD|OCD|SA|CD|UC",all.files)]
DEG.file         = all.files[grep("DEG"             ,all.files)]
DMG.file         = all.files[grep("DMG"             ,all.files)]
LOF.file         = all.files[grep("Lof"             ,all.files)]
MAGMA.files      = patho.files[grep("_MAGMA"        ,patho.files)]
HMAGMA.files     = patho.files[grep("H-MAGMA"       ,patho.files)]
patho.files      = patho.files[grep("MAGMA"         ,patho.files,invert = T)]

####
patho.results = patho.files %>% set_names(strsplit2(.,"\\.LIBD")[,1]) %>%
  imap_dfc(~{
    ne = new.env()
    print(.y)
    load(paste0(dir,.x),envir = ne)
    zz = ne$enrich.matrix.pval[,"PGC"]
    zz1 = -log10(zz)
    zz1[zz1 < -log10(0.05)] = 0
    zz1
    # sig.wins = data.frame(rowSums(zz<0.05)) %>% set_colnames(.y)
  }) %>% set_colnames(gsub("all.biotypes","AB",gsub("protein.coding","PC",colnames(.))))

patho.results = patho.results[, paste0(c("SCZ","MDD","BIP", "SA", "ASD","ADHD", "OCD","PTSD","CD","UC"),c(".AB"))]
colnames(patho.results) = gsub("\\.AB","",colnames(patho.results))
colnames(patho.results)[colnames(patho.results) == "BIP"] = "BD"
row.names(patho.results) = comps
# patho.results = as.data.frame(patho.results["V80",])

####
DEGS.DLPFC.result = list('DEGs DLPFC' = DEG.file)  %>%
  imap_dfc(~{
    ne = new.env()
    print(.y)
    load(paste0(dir,.x),envir = ne)
    zz = ne$enrich.matrix.pval[,DEG.DLPFC_bins]
    zz1 = apply(zz,1,function(v) {
      -log10(metap::sumlog(v)$p)
    })
    zz1[zz1 < -log10(0.05)] = 0
    out = data.frame(zz1) %>% set_colnames(.y)
  })  
DEGS.DLPFC.result = as.data.frame(DEGS.DLPFC.result["V80",])

DEGS.HIPPO.result = list('DEGs HP' = DEG.file)  %>%
  imap_dfc(~{
    ne = new.env()
    print(.y)
    load(paste0(dir,.x),envir = ne)
    zz = ne$enrich.matrix.pval[,DEG.HIPPO_bins]
    zz1 = -log10(zz)
    zz1[zz1 < -log10(0.05)] = 0
    out = data.frame(zz1) %>% set_colnames(.y)
  })  
DEGS.HIPPO.result = as.data.frame(DEGS.HIPPO.result["V80",])

DEGS.CAUDATE.result = list('DEGs CN' = DEG.file)  %>%
  imap_dfc(~{
    ne = new.env()
    print(.y)
    load(paste0(dir,.x),envir = ne)
    zz = ne$enrich.matrix.pval[,DEG.CAUDATE_bins]
    zz1 = -log10(zz)
    zz1[zz1 < -log10(0.05)] = 0
    out = data.frame(zz1) %>% set_colnames(.y)
  }) 
DEGS.CAUDATE.result = as.data.frame(DEGS.CAUDATE.result["V80",])

DMGS.SCZ.result = list('DMG SCZ' = DMG.file)  %>%
  imap_dfc(~{
    ne = new.env()
    print(.y)
    load(paste0(dir,.x),envir = ne)
    zz = ne$enrich.matrix.pval[,DMG_bins]
    zz1 = apply(zz,1,function(v) {
      -log10(metap::sumlog(v)$p)
    })
    zz1[zz1 < -log10(0.05)] = 0
    out = data.frame(zz1) %>% set_colnames(.y)
  })  
DMGS.SCZ.result = as.data.frame(DMGS.SCZ.result["V80",])

LOF.result = list(LOF = LOF.file)  %>%
  imap_dfc(~{
    ne = new.env()
    print(.y)
    load(paste0(dir,.x),envir = ne)
    zz = ne$enrich.matrix.pval[,LOF_bins]
    zz1 = -log10(zz)
    zz1[zz1 < -log10(0.05)] = 0
    out = data.frame(zz1) %>% set_colnames(.y)
  }) 
LOF.result = as.data.frame(LOF.result["V80",])

other.results = cbind(DEGS.CAUDATE.result,DEGS.DLPFC.result,DEGS.HIPPO.result,DMGS.SCZ.result,LOF.result)
colnames(other.results) = c("DEGs CN", "DEGs DLPFC", "DEGs HP", "DMGs", "LoF")

###
MAGMA.result = MAGMA.files %>% set_names(strsplit2(patho.files,"\\.LIBD")[,1]) %>%
  imap_dfc(~{
    ne = new.env()
    print(.y)
    load(paste0(dir,.x),envir = ne)
    if(length(grep("SCZ",.y)) == 0) {zz = ne$PGC_MAGMA_out$all.biotypes$`100kbp`[[1]]$P_fdr
    } else zz = ne$PGC_MAGMA_out$all.biotypes$`100kbp`[[2]]$P_fdr
    zz1 = -log10(zz)
    zz1[zz1 < -log10(0.05)] = 0
    zz1
    # sig.wins = data.frame(rowSums(zz<0.05)) %>% set_colnames(.y)
  }) %>% set_colnames(gsub(".all.biotypes","",gsub(".protein.coding","",colnames(.)))) %>% set_rownames(.,comps) 
MAGMA.result = as.data.frame(MAGMA.result["V80",])

###
HMAGMA.result = HMAGMA.files %>% set_names(strsplit2(patho.files,"\\.LIBD")[,1]) %>%
  imap_dfc(~{
    ne = new.env()
    print(.y)
    load(paste0(dir,.x),envir = ne)
    if(length(grep("SCZ",.y)) == 0) {zz = ne$PGC_MAGMA_out$all.biotypes$Adult_brain[[1]]$P_fdr
    } else zz = ne$PGC_MAGMA_out$all.biotypes$Adult_brain[[2]]$P_fdr
    zz1 = -log10(zz)
    zz1[zz1 < -log10(0.05)] = 0
    zz1
    # sig.wins = data.frame(rowSums(zz<0.05)) %>% set_colnames(.y)
  }) %>% set_colnames(gsub(".all.biotypes","",gsub(".protein.coding","",colnames(.)))) 
HMAGMA.result = as.data.frame(HMAGMA.result["V80",])

colnames(MAGMA.result)[colnames(MAGMA.result) == "BIP"] = "BD"
colnames(HMAGMA.result)[colnames(HMAGMA.result) == "BIP"] = "BD"
MAGMA.result = MAGMA.result[, c("SCZ","MDD","BD", "SA", "ASD","ADHD", "OCD","PTSD","CD","UC")]
HMAGMA.result = MAGMA.result[, c("SCZ","MDD","BD", "SA", "ASD","ADHD", "OCD","PTSD","CD","UC")]

#### tissue activity
load("sda_quantile/ALL/seed_400/tissue.score.0.4corr.RData")
tissue = apply(tissue.score.c,2,as.numeric)
row.names(tissue) = row.names(tissue.score.c)
colnames(tissue) = c("CN","DLPFC","HP")
tissue = as.data.frame(tissue[row.names(patho.results),])
# tissue = as.data.frame(t(tissue["V80",]))

# row.names(patho.results) = row.names(other.results) = row.names(MAGMA.result) = row.names(HMAGMA.result) = 
#   row.names(tissue) = "C80"
row.names(patho.results) = row.names(other.results) = row.names(MAGMA.result) = row.names(HMAGMA.result) =
  row.names(tissue) = gsub("V","C",row.names(patho.results))
  
#### Visualisation for Patho, DEGs, DMGs, LOF and Dx
# scale_patho = colorRamp2(c(0, 3, 6, 9), c("beige", "orange", "red"))   #Scale for patho enrichments
scale_patho = colorRamp2(c(0, 2, 4), c("snow", "orange", "red"))   #Scale for patho enrichments
scale_other = colorRamp2(c(0, 2, 4), c("snow", "mediumseagreen", "darkgreen"))   #Scale for DEG, DMG, LOF and Diagnosis enrichments
scale_tissue = colorRamp2(c(0, 1), c("snow", "lightsteelblue"))   #Scale for DEG, DMG, LOF and Diagnosis enrichments

ht0 = Heatmap(patho.results,
              na_col = "grey", name = "ht0", col = scale_patho, cluster_rows = F, cluster_columns = F,
              row_title    = NULL ,row_title_gp    = gpar(fontsize = 0, fontface = "bold"), row_title_side    = "left", row_title_rot    = 90, #row_labels = SCZ_risk_modules_df$new.network,
              column_title = "GWAS enrichemnt" ,column_title_gp = gpar(fontsize = 0, fontface = "italic"), column_title_side = "top" , column_title_rot = 0,
              
              row_names_side    = "left", row_names_rot    = 0 , row_names_gp    = gpar(fontsize = 0, fontface = "bold"   ),
              column_names_side = "bottom" , column_names_rot = 45, column_names_gp = gpar(fontsize = 0, fontface = 2, lty = 1),
              rect_gp = gpar(col = "black", lty = 1, lwd = 0.5),
              show_heatmap_legend = F, heatmap_legend_param = list(title = "-log10(adjusted p-value)",title_gp = gpar(fontsize = 18), direction = "horizontal", at = c(0,2,4), labels = c(0,2,4), legend_width = unit(7, "lines"), title_position ="topcenter")
              )

ht1 = Heatmap(other.results,
              na_col = "grey", name = "ht1", col = scale_other, cluster_rows = F, cluster_columns = F,
              row_title    = NULL ,row_title_gp    = gpar(fontsize = 0, fontface = "bold"), row_title_side    = "left", row_title_rot    = 90, #row_labels = SCZ_risk_modules_df$new.network,
              column_title = "other enrichment" ,column_title_gp = gpar(fontsize = 0, fontface = "italic"), column_title_side = "top" , column_title_rot = 0,
              
              row_names_side    = "left", row_names_rot    = 0 , row_names_gp    = gpar(fontsize = 0, fontface = "bold"     ),
              column_names_side = "bottom" , column_names_rot = 45, column_names_gp = gpar(fontsize = 25, fontface = 2, lty = 1),
              rect_gp = gpar(col = "black", lty = 1, lwd = 0.5),
              show_heatmap_legend = F, heatmap_legend_param = list(title = "-log10(adjusted p-value)",title_gp = gpar(fontsize = 18,fontfamily = "sans"), direction = "horizontal", at = c(0,2,4), labels = c(0,2,4), legend_width = unit(7, "lines"), title_position ="topcenter")
)

ht2 = Heatmap(MAGMA.result, 
              na_col = "grey", name = "ht2", col = scale_patho, cluster_rows = F, cluster_columns = F,
              row_title    = NULL ,row_title_gp    = gpar(fontsize = 0, fontface = "bold"), row_title_side    = "left", row_title_rot    = 90, #row_labels = SCZ_risk_modules_df$new.network,
              column_title = "MAGMA enrichemnt" ,column_title_gp = gpar(fontsize = 0, fontface = "italic"), column_title_side = "top" , column_title_rot = 0,
              
              row_names_side    = "left", row_names_rot    = 0 , row_names_gp    = gpar(fontsize = 0, fontface = "bold"     ),
              column_names_side = "bottom" , column_names_rot = 45, column_names_gp = gpar(fontsize = 0, fontface = 2, lty = 1),
              rect_gp = gpar(col = "black", lty = 1, lwd = 0.5),
              show_heatmap_legend = F, heatmap_legend_param = list(title = "-log10(adjusted p-value)", title_gp = gpar(fontsize = 8), direction = "horizontal", at = c(0,2,4), labels = c(0,2,4), legend_width = unit(7, "lines"), title_position ="topcenter")
)

ht3 = Heatmap(HMAGMA.result,
              na_col = "grey", name = "ht3", col = scale_patho, cluster_rows = F, cluster_columns = F,
              row_title    = NULL ,row_title_gp    = gpar(fontsize = 0, fontface = "bold"), row_title_side    = "left", row_title_rot    = 90, #row_labels = SCZ_risk_modules_df$new.network,
              column_title = "H-MAGMA enrichment" ,column_title_gp = gpar(fontsize = 0, fontface = "italic"), column_title_side = "top" , column_title_rot = 0,

              row_names_side    = "left", row_names_rot    = 0 , row_names_gp    = gpar(fontsize = 0, fontface = "bold"     ),
              column_names_side = "bottom" , column_names_rot = 45, column_names_gp = gpar(fontsize = 0, fontface = 2, lty = 1),
              rect_gp = gpar(col = "black", lty = 1, lwd = 0.5),
              show_heatmap_legend = F, heatmap_legend_param = list(title = "-log10(adjusted p-value)", 
                                                                   title_gp = gpar(fontsize = 8), direction = "horizontal", at = c(0,2,4), labels = c(0,2,4), legend_width = unit(7, "lines"), title_position ="topcenter")
)

ht4 = Heatmap(tissue,
              na_col = "grey", name = "ht4", col = scale_tissue, cluster_rows = F, cluster_columns = F,
              row_title    = NULL ,row_title_gp    = gpar(fontsize = 0, fontface = "bold"), row_title_side    = "left", row_title_rot    = 90, #row_labels = SCZ_risk_modules_df$new.network,
              column_title = "tissue specificity" ,column_title_gp = gpar(fontsize = 0, fontface = "italic"), column_title_side = "top" , column_title_rot = 0,

              row_names_side    = "left", row_names_rot = 0 , row_names_gp    = gpar(fontsize = 0, fontface = "bold"     ),
              column_names_side = "bottom" , column_names_rot = 45, column_names_gp = gpar(fontsize = 25, fontface = 2, lty = 1),
              rect_gp = gpar(col = "black", lty = 1, lwd = 0.5),
              show_heatmap_legend = F, heatmap_legend_param = list(title = "tissue specificty", title_gp = gpar(fontsize = 18), direction = "horizontal", at = c(0,1), labels = c(0,1), legend_width = unit(7, "lines"), title_position = "topcenter")
)

d = draw(ht0 + ht2 + ht3 + ht1 + ht4,
         column_title_gp = gpar(fontsize = 10, fontface = "bold"),
         auto_adjust = T, merge_legends = T,
         ht_gap = unit(c(15), "mm"),
         padding = unit(c(80, 20, 80, 20), "mm"), #bottom, left, top, right paddings)
         heatmap_legend_side = "top"
) 
```
