## Supplementary material for "Dopamine and schizophrenia from bench to bedside: Discovery of a striatal co-expression risk gene set that predicts in vivo measures of striatal function": Compressed zipped directory containing scripts used for this study.: C80_KCL_genetic.ancestries.R

```
##### load metanalaysis of NC and SCZ 
pop = c("ALL","noAFR.noEAS","CAUC","CAUC.ancestry")
striatum = c("whole_striatum","sensorimotor_striatum","limbic_striatum","associative_striatum")
df.final = purrr::map_dfr(pop, function(z){
  load(paste0("ENIGMA_GE/KCL.NC.SCZ/metanalysis.KCL.NC.SCZ_zscore_",z,".RData"))
  df = purrr::map_dfr(striatum,function(x){
    df1 = data.frame(as.data.frame(confint(df.meta$kbp100$PGC3.LIBD[[x]]$V80_PRS$PRS2,random = F, fixed = T, level = 1-(0.05/1))),
                     # lower = df.meta$kbp100$PGC3.LIBD[[x]]$V80_PRS$PRS1$ci.lb,
                     # upper = df.meta$kbp100$PGC3.LIBD[[x]]$V80_PRS$PRS1$ci.ub,
                     type = "C80 Parsed\n Polygenic Risk Score",
                     striatum = x,
                     pop = z)
    df2 = data.frame(as.data.frame(confint(df.meta$kbp100$PGC3.LIBD[[x]]$V80_compl$PRS2,random = F, fixed = T, level = 1-(0.05/1))),
                     # lower = df.meta$kbp100$PGC3.LIBD[[x]]$V80_compl$PRS1$ci.lb,
                     # upper = df.meta$kbp100$PGC3.LIBD[[x]]$V80_compl$PRS1$ci.ub,
                     type = "Complementary C80\n Parsed Polygenic Risk Score",
                     striatum = x,
                     pop = z)
    df = rbind(df1,df2)
  })
})
df.final$pop = factor(df.final$pop, levels = pop)
df.final$striatum = gsub("_striatum","",df.final$striatum)
df.final$striatum = factor(df.final$striatum, levels = gsub("_striatum","",striatum))

library(ggplot2)
library(RColorBrewer)
col = brewer.pal(4,"Pastel1")
p = ggplot(df.final, aes(x = pop, y = estimate, fill = pop)) + 
  geom_bar(stat = "identity", color = "black") + 
  geom_errorbar(aes(ymin = ci.lb, ymax = ci.ub), width = .2, position = position_dodge(.9)) +
  facet_grid(type~striatum) +
  scale_fill_manual(values = col, labels = c("ALL", "no AFR and no EAS", "EUR", "EUR ancestry score")) +
  xlab("") +
  ylab(expression(Fisher*"'"*s~z[r])) +
  theme_classic(base_size = 26) +
  theme(axis.text.x = element_blank(),
        legend.position = "bottom",
        strip.text.y = element_text(size = 16)) 
p  
ggsave(filename = paste0("sda_quantile/Paper/C80_KCL_different.ancestries.svg"),plot = p, width=10.5, height=7.6,dpi = 600)


  
  
```
