## Supplementary material for "Dopamine and schizophrenia from bench to bedside: Discovery of a striatal co-expression risk gene set that predicts in vivo measures of striatal function": Compressed zipped directory containing scripts used for this study.: cell.specificity_plot.R

```
require(reshape2)
library(patchwork)
library(ggplot2)

#### cell specificity
# load("sda_quantile/ALL/seed_400/PIP_0.5/output/Cell_specificity/GSEA_LIBD_ALL_CAUD.DLPFC.HIPPO_filtered_0.4corr_allKI_mouse.RData")
load("GTEx/v8/seed_400/PIP_0.5/Cell_specificity/GSEA_GTEx_DLPFC.CN.HP_allKI_mouse.RData")
mouse.cell = results.log10
mouse.cell = as.data.frame(t(mouse.cell["V18",]))
colnames(mouse.cell)[1] = gsub("_","/",colnames(mouse.cell)[1])
# load("sda_quantile/ALL/seed_400/PIP_0.5/output/Cell_specificity/GSEA_LIBD_ALL_CAUD.DLPFC.HIPPO_filtered_0.4corr_DRONC_human.RData")
load("GTEx/v8/seed_400/PIP_0.5/Cell_specificity/GSEA_GTEx_DLPFC.CN.HP_DRONC_human.RData")
human.cell = results.log10
human.cell = as.data.frame(t(human.cell["V18",]))

human.df = melt(human.cell)
human.df$type = "human"
mouse.df = melt(mouse.cell)
mouse.df$type = "mouse"
df = rbind(human.df,mouse.df)

human <- ggplot(human.df, aes(x = variable, y = value)) +
  geom_bar(stat = "identity") +
  geom_hline(yintercept = -log10(0.05), linetype="dashed", color = "red") +
  # geom_text(y = 10, x = 9, label = "-log10(.05)", color = "red", size = 6) +
  facet_wrap(.~type) +
  ylim(0,max(df$value)) +
  ylab("-log10(adjusted p-value)") + xlab("") +
  theme_classic(base_size = 20) +
  theme(axis.text.x = element_text(angle = 45,vjust = 0.5),
        # plot.margin = unit(c(1,1,1,1),"cm"),
        strip.placement = "inside")

mouse <- ggplot(mouse.df, aes(x = variable, y = value)) +
  geom_bar(stat = "identity") +
  geom_hline(yintercept = -log10(0.05), linetype="dashed", color = "red") +
  # geom_text(y = 3, x = 5, label = "-log10(.05)",color = "red") +
  facet_wrap(.~type) +
  ylim(0,max(df$value)) +
  ylab("") + xlab("") +
  theme_classic(base_size = 20) +
  theme(axis.text.x = element_text(angle = 45,hjust = 1),
        strip.placement = "inside")

my_graph = human + mouse + plot_layout(widths = c(5,9))
# my_graph
ggsave(filename = "sda_quantile/Paper/C18_cell.specificty.svg",plot = my_graph, width=12, height=7,dpi = 700)
```
