## Supplementary material for "Dopamine and schizophrenia from bench to bedside: Discovery of a striatal co-expression risk gene set that predicts in vivo measures of striatal function": Compressed zipped directory containing scripts used for this study.: code_for_analysis_plot.R

```
library(ggplot2)

df = read.table("sda_quantile/Paper/plot_mbx.txt", header = T) ## standardized values
df = cbind(df,LIBD_MBX3_C18_PRS)
summary(lm("BOLD~kbp100_C18_PRS1",data = df))
summary(lm("BOLD~kbp100_C18_compl.PGC3.LIBD_PRS1",data = df))

# df = read.table("sda_quantile/Paper/plot_uniba.txt", header = T) ## standardized values
my_graph <- ggplot(df, aes(x = kbp100_C18_compl.PGC3.LIBD_PRS1, y = BOLD)) +
  geom_point(size = 6,alpha = 0.8, 
             color = "black") +
  stat_smooth(method = "lm",
              se = T,
              linewidth = 1,
              color = "black",
              fullrange = T) +
  ylim(-3,3) + xlim(-3,3) +
  xlab("Complementary C18\nParsed Polygenic Risk Score") +
  ylab(paste0("Reward Anticipation\n","BOLD Response")) +
  theme_classic(base_size = 30)

my_graph
ggsave(filename = "sda_quantile/Paper/C80_MBX.svg",plot = my_graph, width=9.5, height=7,dpi = 600)
# ggsave(filename = "sda_quantile/Paper/C80_UNIBA.svg",plot = my_graph, width=9.5, height=7,dpi = 600)
```
