## Supplementary material for "Dopamine and schizophrenia from bench to bedside: Discovery of a striatal co-expression risk gene set that predicts in vivo measures of striatal function": Compressed zipped directory containing scripts used for this study.: code_to_plot.KCL.R

```
library(ggplot2)
library(metafor)
##### load NC or SCZ data
load("sda_quantile/Paper/plot_kcl.NC.RData")
df.NC = df
df.NC$Diagnosis = "NC"
load("sda_quantile/Paper/plot_kcl.SCZ.RData")
df.SCZ = df
df.SCZ$Diagnosis = "SCZ"
df = rbind(df.NC,df.SCZ)

my_graph <- ggplot(df, aes(x = varX, y = varY,color = Diagnosis)) +
  geom_point(size = 6,alpha = 0.8) +
  stat_smooth(method = "lm",
              se = T,
              linewidth = 1,
              fullrange = T) +
  scale_color_manual(values = c("black", "grey")) +
  ylim(-3,3) + xlim(-3,3) +
  xlab("C80 Parsed Polygenic Risk Score") +
  ylab("Presynaptic Dopamine\nSynthesis Capacity (Ki)") +
  theme_classic(base_size = 30) +
  theme(legend.position = c(0.15,0.86))

my_graph
ggsave(filename = "sda_quantile/Paper/C80_KCL.svg",plot = my_graph, width=9.5, height=6,dpi = 600)

##### load metanalaysis of NC and SCZ 
load("sda_quantile/Paper/plot_meta.kcl.NC.SCZ.RData")
pdf(file = paste0("sda_quantile/Paper/C80_KCL_meta.PRS1.whole.pdf"),width=10.5, height=6)
forest(meta,
           header = c("Diagnosis"),annotate = T,slab = c("NC","SCZ"), 
           cex = 2.5, cex.lab = 2,cex.axis = 2.5, level = 99.5
)
dev.off()
# title(paste0("C80 Stratified Polygenic Risk Score association with whole striatum Ki"), cex.main = 1.2,font.main =2)
```
