## Supplementary material for "Dopamine and schizophrenia from bench to bedside: Discovery of a striatal co-expression risk gene set that predicts in vivo measures of striatal function": Compressed zipped directory containing scripts used for this study.: code_to_plot.KCL_compl.R

```
library(ggplot2)
library(metafor)
##### load NC data
load("/Users/leonardo/Library/CloudStorage/OneDrive-UniversitàdegliStudidiBari/ENIGMA_GE/KCL.NC.SCZ/metanalysis.KCL.NC.SCZ_CAUC.ancestry.score.RData")
df.NC.PRS2.whole = df.transpose$kbp100$PGC3.LIBD$whole_striatum$V80_PRS$PRS2$NC$model
df.NC.PRS1.associative = df.transpose$kbp100$PGC3.LIBD$associative_striatum$V80_PRS$PRS1$NC$model
df.NC.PRS2.associative = df.transpose$kbp100$PGC3.LIBD$associative_striatum$V80_PRS$PRS2$NC$model

df.NC.PRS1.whole.compl = df.transpose$kbp100$PGC3.LIBD$whole_striatum$V80_compl$PRS1$NC$model
df.NC.PRS2.whole.compl = df.transpose$kbp100$PGC3.LIBD$whole_striatum$V80_compl$PRS2$NC$model
df.NC.PRS1.associative.compl = df.transpose$kbp100$PGC3.LIBD$associative_striatum$V80_compl$PRS1$NC$model
df.NC.PRS2.associative.compl = df.transpose$kbp100$PGC3.LIBD$associative_striatum$V80_compl$PRS2$NC$model

df.NC = sapply(grep("df\\.NC",ls(),value = T),function(x) get(x),simplify = F)
df.NC = lapply(df.NC,function(x){
  mod = paste0(colnames(x)[1],"~",paste0(colnames(x)[3:ncol(x)],collapse = "+"))
  x$varY = scale(lm(as.formula(mod),data = x)$residuals)
  x$varX = scale(x[[2]])
  x$Diagnosis = "NC"
  x[,c("varY","varX","Diagnosis")]
})

##### load SCZ data
df.SCZ.PRS2.whole = df.transpose$kbp100$PGC3.LIBD$whole_striatum$V80_PRS$PRS2$SCZ$model
df.SCZ.PRS1.associative = df.transpose$kbp100$PGC3.LIBD$associative_striatum$V80_PRS$PRS1$SCZ$model
df.SCZ.PRS2.associative = df.transpose$kbp100$PGC3.LIBD$associative_striatum$V80_PRS$PRS2$SCZ$model

df.SCZ.PRS1.whole.compl = df.transpose$kbp100$PGC3.LIBD$whole_striatum$V80_compl$PRS1$SCZ$model
df.SCZ.PRS2.whole.compl = df.transpose$kbp100$PGC3.LIBD$whole_striatum$V80_compl$PRS2$SCZ$model
df.SCZ.PRS1.associative.compl = df.transpose$kbp100$PGC3.LIBD$associative_striatum$V80_compl$PRS1$SCZ$model
df.SCZ.PRS2.associative.compl = df.transpose$kbp100$PGC3.LIBD$associative_striatum$V80_compl$PRS2$SCZ$model

df.SCZ = sapply(grep("df\\.SCZ",ls(),value = T),function(x) get(x),simplify = F)
df.SCZ = lapply(df.SCZ,function(x){
  mod = paste0(colnames(x)[1],"~",paste0(colnames(x)[3:ncol(x)],collapse = "+"))
  x$varY = scale(lm(as.formula(mod),data = x)$residuals)
  x$varX = scale(x[[2]])
  x$Diagnosis = "SCZ"
  x[,c("varY","varX","Diagnosis")]
})

names(df.NC) = names(df.SCZ) = gsub("df\\.NC\\.","",names(df.NC))
df = purrr::map2(df.NC,df.SCZ,rbind)

purrr::iwalk(df,~{
  if(ncol(limma::strsplit2(.y,"\\.")) == 3 ){ ylab = ""; xlab = "Complementary C80\n Parsed Polygenic Risk Score"
  } else {ylab = "Presynaptic Dopamine\nSynthesis Capacity (Ki)"; xlab = "C80 Parsed Polygenic Risk Score"}
  my_graph <- ggplot(.x, aes(x = varX, y = varY,color = Diagnosis)) +
    geom_point(size = 6,alpha = 0.8) +
    stat_smooth(method = "lm",
                se = T,
                linewidth = 1,
                fullrange = T) +
    scale_color_manual(values = c("black", "grey")) +
    ylim(-3,3) + xlim(-3,3) +
    xlab(xlab) +
    ylab(ylab) +
    theme_classic(base_size = 30) +
    theme(legend.position = c(0.15,0.86))
  
  ggsave(filename = paste0("sda_quantile/Paper/C80_KCL_",.y,".svg"),plot = my_graph, width=10.5, height=6,dpi = 600)
})


##### load metanalaysis of NC and SCZ 
load("/Users/leonardo/Library/CloudStorage/OneDrive-UniversitàdegliStudidiBari/ENIGMA_GE/KCL.NC.SCZ/metanalysis.KCL.NC.SCZ_zscore.RData")
meta.PRS2.whole = df.meta$kbp100$PGC3.LIBD$whole_striatum$V80_PRS$PRS2
meta.PRS1.associative = df.meta$kbp100$PGC3.LIBD$associative_striatum$V80_PRS$PRS1
meta.PRS2.associative = df.meta$kbp100$PGC3.LIBD$associative_striatum$V80_PRS$PRS2

meta.PRS1.whole.compl = df.meta$kbp100$PGC3.LIBD$whole_striatum$V80_compl$PRS1
meta.PRS2.whole.compl = df.meta$kbp100$PGC3.LIBD$whole_striatum$V80_compl$PRS2
meta.PRS1.associative.compl = df.meta$kbp100$PGC3.LIBD$associative_striatum$V80_compl$PRS1
meta.PRS2.associative.compl = df.meta$kbp100$PGC3.LIBD$associative_striatum$V80_compl$PRS2

df.meta = sapply(grep("^meta",ls(),value = T),function(x) get(x),simplify = F)


purrr::iwalk(df.meta,~{
  pdf(file = paste0("sda_quantile/Paper/C80_KCL_",.y,".pdf"),width=10.5, height=6)
  forest(.x,
         header = c("Diagnosis"),annotate = T,slab = c("NC","SCZ"), 
         cex = 2.5, cex.lab = 2,cex.axis = 2.5, level = 99.5
  )
  dev.off()
})
```
