## Supplementary material for "Dopamine and schizophrenia from bench to bedside: Discovery of a striatal co-expression risk gene set that predicts in vivo measures of striatal function": Compressed zipped directory containing scripts used for this study.: Dx.PRS_plot.R

```
library(ggplot2)
library(metafor)
library(patchwork)
##### load NC or SCZ data
load("sda_quantile/ALL/seed_400/data.SDA.ALL.04corr.RData")
df = data[data$Age >= 17,]
my_graph_C80 <- ggplot(df, aes(x = Dx, y = V80)) +
  geom_boxplot(fill = "darkgrey",notch = T,width = 0.5) +
  geom_jitter(width = 0.1,color = "black",alpha = 0.6, size = 3) +
  scale_x_discrete(labels = c("NC","SCZ")) +
  ylim(-1,1) +
  ylab("C80") + xlab("") +
  theme_classic(base_size = 26) 
  # theme(plot.margin = unit(c(2,8,2,2),"cm"))

my_graph_C109 <- ggplot(data = data[data$Age >= 17,], aes(x = Dx, y = V109)) +
  geom_boxplot(fill = "darkgrey",notch = T,width = 0.5) +
  geom_jitter(width = 0.1,color = "black",alpha = 0.6, size = 3) +
  scale_x_discrete(labels = c("NC","SCZ")) +
  ylim(-1,1) +
  ylab("C109") + xlab("") +
  theme_classic(base_size = 26) 
  # theme(plot.margin = unit(c(2,8,2,2),"cm"))

df = df[df$Race %in% "CAUC",]
df$PRS6.PGC2 = scale(df$PRS6.PGC2)
df$PRS6.PGC3 = scale(df$PRS6.PGC3)
my_PRS_C80 <- ggplot(df, aes(x = PRS6.PGC2, y = V80)) +
  geom_point(size = 4, 
             color = "black",alpha = 0.6) +
  stat_smooth(method = "lm",
              se = T,
              linewidth = 1,
              color = "black",
              fullrange = T) +
  ylim(-1,1) +
  xlim(-3,3) +
  xlab("Whole genome PRS") +
  ylab("") +
  theme_classic(base_size = 26) 
  # theme(plot.margin = unit(c(2,6,2,2),"cm"))

load("~/OneDrive - Università degli Studi di Bari/sda_quantile/Heatmap_data/LM_detect.final.RData")
df = PRS_assoc$V109$PRS6.PGC3$model
df$PRS6.PGC3 = scale(df$PRS6.PGC3)
my_PRS_C109 <- ggplot(df, aes(x = PRS6.PGC3, y = V109)) +
  geom_point(size = 4, 
             color = "black",alpha = 0.6) +
  stat_smooth(method = "lm",
              se = T,
              linewidth = 1,
              color = "black",
              fullrange = T) +
  ylim(-1,1) +
  xlim(-3,3) +
  xlab("Whole genome PRS") +
  ylab("") +
  theme_classic(base_size = 26) 
  # theme(plot.margin = unit(c(2,6,2,2),"cm"))

p = my_graph_C80 + my_PRS_C80 + plot_layout(widths = c(6, 9))
p2 = my_graph_C109 + my_PRS_C109 + plot_layout(widths = c(6,9))

p
p2
ggsave(filename = "sda_quantile/Paper/C80.svg",plot = p, width=8.5, height=5,dpi = 600)
ggsave(filename = "sda_quantile/Paper/C109.svg",plot = p2, width=8.5, height=5,dpi = 600)
```
