## Supplementary material for "Dopamine and schizophrenia from bench to bedside: Discovery of a striatal co-expression risk gene set that predicts in vivo measures of striatal function": Compressed zipped directory containing scripts used for this study.: GO_plot.R

```
library(patchwork)
library(ggplot2)
library(ggplotify)

BP = ll.GO$BP$V80
MF = ll.GO$MF$V80
CC = ll.GO$CC$V80
KEGG = ll.KEGG$KEGG$V80
Reactome = ll.Reactome$Reactome$V80
rm(list=setdiff(ls(),c("BP","MF","CC","KEGG","Reactome")))
Annotations = sapply(ls(),get,USE.NAMES = T,simplify = F)
Annotations = lapply(Annotations, function(x){
  x$BgRatio = as.numeric(limma::strsplit2(x$BgRatio,'/')[,1])/as.numeric(limma::strsplit2(x$BgRatio,'/')[,2])
  x$fold = x$GeneRatio/x$BgRatio
  x
})
Annotations = lapply(Annotations, function(x) {x[order(x$p.adjust),]; x})
Annotations = lapply(Annotations, function(x) {x = x[c(1:10),]; x$Description = as.character(x$Description);x})

Annotations.plot = lapply(Annotations,function(x){
  x$Description = factor(x$Description, levels = rev(x$Description))
  p = ggplot(x, aes(x= fold, y=Description, color = p.adjust,fill = p.adjust)) +
    geom_point(size = 5,shape = 23) + xlab("Fold enrichment") + ylab("") +
    scale_color_continuous(low = "red", high = "blue", name = x$p.adjust, guide= guide_colorbar(title = "adjusted p-value", reverse = T)) +
    scale_fill_continuous(low = "red", high = "blue", name = x$p.adjust, guide = guide_none()) +
    ggtitle("") + theme_linedraw(base_size = 25) +  
    theme(legend.text = element_text(size = 16))
})

p = p = Annotations.plot$CC + theme(legend.position = "right") + scale_y_discrete(position = "right")
p
ggsave(filename = "sda_quantile/Paper/C18_CC.svg",plot = p, width=15, height=7,dpi = 600)

# df = Annotations$CC
# ggplot(df, aes(x= GeneRatio, y=Description, size = Count, color = p.adjust)) +
#   geom_point() + xlab("GeneRatio") +
#   scale_color_continuous(low = "red", high = "blue", name = df$p.adjust, guide= guide_colorbar(title = "p.adjust", reverse = T)) +
#   ggtitle("") + theme_dose(12) + scale_size(range = c(3, 8), guide = guide_legend(title = "Count"))  +
#   theme(axis.text.y = element_text(size = 10))  
```
