## Supplementary material for "Dopamine and schizophrenia from bench to bedside: Discovery of a striatal co-expression risk gene set that predicts in vivo measures of striatal function": Compressed zipped directory containing scripts used for this study.: KCL_PCs.R

```
library(tidyverse)
sample = c("KCL.NC.SCZ")
eigenval <- scan(paste0("ENIGMA_GE/PCA/HM3.",sample,".mds.mds.eigvals"))
pve <- data.frame(PC = 1:length(eigenval), pve = eigenval/sum(eigenval)*100)
mds.cluster <- read.csv(paste0("ENIGMA_GE/PCA/HM3.",sample,".mds.mds"), sep="")
mds.cluster = mds.cluster[,c(1:3,grep("C1$|C2$",colnames(mds.cluster)))]
populations = c("CEU",
                "CHB",
                "YRI",
                "TSI",
                "JPT",
                "CHD",
                "MEX",
                "GIH",
                "ASW","LWK",
                "MKK")
super.populations = c("AFR","EAS","EUR","SAS","AMR")
mds.cluster$Ethnicity = factor(sapply(mds.cluster$FID,function(x) {
  
  if(x %in% populations){ populations[populations == x]
    
  } else sample
  
  
}),levels = c(sample,populations))
mds.cluster$populations = factor(sapply(mds.cluster$FID,function(x) {
  
  if(x %in% c("YRI","LWK","ASW","MKK")){ super.populations[1]
  } else if(x %in% c("CHB","JPT","CHD")){ super.populations[2]
  } else if(x %in% c("CEU","TSI")){ super.populations[3]
  } else if(x %in% c("GIH")){ super.populations[4]
  } else if(x %in% c("MEX")){ super.populations[5]
  } else sample
  
  
}),levels = c(sample,super.populations))
mds.cluster$alpha = sapply(mds.cluster$Ethnicity,function(x) {
  
  if(x %in% sample) 1 else 0.5 
  
})
mds.cluster$populations = as.character(mds.cluster$populations)
mds.cluster$populations = gsub("KCL.NC.SCZ","KCL",mds.cluster$populations)
mds.cluster$populations = factor(mds.cluster$populations, levels = c("KCL","AFR","AMR","EAS","EUR","SAS"))
df = data.frame(x = c(0))
p = ggplot(mds.cluster, aes(C1, C2, color = populations)) + 
  geom_point(size = 4,alpha = mds.cluster$alpha) + 
  xlab(paste0("PC1")) +
  ylab(paste0("PC2")) +
  scale_color_brewer(palette = "Set1") +
  geom_segment(x = -0.15, y = 0.013, xend = 0.047, yend = 0.013, linetype = "dashed",color = "black") +
  geom_segment(x = 0.047, y = 0.013, xend = 0.047, yend = -1, linetype = "dashed",color = "black") +
  geom_segment(x = -0.15, y = -0.049, xend = 0, yend = -0.049, linetype = "dashed",color = "black") +
  geom_segment(x = 0, y = -0.049, xend = 0, yend = -1, linetype = "dashed",color = "black") +
  theme_classic(base_size = 26) +
  theme(legend.position = "right") 
p

ggsave(filename = paste0("sda_quantile/Paper/KCL_PCs.svg"),plot = p, width=10.5, height=7.6,dpi = 600)
```
