## Supplementary material for "Dopamine and schizophrenia from bench to bedside: Discovery of a striatal co-expression risk gene set that predicts in vivo measures of striatal function": Compressed zipped directory containing scripts used for this study.: LIBD.GTEX_replication.R

```
require(ggplot2)

load(paste0("sda_quantile/REPLICATION/GTEX/jaccard_index_sequential_replication_empPvalue.04corr.RData"))
#### data process JI ####
JI.df = data.frame(p = as.numeric(results$null_p),
                   LIBD = sapply(names(results$pair_correlation),function(x)limma::strsplit2(x,"\\.")[,1]),
                   GTEx = results$pair_component,
                   metric = round(results$pair_correlation,3))
JI.df$type = "JI"
JI.df$LIBD = gsub("V","C",JI.df$LIBD)
JI.df$GTEx = gsub("V","C",JI.df$GTEx)
JI.df$metric_p = JI.df$metric + 0.02

load(paste0("sda_quantile/REPLICATION/GTEX/gene_loading_sequential_replication_empPvalue.04corr.RData"))
#### data process gene loading ####
loading.df = data.frame(p = as.numeric(results$null_p),
                        LIBD = sapply(names(results$pair_correlation),function(x)limma::strsplit2(x,"\\.")[,1]),
                        GTEx = results$pair_component,
                        metric = round(results$pair_correlation,3))
loading.df$metric_p = loading.df$metric + 0.02
loading.df$metric = loading.df$metric*-1
loading.df$metric_p = loading.df$metric_p*-1
loading.df$type = "Gene loading"
loading.df$LIBD = gsub("V","C",loading.df$LIBD)
loading.df$GTEx = gsub("V","C",loading.df$GTEx)

df = rbind(JI.df,loading.df)
df$LIBD = factor(df$LIBD,levels = rev(JI.df$LIBD))
df = df[order(df$LIBD),]
df$label_p = ""
df$label_p[df$p < .05] = "*"
df$label_p[df$p < .001] = "**"
df$GTEx_JI = df$GTEx_loading = df$GTEx
df$GTEx_JI[!df$type %in% "JI"] = ""
df$GTEx_loading[df$type %in% "JI"] = ""

##### plot
guide_axis_label_trans <- function(label_trans = identity, ...) {
  axis_guide <- guide_axis(...)
  axis_guide$label_trans <- rlang::as_function(label_trans)
  class(axis_guide) <- c("guide_axis_trans", class(axis_guide))
  axis_guide
}

guide_train.guide_axis_trans <- function(x, ...) {
  trained <- NextMethod()
  trained$key$.label <- x$label_trans(trained$key$.label)
  trained
}

p = ggplot(df, aes(x = LIBD, y = metric, fill = type)) +
  geom_bar(stat = "identity",width = 0.9) +
  coord_flip() +
  scale_fill_manual(values = alpha(c("dodgerblue","darkorange"),0.5)) +
  scale_y_continuous(breaks = c(-1, -0.75, -0.5,-0.25, 0, 0.25, 0.5, 0.75, 1),
                     labels = c("1", "0.75","0.5", "0.25", "0", "0.25", "0.5", "0.75","1"),
                     limits = c(-1,1)) +
  guides(y.sec = guide_axis_label_trans(~.)) +
  geom_text(y = df$metric_p, label = df$label_p, nudge_x = -0.25, size = 5) + 
  geom_text(y = 1, label = df$GTEx_JI, nudge_x = 0.1, nudge_y = 0.4, size = 6) +
  geom_text(y = -1, label = df$GTEx_loading, nudge_x = 0.1, nudge_y = -0.1, size = 6) +
  ylab("") + xlab("") +
  theme_classic(base_size = 30) +
  theme(legend.position = "none",
        axis.text.y = element_text(size = 20,face = "bold"),
        axis.ticks.y = element_blank())

ggsave(filename = "sda_quantile/Paper/LIBD.GTEX_replication.svg",plot = p, width=10, height=18,dpi = 600)
```
