## Supplementary material for "Dopamine and schizophrenia from bench to bedside: Discovery of a striatal co-expression risk gene set that predicts in vivo measures of striatal function": Compressed zipped directory containing scripts used for this study.: venn_plot.R

```
require(seqsetvis)
source("mir-137/Venn_diagram_ggplot.R")

###############
venn = list(JI = df.ji$names[df.ji$p > 3],
            'Gene loading' = df.gl$names[df.gl$p > 3])

plot = seqsetvis::ssvFeatureVenn(venn[-1],counts_txt_size = 0,
                                 line_width = 1.5,group_names = c("JI","Gene loading"),
                                 circle_colors = c("darkorange","dodgerblue")
)
p = plot + theme(legend.text = element_text(size = 20))
ggsave(filename = "sda_quantile/Paper/replication_venn.svg",plot = p, width=12, height=7,dpi = 600)


##############
length(intersect(V80_symbols$V1,
                 NAs_DRD1_DRD2_MSN$MSN.D2_A_1vAll))

to.remove = intersect(NAs_DRD1_DRD2_MSN$MSN.D1_A_1vAll,NAs_DRD1_DRD2_MSN$MSN.D2_A_1vAll)

D2_A = setdiff(NAs_DRD1_DRD2_MSN$MSN.D2_A_1vAll,to.remove)
D1_A = setdiff(NAs_DRD1_DRD2_MSN$MSN.D1_A_1vAll,to.remove)

venn = list(C80 = V80_symbols$V1,
            'D1 MSN' = NAs_DRD1_DRD2_MSN$MSN.D1_A_1vAll,
            'D2 MSN' = NAs_DRD1_DRD2_MSN$MSN.D2_A_1vAll)

plot = seqsetvis::ssvFeatureVenn(venn,
                                 line_width = 1.5,
                                 circle_colors = c("darkorange","dodgerblue","seagreen3")
)
p = plot + theme(legend.text = element_text(size = 20))
ggsave(filename = "sda_quantile/Paper/C80.D2.D1_venn.svg",plot = p, width=12, height=7,dpi = 600)

p

###############
venn = list(C80 = 1:34,
            C18 = c(1:16,35:51))

plot = seqsetvis::ssvFeatureVenn(venn,counts_txt_size = 0,
                                 line_width = 1.5,group_names = c("C80","C18"),
                                 circle_colors = c("darkorange","seagreen3")
)
p = plot + theme(legend.text = element_text(size = 20))
ggsave(filename = "sda_quantile/Paper/C80.C18_venn.svg",plot = p, width=12, height=7,dpi = 600)
```
